# The neural activity of auditory conscious perception

**DOI:** 10.1101/2023.01.12.523829

**Authors:** Kate L. Christison-Lagay, Noah C. Freedman, Christopher Micek, Aya Khalaf, Sharif I. Kronemer, Mariana M. Gusso, Lauren Kim, Sarit Forman, Julia Ding, Mark Aksen, Ahmad Abdel-Aty, Hunki Kwon, Noah Markowitz, Erin Yeagle, Elizabeth Espinal, Jose Herrero, Stephan Bickel, James Young, Ashesh Mehta, Kun Wu, Jason Gerrard, Eyiyemisi Damisah, Dennis Spencer, Hal Blumenfeld

## Abstract

Although recent work has made significant headway in understanding the temporal and spatial dynamics of the neural mechanisms of conscious perception, much of that work has focused on visual paradigms. To determine whether there are shared mechanisms for perceptual consciousness across sensory modalities, here we developed a task to test within the auditory domain. Participants (n=31) completed an auditory perceptual threshold task while undergoing intracranial electroencephalography (icEEG) for intractable epilepsy. Intracranial recordings from over 2,800 grey matter electrodes representing widespread cortical coverage were analyzed for power in the high gamma range (40–115 Hz)—a frequency range that reflects local neural activity. For trials that were perceived, we find activity in early auditory regions which is accompanied by activity in the right caudal middle frontal gyrus, and shortly thereafter by activity in non-auditory thalamus. This is followed by a wave of activity that sweeps through the higher auditory association regions and into parietal and frontal cortices, similar to the wave observed in our visual conscious perception paradigm. However, for not perceived trials, we find that significant activity is restricted to early auditory regions (and areas immediately adjacent to the Sylvian fissure). These findings show that the broad anatomical regions of cortical and subcortical networks involved in auditory perception are similar to the networks observed with vision, suggesting shared general mechanisms for conscious perception.

## 1. Introduction

The nature of consciousness has intrigued humankind for millennia, and is still an area of active inquiry across multiple disciplines including philosophy, psychology, and neuroscience. Although many of the questions surrounding the topic are still wide-open—what does it mean to be conscious? Are animals conscious? Could consciousness emerge from artificial intelligence?—there has been significant research over recent years dedicated to determining and describing the neural activity underlying conscious perception.

Sensory perception is often used for a proxy of consciousness, as it can be easily manipulated and interrogated by experimenters. Although decades of research have studied the neural mechanisms underlying consciousness and perception ^1^, the spatial and temporal dynamics of neural activity following consciously perceived sensory events remain incompletely understood. Previous findings from our laboratory and others have demonstrated that conscious perception is marked by activity in arousal and attention networks ^2–5^ and wide-spread activity that propagates from early sensory regions to higher order association regions of the cortex, as well as a downregulation of the default mode network ^4, 5^. Notably, the role of later, sustained activity in higher-order cortical regions is the subject of disagreement among consciousness researchers, and may be in part attributable to other task-related processes associated with perceptual report rather than conscious perception *itself.* Based on the literature and our previous findings in a visual perception task, we hypothesize that there are four stages that characterize conscious perception: (1) **detection** of stimuli in primary sensory regions (and perhaps other areas) (2) a transient neuromodulatory **pulse** in subcortical-cortical arousal and salience networks, (3) a **switch** on/off in sensory cortex, detection and task negative networks gating information flow, and (4) a sustained **wave** of cortical hierarchical processing and memory encoding for subsequent report ^6^. However, as this hypothesis is inspired almost exclusively by work within the visual domain, it remains unclear whether these mechanisms are shared across sensory systems.

Next to vision, audition comes in a distant second place in terms of best-studied conscious perception modalities. Moreover, much of the existing work on conscious auditory perception has been done on auditory scene analysis—that is, how the brain is able to detect, extract and then segregate and group the auditory environment into distinct percepts based on spatial, spectral and temporal regularities ^7–12^. Studies of auditory scene analysis have helped delineate the auditory processing streams and establish which cognitive processes occur at various levels of those streams ^7, 8, 12–19^. This literature is augmented by a significant number of studies specifically examining how language sounds are processed, as language processing involves several well-studied dedicated pathways and additional cognitive resources ^20–23^. However, these studies presuppose that conscious perception is occurring—they are generally carried out with fully audible stimuli, and often look at how the neural encoding of different percepts *differ:* for example, how one vs two auditory streams are differentially encoded ^13, 24–27^, or how a target evokes different activity than its standards in an oddball paradigm ^28–31^.

Although these studies are of critical importance in understanding auditory perception, it is also important to understand the underlying spatiotemporal dynamics that all these percepts share: that is, what happens when we consciously perceive *any* sound. Studies that focus on this aspect of conscious auditory perception—looking at the neural activity underlying whether a sound is consciously perceived or not— generally fall into two camps of paradigms: those that use near-threshold level sounds (presented either in silence, or embedded in a noisy background) and those which compare sounds presented with an auditory mask vs the same sound presented without a mask. There are remarkably few peri-threshold studies focused on poststimulus activity, but the existing studies have consistently shown that perceived sounds exhibit an early (~100 ms post stimulus) response associated with the auditory cortex and later (~300 ms) poststimulus component in more frontal regions ^32–35^. Furthermore, while unperceived trials may elicit a degree of response in the auditory cortex, there is a greater response if the stimulus is perceived ^36–39^. However, peri-threshold stimuli elicit a smaller neural response than suprathreshold stimuli, and for this reason, masking studies—in which perception of a suprathreshold target is manipulated by the presence (or absence) of additional sounds—provide an appealing alternative to near-threshold stimulus studies.

While masking studies are a powerful tool for studying conscious perception, they rely on the use of additional stimuli, which makes it impossible to fully disentangle differences in neural responses due to perception versus the physical differences in stimulus conditions. Here, we use a threshold-level stimulus in which physically identical stimuli were later reported by the participant as perceived or not. By using identical stimuli, any differences in neural activity we observe is due to processes endogenous to perceptual processing. We coupled this novel auditory paradigm with widespread recordings from a large population of participants undergoing intracranial encephalography. This technique provides unique access to highly spatially- and temporally-resolved signals in human participants. We focused our analysis on broadband gamma power (40– 115 Hz), which has been shown to be a proxy for local neuronal spiking activity ^40–42^. Here we show the first study (to our knowledge) providing whole-brain, highly spatiotemporally-resolved dynamics of the neural representation of auditory perception (or non-perception) in a non-masked auditory perceptual task.

## 2. Methods

### 2.1. Participants

All research procedures were approved by the local institutional review boards (IRB) at Yale University, The MountSinai Hospital and Northwell Health. A total of 38 adult patients undergoing craniotomy with intracranial electrodes for seizure monitoring participated in the study. 33 participants (including two repeat participants who underwent multiple surgeries to implant intracranial electrodes; females=16; mean age=38.8, range 19–69 years old) were recruited from the Yale Comprehensive Epilepsy Program; 4 additional participants (including one repeat participant who underwent multiple surgeries to implant intracranial electrodes) were recruited from Northwell Health, and 1 participant was recruited from Mount Sinai. Participants were recruited following approval from their medical team and participated following the participants’ written informed consent. Seven participants were excluded from analysis due to inadequate data collection (< 150 trials or inability to complete the task) or excess noise in the recordings; the remaining 31 patients were included in the analyses.

The implanted intracranial electrodes included subdural grids, strips, and depth electrodes. Electrode type, number, and placement were determined by the clinical team overseeing each case. Each participant was implanted with an average of 187 electrodes (range: 80–368) yielding a total of 5793 intracranial electrode contacts across all subjects. Visual inspection of co-registered structural MRI and whole-brain computed tomography (CT) images showed that of those 5793 electrodes, 2805 were in grey matter and were distributed bilaterally across cortical surface and depth sites (see Figure 2A). Analyses were carried out only for electrodes localized in grey matter.

### 2.2 Auditory paradigm

The behavioral task tested conscious auditory perception of target auditory stimuli embedded in a white noise background (produced by MATLAB function randn). Target stimuli were a waterdrop, a whistle, and a ‘laser’. The waterdrop and whistle were sourced from freesound.org; the laser sound effect was produced in Audacity® (https://audacityteam.org/) by stacking linearly interpolated downward octave slides ranging from 50 Hz–12.8 kHz. Sounds were passed through high (50 Hz, 6 dB per octave roll off) and low-pass filters (12 kHz, 48dB per octave roll off) to attenuate frequencies outside of that range, and were clipped to 75 ms. Sounds were selected because within lab testing showed they could be distinguished from each other and identified using only 75 ms of auditory stimuli. All initial sound editing was done using Audacity®. Target sounds were imported into MATLAB (The Mathworks Inc., Natick MA, United States). The amplitude of each target stimulus’ soundwave was scaled within MATLAB on a run-to-run basis to approximate the participant’s perceptual threshold (see below) and embedded into the MATLAB-generated white noise background.

The behavioral task was run on a MacBook Pro (13” Macbook Pro, 2015 with High Sierra OS), and sounds were presented through the laptop’s built-in speakers. Concurrent with the auditory stimuli, participants were asked to fixate on a central white fixation cross that appeared against a visual noise background on the laptop’s screen. For consistency across auditory and a previous visual task ^4^, the screen was placed 85 cm from the participant’s nose bridge. Extraneous stimuli were minimized by closing the window shades, turning off the lights and TV, and closing the door. Behavioral responses were recorded through button presses on a 4-button response box (Current Designs, Inc.) that was placed under the participant’s right hand.

Participants first underwent a calibration run to determine 50% perception target intensities. Specifically, auditory target intensities were presented at varying intensities over a white noise background with randomly jittered interstimulus interval of 1–2 s. Eight intensities spanning from undetectable to fully audible were played; each of the three 75 ms duration sounds were presented (in random order) at each intensity (also in random order) five times, resulting a total of 120 stimuli presented during calibration. Participants were instructed to press their ‘yes’ button immediately following detection of a target sound (‘yes’ button was counterbalanced to be ‘1’ or ‘2’ across participants). Following the calibration, the results of each target sound were fitted to a sigmoidal cumulative Weibull distribution function to estimate each sound’s 50% detection threshold.

After calibration, participants completed testing runs each consisting of 50 trials. Trials began with a randomly jittered 3–5 s pre-stimulus interval, during which auditory white noise played concurrently with visual noise as participants fixated on a white fixation cross (Figure 1A). After the pre-stimulus interval, one of the target stimuli was embedded in the auditory noise at its previously calculated threshold intensity. The target stimulus was followed by a randomly selected 3, 4, or 5 s post-stimulus period, during which the auditory white noise, visual noise and fixation cross continued to be presented. Following the post-stimulus interval, two non-timed, forced-choice questions were displayed on the laptop screen regarding (1) whether the participant heard a target stimulus, and (2) the identity of the target stimulus (Figure 1A). The participant was instructed to answer the identification question truthfully if they heard a stimulus and to press a random button if they did not. They were also informed that some of the trials would be blank (no target sound), but were not informed that the blank rate was 14%. Following the button press for the second question, the next trial began immediately. Participants reported behavior using their right hand (as with calibration, the ‘yes’ button was ‘1’ or ‘2’, counterbalanced across participants). At the end of each run, the 50% threshold intensity for each target was recalculated based on the participant’s performance, by updating the Weibull distribution parameters. This was accomplished by appending the most recent run’s behavior into the previous calibration and any previous run’s data for the given target stimulus; because each stimulus’s intensity was kept constant during a given run, in effect this added a run’s worth of trials at a single threshold-approximate intensity for each sound to the previously collected data. The entire dataset across all presented intensities (calibration and all runs to date) were refitted to a sigmoidal cumulative Weibull distribution function to estimate an updated 50% detection threshold for each sound, and this threshold value was used for the next run.

**Figure 1.**
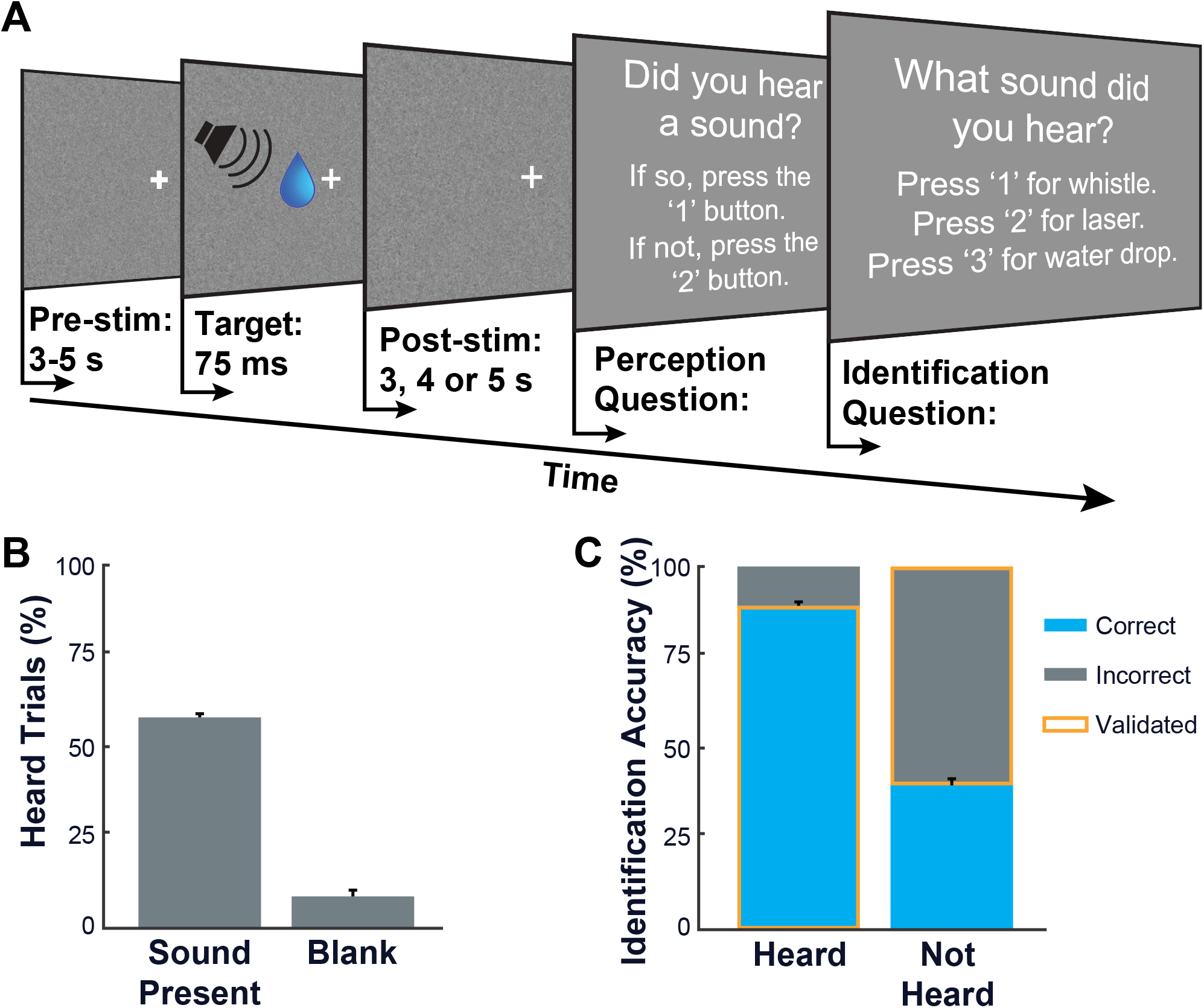
Auditory threshold task and behavior. **(A)** Single trial of auditory perceptual threshold task. A 3-5 second prestimulus interval was presented, consisting of auditory white noise played concurrently with visual noise with a central, white fixation cross displayed on a laptop. Following this, in 86% of trials, an auditory target stimulus (waterdrop, whistle or ‘laser’) was played at the participant’s previously-calibrated 50% detection threshold; in the remaining 14% of trials, no target stimulus was played (‘blanks’). Target (or blank) presentation was followed by a post-stimulus interval of 3, 4 or 5 seconds of visual and auditory noise. Following the post-stimulus interval, two non-timed, forced-choice questions were displayed on the laptop screen regarding (1) whether the participant heard a target stimulus (‘perception question’; shown here with ‘1’ for Yes and ‘2’ for No, but this was randomized by participant), and (2) the identity of the target stimulus (‘identification question’). Following button press for the second question, the next trial immediately began with a total of 50 trials per run. **(B)** Responses to perception question. In trials in which a target sound was present, 58.8% of trials were reported as heard; in trials without a target sound (‘blanks’), 8.5% of trials were reported as heard. Error bars indicate SEM. (**C**) Responses to identification question. For trials in which a target sound was presented, targets were correctly identified in 89.2% of trials. In contrast, participations correctly identified only 40.2% of the target sounds when they were reported as not heard. Outlined in yellow are the validated and confirmed “perceived” and “not perceived” trials analyzed in this study, heretofore referred to simply as “Perceived” and “Not Perceived.” N = 31 participants.

The task was implemented in MATLAB (Matlab 2018a, The Mathworks Inc., Natick MA, United States) using the Psychophysics Toolbox (‘Psychtoolbox’) extensions ^43–45^.

### 2.3. Behavioral analysis

Two sets of behavioral analyses were performed. First, on a participant basis, we analyzed the percentage of trials in which (1) a target sound was present and the participant indicated they heard the target and in which (2) no target sound was present, but the participant indicated they heard a target (Figure 1B). Second, for trials in which a target sound was presented, we analyzed how well participants were able to identify a target sound as a function of whether they said they heard it or not (Figure 1C). Trials in which the participant reported that they heard the target sound (question 1), and they correctly identified it (question 2) were considered to be validated and confirmed as *Perceived;* and trials in which the participant said they did not hear the sound (question 1), and they incorrectly identified the sound played (question 2) were considered to be confirmed as *Not Perceived.* Only the trials confirmed as Perceived or Not Perceived were considered for further analyses. For each participant, we excluded all data for a sound if the percentage of heard trials that were correctly identified was < 60%, and only trials for the remaining sounds were analyzed. If more than one target sound fell below this criterion, the participant was eliminated. These criteria were included to ensure that participants understood and were properly performing the task. One participant was eliminated because two targets fell below this threshold (the final sound was narrowly above the threshold); two additional participants had a single target sound eliminated from analyses (resulting in the exclusion of 28.5% and 29% of trials for these participants, accounting for .094% of overall trials across all participants). Overall, this criterion eliminated 2.9% of trials across all participants.

### 2.4. Intracranial EEG data acquisition

The intracranial EEG data collected at Yale New Haven Hospital epilepsy monitoring unit were recorded with a Natus Neurolink/Braintronics amplifier and relevant data segments were exported using Natus NeuroWorks software (www.neuro.natus.com)(sampling rate 4096 Hz, bandpass filter with corner frequencies of 0.1 Hz and 1.33 kHz; 1 patient sampled at 1024 Hz, bandpass filter with corner frequencies of 0.1 Hz and 400 Hz). Recordings at Northwell were conducted with Tucker-Davis Technologies using a PZ5M amplifier (sampling rate 3 kHz, bandpass filter with corner frequencies of 0.4 Hz and 1.5 kHz) and exported using Tucker-Davis software. Recordings at Mt. Sinai were conducted with Natus NeuroWorks (sampling rate 2048 Hz, with bandpass filter with corner frequencies of 0.1 Hz and 670 Hz). The clinical team selected the reference recording electrode for each participant that best reduced the visible EEG artifacts.

To ensure proper synchronization between task event times and the corresponding EEG recordings, transistor-transistor logic (TTL) pulses of varying durations depending on the event type were initiated by the experimental laptop at the onset of task events (trial and target sound onset, questions, button presses), generated by an Arduino Uno (R3; Smart Projects) and delivered directly to either to an open, dedicated channel in the icEEG recording montage or to a digital trigger input port. The varying durations of the TTL pulses were used to encode different event types, and allowed us to temporally align task events with the icEEG recordings.

### 2.5. Epoch segmentation and preprocessing

Four-second epochs centered at the onset of the target stimulus were extracted for each trial via MATLAB 2019a. Trial epochs containing clinical or subclinical epileptic events determined by visual inspection were excluded from the analysis. The remaining epochs were processed using a four-stage artifact rejection pipeline adapted from previous intracranial EEG studies within the lab ^4, 46–48^; at each stage, only trials that survived previous steps were included in analyses. First, on an electrode basis, each epoch’s power spectrum was estimated using Welch’s power spectral density method in MATLAB 2019a. Epochs displaying a topographical prominence greater than 200 μV/Hz in a frequency peak above 30 Hz were rejected to eliminate trials with excess electrical line or other high frequency noise. Next, on an electrode and epoch basis, the meansquare error (MSE) relative to zero was calculated; trials with an MSE of less than 200 μV^2^ were discarded to remove low amplitude signals resulting from loose or unplugged electrodes. Third, again on an electrode basis, the mean voltage time course was calculated for the remaining epochs. Then, on an epoch basis, the MSE between each trial and the mean trace for that electrode was calculated. Any epochs that exceeded an MSE of 3000 μV^2^ were considered excessively noisy. If the number of epochs that exceeded this threshold was less than or equal to 20% of the epochs for that electrode, all epochs above this threshold were excluded; if more than 20% of epochs exceeded this threshold, the noisiest 20% of epochs were discarded (following Herman, Smith ^4^). Lastly, the standard deviation was calculated across epochs for each electrode; epoch data from any given electrode were rejected if one or more samples in that epoch for that electrode had a voltage amplitude more than 5 standard deviations away from the electrode’s mean.

### 2.6. Broadband z-score gamma power calculation

Epochs that passed initial artifact rejection underwent spectral analysis to extract timeseries of broadband gamma power (defined here as 40–115 Hz)—a frequency band that has been shown to reflect local neural activity ^40–42, 49^. Gamma frequencies were extracted by applying a bandpass infinite impulse response filter (IIR, 40^th^-order, lower 3-dB frequency 40 Hz, higher 3-dB frequency 115 Hz) using the filtfilt function implemented with designfilt in MATLAB 2019a. Gamma power was calculated by squaring the filtered signal and then averaging within 100 ms windows beginning with the first epoch sample; bins were moved forward 25 ms such that there was a 75 ms overlap across bins to smooth the power signals over time. This resulted in 157 windows that tiled from 2000 ms preceding stimulus onset to 2000 ms following stimulus onset. In a final artifact rejection step, epochs were excluded if they contained a window that exceeded 20 standard deviations of the smoothed gamma power mean, calculated across epochs within a given electrode.

Epochs were then sorted by trial type, and the data was pruned to remove the first and last 500 ms of each epoch to eliminate filter edge effects. We found that the data from 1500 to ~500 ms pre-stimulus were not quiescent due to residual activity from the prior question/answer period, and therefore not ideal to use as a baseline period. Therefore, we used the period of 500 ms pre-stimulus to 0 ms (stimulus onset) as the ‘baseline’ period for all analyses. A common baseline was calculated across Perceived and Not Perceived trials, to ensure that post-stimulus differences described cannot be attributed to normalizing Perceived and Not Perceived trials to differential pre-stimulus states. This common baseline was calculated on an electrode basis by averaging the mean gamma power from −500 ms to 0 ms within trial for all Perceived and Not Perceived trials. The resulting trial-wise baseline time averages were then used to (1) calculate a standard deviation across all Perceived and Not Perceived trials for a given participant and (2) averaged across all Perceived and Not Perceived trials for a given participant to arrive at a baseline grand mean. This standard deviation and baseline grand mean were used to calculate each Perceived and Not Perceived trial’s z-score. The electrode’s mean z-scored gamma powers for Perceived and Not Perceived trials were calculated by averaging the resulting z-scores within trial type.

### 2.7. Mapping z-scored gamma power to brain

BioImage Suite was used to determine electrode locations in MNI space (Colin 22 brain template) based on each patient’s pre-op MRI and post-op CT. A triangular mesh representing the standard MNI cortical surface was created in BioImage Suite (http://bioimagesuite.yale.edu/), and electrodes were assigned to the nearest vertex. To visualize the entire cortical surface—and importantly for our study, the auditory cortex—we decided to display our data on an inflated brain. To do so, we used the MNI2FS MATLAB toolbox to render MNI coordinates on a canonical FreeSurfer inflated brain mesh^50^. All brain visualizations within our paper are shown at a level 5 (max available: 6) inflation. Z-scored gamma power values associated with each electrode were mapped onto the inflated brain vertices as described previously ^4, 48^. In short, to create a biologically feasible model of neural activity that was measured at each electrode, for each of a participant’s electrodes, all vertices within a 1mm spherical radius from the electrode’s assigned vertex were set to the same z-score value. Linear interpolation descending from the full z-score value to 0 was performed from a radius of 1 to 15 mm relative to the electrode’s vertex. If a participant had two or more electrodes that had overlapping vertices within their 15 mm radius, the z-scores calculated for each contributing electrode at that vertex were summed. Z-scores of each vertex were then combined across participants using a weighted average of the participants’ z-scores as previously described, consisting of the average z-score across participants at each vertex multiplied by the square root of number of participants with contributing data at that vertex ^4, 46–48^. This process was separately repeated for each 100 ms timepoint with 25 ms overlap.

### 2.8. Statistical analyses

To help overcome the multiple comparisons problem faced by high-dimensional datasets, we used a spatiotemporal cluster-based permutation test to identify the statistically significant changes in post-stimulus z-scored gamma power compared to the baseline ^51^. To do so, we divided the brain vertex surface into bilateral parcellation maps comprising 80 non-overlapping regions (40 regions per hemisphere) (Figure 2B). As previously described by Khalaf, Kronemer ^52^, this was achieved by applying a *k*-means clustering algorithm to the 3-dimensional coordinates of the brain map vertices such that vertices in close proximity were clustered into parcels. One hundred (100) such maps with slightly different parcels were generated for statistical analysis (see below). Electrodes were assigned to the parcel in which their center vertex fell. We then applied cluster-based permutation tests, adapted from the Mass Univariate ERP Toolbox ^53^ to the data. Separate sets of permutation tests were run for Perceived trials, for Not Perceived trials, and for Perceived-Not Perceived trials. For Perceived – Not Perceived trials, the electrode-level gamma power z-scores for Not Perceived trials were subtracted from those of Perceived trials before statistics were calculated.

**Figure 2.**
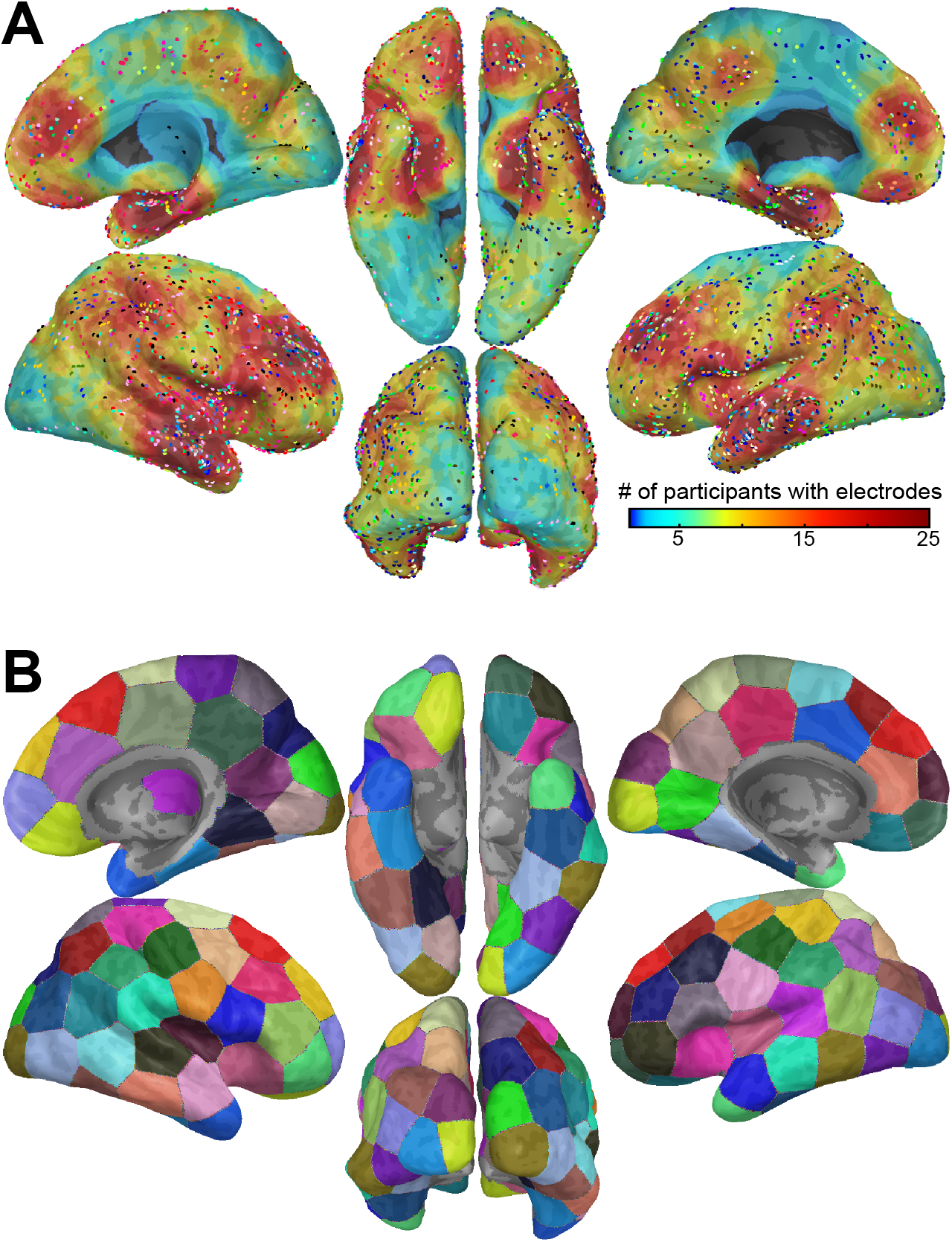
Visualizations of electrode distribution and parcellation. **(A)** Distribution of electrodes in the 31 study participants. Lateral, medial, ventral and posterior views of both hemispheres are shown on an inflated brain. Each participant’s gray matter electrodes are colored by a different color and displayed at the closest vertex. Background color (on inflated brain) display the number of participants contributing to the signal at each location on the brain surface. Total number of grey matter electrodes across all subjects = 2805. **(B)** Example of the 80-parcel bilateral parcellation map (40 parcels per hemisphere) generated using k-mean clustering algorithm. 100 such parcellation maps were constructed to ensure statistical findings were robust to boundaries of parcellations (see Methods).

To perform statistics, we first created a cluster-based permutation distribution, where the permuted values consisted of mean z-scored gamma power across electrodes within parcels and 100-ms time windows compared to the corresponding baseline values using a paired, 2-tailed t-test. Mean baseline activity was defined as the mean z-scored gamma power in the 500 ms preceding the stimulus (see 2.6. Broadband z-score gamma power calculation). Mean baseline z-score values were calculated separately for Perceived, Not Perceived, and Perceived – Not Perceived conditions; the mean baseline z-score activity for each of these was non-zero. Before calculating the t-test for each permutation, we first randomly shuffled the sign of gamma power values for each electrode to be positive or negative - in effect, adding or subtracting it from baseline. The paired, 2-tailed t-test was then performed, which generated a set of t-values across parcels and 100-ms time windows. Spatiotemporal clusters were defined based on adjacent parcels in space and time, and therefore the cluster-based permutation test was run separately for each hemisphere. A parcel and time point were eligible to join a cluster based on spatiotemporal adjacency if the t-value fulfilled a set alpha threshold. Positive and negative clusters were defined separately using this threshold. Because we randomly shuffled the sign of gamma power prior to calculating the z-score, we assumed the permutation distribution to be symmetrical. This allowed us to facilitate the remaining calculations by retaining only negative clusters to create a one-sided distribution. Therefore, the alpha threshold was set at 0.025 (equivalent to 0.05 in a 2-sided distribution). Summed t-values for each spatiotemporal cluster were then computed by taking the sum of the t-values for all parcels and time points within each cluster. For each permutation, we retained only the negative cluster with the largest absolute t-value. This procedure was performed across 2000 permutations, and a permutation distribution was created from the largest absolute t-value from each permutation. To identify significant clusters in the unpermuted data, positive and negative clusters were identified separately. They were considered significant if their summed t-values had an absolute value above the outer 2.5% of the permutation distribution (equivalent to *p* < 0.05 for a 2-sided distribution). Vertices that fell within a given cluster were assigned that cluster’s significance status.

The *k*-means algorithm used to create our parcellation maps chose a random seed to initialize the center of parcels, and therefore the algorithm does not produce identical maps on repeated runs. This results in different sets of vertices being grouped together (i.e., parcel locations and boundaries differ across runs). To ensure that statistical testing was robust to changes in parcel locations and to provide for greater spatial resolution, we generated 100 separate parcellation maps by running the *k*-means algorithm 100 times. For each of these 100 parcellation maps, we ran a separate spatiotemporal cluster-based permutation test as described above. This yielded 100 binary significance maps, where each vertex was assigned a 1 if it fell within a significant cluster, or a 0 if it did not. We used a majority vote approach across the 100 resultant significance maps to determine whether a vertex should be considered significant; vertices that reached significance in more than 50% of the significance maps were considered significant. We display the weighted average z-scored gamma power (see Section 2.7) only for these data points (Figures 3 and 4; Supplementary Presentation S1).

**Figure 3.**
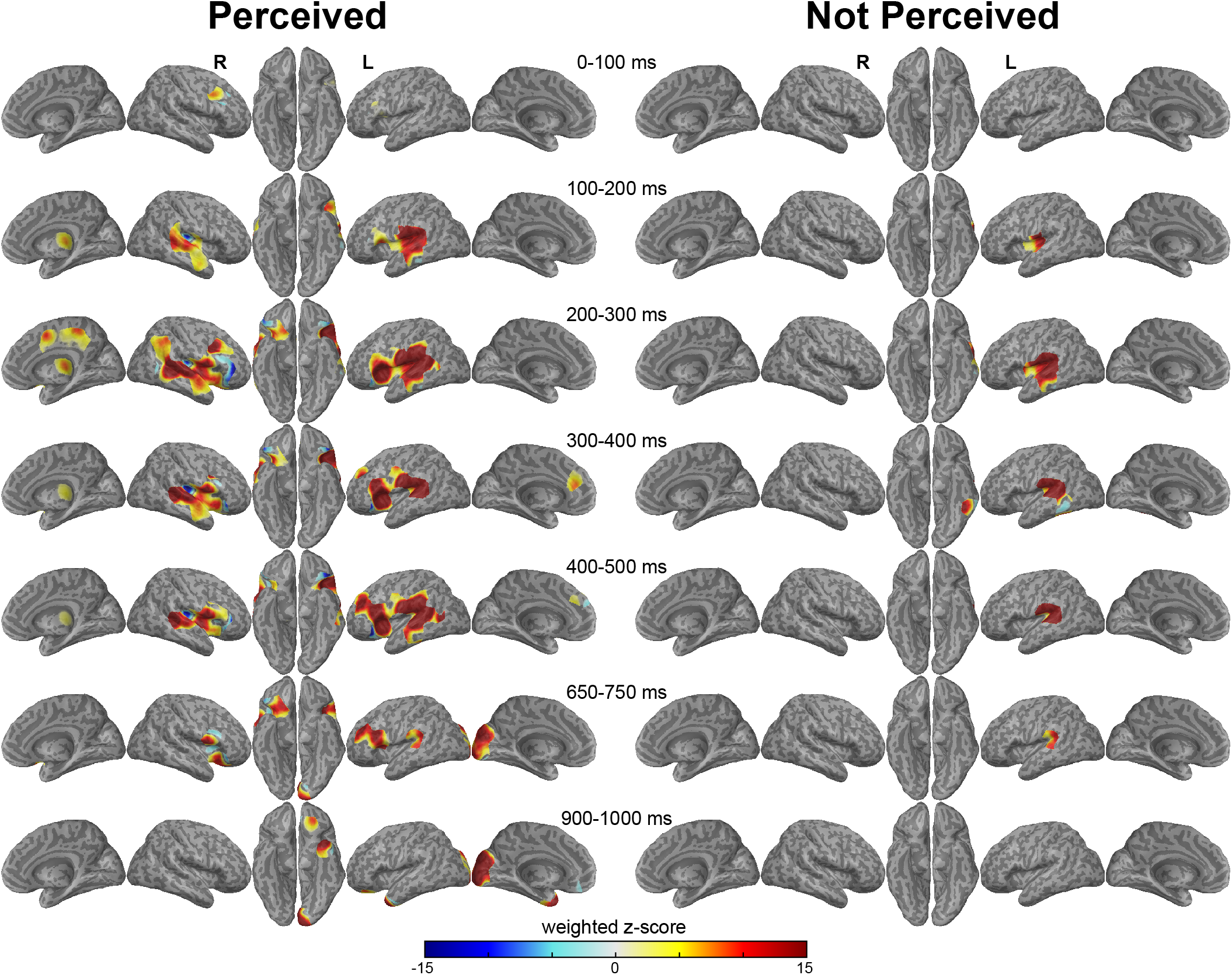
Statistically significant gamma power changes in Perceived and Not Perceived trials for threshold auditory stimuli. **(A)** Perceived trials. Gamma power increases were first observed in the right caudal frontal gyrus (overlapping with the frontal eye fields) and soon thereafter in bilateral early auditory regions and central thalamic areas (~100-200 ms after stimulus presentation). Increases spread into higher order auditory and frontal regions, and persisted until approximately 750 ms post stimulus. **(B)** Not Perceived trials. Significant activity was restricted to the left early auditory and auditory adjacent regions. However, it persisted from ~100-750 ms post stimulus onset. Vertices are only shown if they were found to be statistically significant during that timepoint; color reflects weighted z-scored gamma power. N = 31 participants. For display of all 100 ms time points at 25 ms intervals see Supplementary Presentation S1.

**Figure 4.**
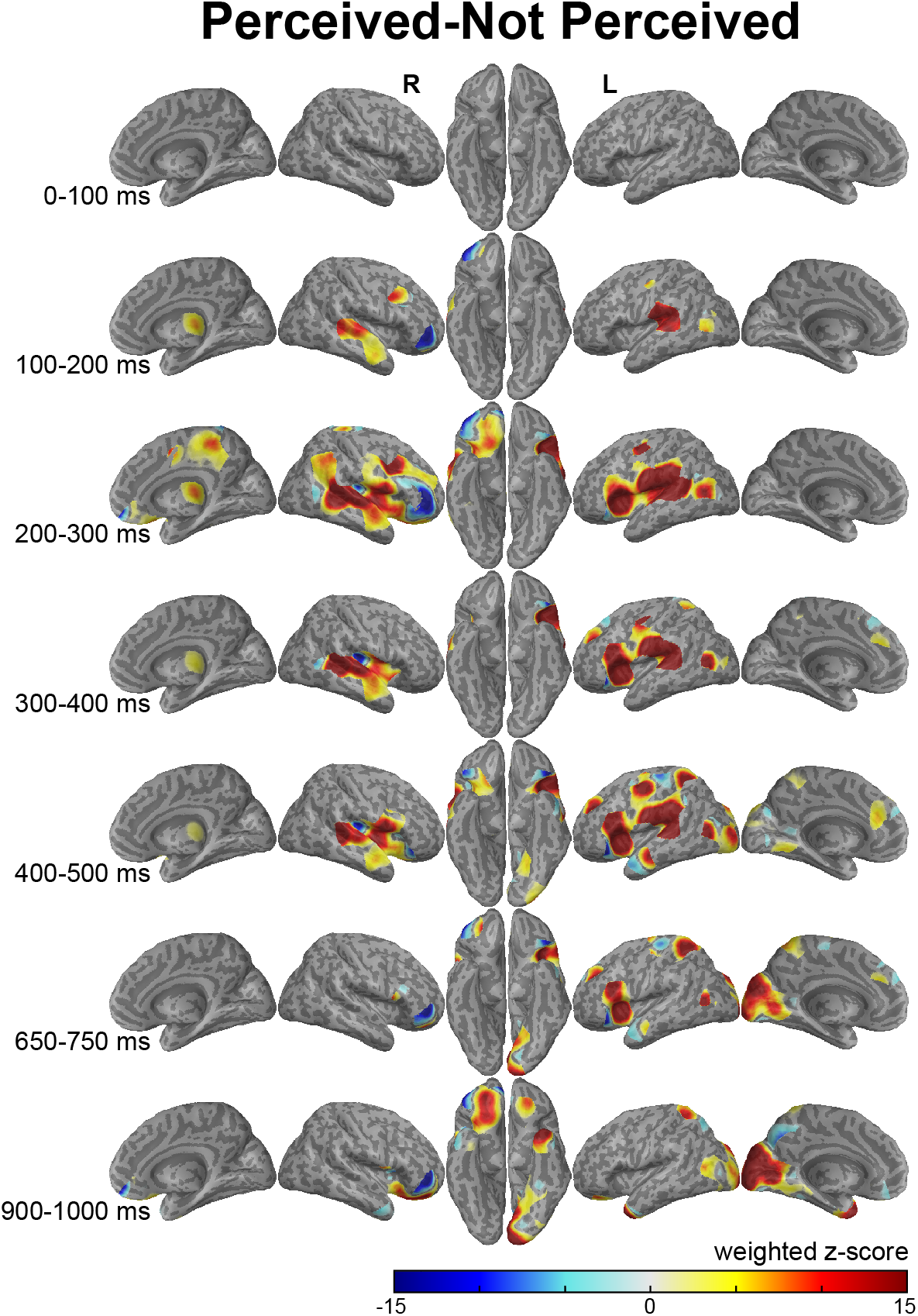
Statistically significant gamma power changes in Perceived – Not Perceived trials for threshold auditory stimuli. Differences in gamma power were first observed in the right caudal frontal gyrus (overlapping with the frontal eye fields), bilateral early auditory regions and central thalamic areas with greater increases for Perceived trials. The greater increases for Perceived trials then spread into higher order auditory and frontal regions, and persisted until approximately 750 ms post stimulus. Relative decreases were noted to be greater mainly for Perceived trials in ventrolateral regions of the right prefrontal cortex. Vertices are only shown if they were found to be statistically significant during that timepoint; color reflects weighted z-scored gamma power. Same data and participants as in Figure 3. N = 31 participants. For display of all 100 ms time points at 25 ms intervals see Supplementary Presentation S1.

### 2.9. Regions of interest

The cortical mesh was divided into anatomical regions by customizing preexisting regions of interest (ROIs) from MarsBaR ^54^ and the Harvard-Oxford Atlas (HOA) (https://fsl.fmrib.ox.ac.uk/fsl/fslwiki/Atlases). Customization was necessary to ensure that grey-matter vertices making up the cortical mesh were assigned to precisely one ROI (i.e., that no vertex within regions of interest were unassigned, and there were no overlapping borders) when projected onto the mni2fs inflated brain. Six anatomical ROIs involved in auditory perception and showing task-specific neural changes were selected for further comparison. The ROIs (Figure 5A) included bilateral representations of the early auditory regions (a combination of Heschl’s gyrus (MarsBaR Heschl’s and HOA Heschl’s H1/H2), the planum temporale (HOA planum temporale) and the posterior superior temporal gyrus (HOA superior temporal gyrus, posterior); the caudal middle frontal gyrus (HOA middle frontal gyrus, caudal portion only); the anterior insula (HOA insular cortex, anterior portion only); the inferior frontal gyrus pars opercularis (HOA inferior temporal gyrus, pars opercularis); the supramarginal gyrus (HOA supramarginal gyrus, anterior and posterior combined); and the thalamus (restricted to the right hemisphere, as there were no electrodes in the left thalamus; MarsBaR thalamus).

**Figure 5.**
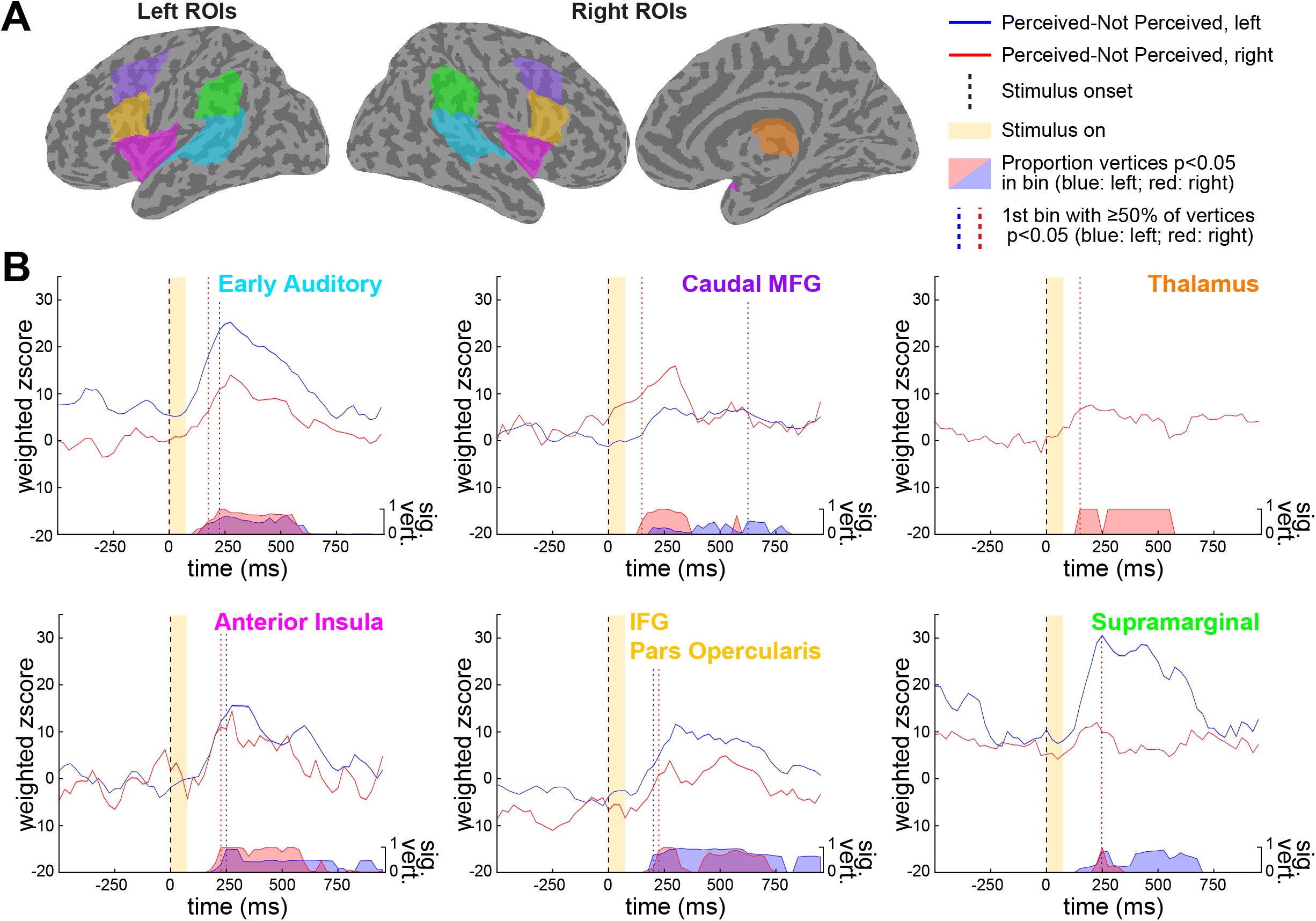
Gamma power time courses of 6 ROIS. (**A**) Six anatomical regions of interest (ROIs) were selected. The ROIs include early auditory regions, the caudal middle frontal gyrus (MFG), the thalamus, the anterior insula, the inferior frontal gyrus (IFG) pars opercularis and the supramarginal gyrus. Their locations are displayed on ventral (left and right) and medial (right) views on a brain. Electrodes were only present in the right thalamus, and so the left is not displayed. (**B**) Perceived – Not Perceived Analysis. Upper traces in each plot show average activity of all vertices within the ROI that achieved significance at any time point. Lower traces show the proportion of vertices that were significant (sig. vert.) for each time bin. Vertical colored dotted lines indicate the time bin during which 50% or more of significant vertices first achieved significance for the ROI in the left (blue) and right (red) hemisphere. The earliest significant changes were observed in early auditory, caudal MFG and thalamic regions (top row); slightly later changes in gamma power were observed in anterior insula, pars opercularis and the supramarginal gyrus. Vertical black dashed lines and yellow shaded regions, respectively, indicate onset timing and duration of the 75ms auditory stimulus. Same data and participants as in Figure 4 (N = 31).

For each ROI, vertices within that ROI that reached significance (defined as above, where the vertex was significant in >50% of significance maps) at any of the analyzed timepoints were included as part of that ROIs population at *all* timepoints. ROI timecourses took the average of the gamma power z scores across all included vertices at each time point; gamma power z scores for each vertex were calculated as described above (see section 2.7. Mapping z-scored gamma power to brain). The Perceived – Not Perceived timecourse for the ROIs were displayed separately for each hemisphere (Figure 5B).

As an additional measure of the magnitude of changes over time, and the earliest times that these changes occurred, we also calculated and displayed the proportion of vertices which reached significance at each time point. This is displayed on the axis below the timecourse data; where 1 indicates that all vertices that reached significance were significant at *that* timepoint, and 0 indicates that no significant vertices were significant at that timepoint (see Figure 5B). To assess the earliest times that changes occurred, we also calculated the first timepoint at which ≥50% or more of the significant vertices for that ROI achieved significance (displayed as the vertical blue and red dotted lines in Figure 5B).

### Data Availability

All data are available from the manuscript and the supplementary materials. Data are available at http://christisonlagay-blumenfeld-data.yale.edu <<Note: this URL will be open at the time of publication>>

### Code Availability

All codes are available from the manuscript and the supplementary materials. Codes are available at http://christisonlagay-blumenfeld-data.yale.edu <<Note: this URL will be open at the time of publication>>

## 3. Results

### 3.1. Behavior

On average, participants reported hearing a target stimulus for 58.0 % (±2.0% SEM) of trials in which a target was present and in 8.5% (± 1.4%) of trials in which a target was not present (blanks) (Figure 2B). Trial reporting remained consistent across the three post-stimulus delays and across target stimulus identity (delays: 3 seconds: 58.7 ± 2.2%; 4 seconds: 56.8 ± 2.1%; 5 seconds: 58.3 ± 2.0%; errors given in SEM; 1-way ANOVA across delay condition revealed no significant difference, [F(2,90)=0.25;*p*=0.78]; stimulus identity: whistle: 56.2 ± 2.8%; laser: 63.1 ± 2.3%; waterdrop: 54.4 ± 2.9%; errors given in SEM; 1-way ANOVA across stimulus revealed no significant difference, [F(2,88)=2.94; *p* =0.06])). A consistent proportion of stimuli reported over different delay times suggests that when stimuli are not reported this is truly due to lack of perception, and not due to forgetting that stimuli were perceived ^4^.

When participants indicated they heard a target for non-blank trials, they correctly indicated the sound’s identity in 89.2% (± 1.4% SEM) of trials; sound identification accuracy for unheard trials was close to, but slightly above, chance (40.2 ± 2.0% SEM; chance is 33.3%) (Figure 3C). Identification accuracy across targets remained consistent for heard trials (whistle: 90.1 ± 1.5%; laser: 89.1 ± 1.7%; waterdrop: 88.4 ± 1.8 %; errors given in SEM; 1-way ANOVA across stimulus revealed no significant difference, [F(2,88)=0.3; *p* =0.74]), and was also consistent for not heard trials (whistle: 32.6 ± 4.5%; laser: 40.1 ± 4.1%; waterdrop: 42.5 ± 4.9%; errors given in SEM; 1-way ANOVA across stimulus revealed no significant difference, [F(2,88)=1.36; *p*=0.26]). Participants completed 393 trials (7.9 runs) on average (median=350 trials [7 runs]; range=150–1208 trials). Because there were no significant differences across target stimulus identity or post-stimulus delay, we combined data across these trial conditions. Trials that were reported as heard and in which sounds were correctly identified were considered validated as ‘Perceived’; and trials reported as not heard and in which identification was incorrect were considered as validated ‘Not Perceived’ (Figure 2C).

### 3.2. Temporal and spatial signals

#### 3.2.1 Statistical brain maps

To investigate the spatiotemporal progression of neural activity related to conscious auditory perception, we collected data from a large number of intracranial EEG electrodes spread across both hemispheres of the cerebral cortex. We compared broadband gamma power (40–115 Hz; this frequency band has been associated with the firing rate of the local neuronal population ^40–42, 49^ associated with stimuli that were perceived (Figure 3, left panel) or not (Figure 3, right panel), and the difference in activity associated with perceived vs not perceived stimuli (Figure 4).

The earliest significant change in Perceived trials was an increase in gamma power, and somewhat surprisingly, this earliest change was observed in the right caudal middle frontal gyrus—an area close to the frontal eye fields ^55^ (Figure 3). The timing of this earliest change with auditory perception began in the bin centered at 25 ms after target stimulus onset (target duration = 75 ms; see Supplementary Presentation S1). This was followed by increases bilaterally in the auditory cortex accompanied by bilateral inferior frontal increases (Figure 3 Perceived trials; see also Supplementary Presentation S1, bin center: 75 ms), and by an increase in the right thalamus (Figure 3 Perceived trials; bin center: 125 ms, Supplementary Presentation S1; thalamic coverage was restricted to the right hemisphere). These increases expanded outwards from early auditory regions to the anterior insula (bilateral), frontal gyri (left: superior frontal gyrus; bilateral: inferior and middle frontal gyri), orbital frontal cortex (right hemisphere) and the parietal supramarginal gyrus (bilateral).

In the right hemisphere, this wave that expanded outward from the early auditory regions began to recede ~300 ms post stimulus onset. The significant increase in the right caudal middle frontal gyrus decreased at 350 ms post stimulus onset, and disappeared by 375 ms post stimulus onset. The last significant activity in the thalamus was observed at 450 ms post stimulus onset; significant increases persisted in the anterior insula until 550 ms, early auditory regions until 625 ms, inferior frontal gyrus until 800 ms, and in orbital frontal cortex until 850 ms post stimulus onset.

In the left hemisphere, significant increases persisted longer and with greater spatial extent than in the right hemisphere (Figure 3; Supplementary Presentation S1). The wave of significant increases in the left hemisphere began to recede at ~600 ms post stimulus onset, with activity that persisted (but in decreasing spatial extent and magnitude) in early auditory regions, supramarginal gyrus, anterior insula, pars opercularis, and anterior middle frontal gyrus. The last activity significant over baseline was observed at 700 ms for early auditory regions and the supramarginal gyrus, at 725 ms in the inferior frontal gyrus pars opercularis, at 775 ms in the anterior middle frontal gyrus and at 825 ms post stimulus onset in the anterior insula. Unlike the right hemisphere, which had no significant increases or decreases after the wave subsided, the left hemisphere showed increases in medial visual cortex beginning 600 ms and in the orbital frontal cortex and temporal pole beginning at 750 ms post stimulus onset; these three areas continued to show significantly increased activity past the end of our analysis window.

Although the significant activity in Perceived trials was dominated by increases in gamma power, it is of note that the right posterior insula (from 100–525 ms) and small regions contained within the inferior frontal gyrus pars orbitalis and orbital frontal cortex (bilaterally, from ~200–825 ms) showed decreases relative to baseline.

The significant changes observed in Not Perceived trials were more limited in their spatial extent than those seen in Perceived trials; indeed, significant changes were limited to the left hemisphere (Figure 3; Supplementary Presentation S1). The earliest significant changes in Not Perceived occurred at 150 ms post stimulus onset; this included a modest decrease in gamma power in the posterior insula, and modest increases in the anterior insula, a small portion of early auditory regions and in the central operculum/upper bank of the Sylvian fissure. The increases observed in the early auditory regions and in the central operculum spread and strengthened until 425 ms, moving somewhat posterior towards the supramarginal gyrus and temporoparietal junction at later timepoints. By 450 ms, these began to decrease in strength and size; the last significant activity for these regions is noted at 800 ms post stimulus onset. The modest decrease in gamma power in the posterior insula got smaller and weaker over time; the final observed decrease occurred at 725 ms post stimulus onset. Decreases in gamma power were also observed in the middle temporal (and to a lesser extent, inferior temporal) gyrus from 225–375 ms post stimulus, and then briefly again from 725–750 ms.

Finally, to visualize the difference between Perceived and Not Perceived trials, we analyzed the resultant activity of Perceived trials – Not Perceived trials (Figure 4. Using the Perceived – Not Perceived signal, our earliest significant change was centered at 100 ms post stimulus onset, in the right early auditory region (Supplementary Presentation S1). This activity was quickly followed (in the next bin, centered at 125 ms) by expanded bilateral auditory activity, and increases in gamma power in the thalamus and right caudal middle frontal gyrus. These clusters of significant increases in Perceived – Not Perceived signals expanded outward to encompass larger portions of the lateral surface of the brain, including lateral temporal cortex, occiptotemporal and temporoparietal areas, the upper banks of the Sylvian fissure, insula (bilateral: anterior insula; left: posterior insula), and frontal gyri (bilateral: middle and caudal inferior frontal gyri; left: superior frontal gyrus). Significant Perceived – Not Perceived relative increases were also noted in orbital frontal cortex and the temporal pole. In the orbital frontal cortex, right hemisphere increases began in tandem with the wave of increases seen elsewhere across the cortex, but persisted until the end of the analysis window; increases in the left hemisphere began as the wave of increases observed elsewhere reached its peak and, like the right hemisphere, persisted until the end of the analysis window. Activity in the temporal poles began towards the end of the wave of increases observed across the lateral surfaces of the brain, and persisted beyond the end of the analysis window. On the medial aspects of the brain, brief increases in gamma power were observed in the right precuneus and posterior portions of the right cingulate from 200–275 ms; brief, early activity in the right medial visual cortex was observed from 300–325 ms; and sustained activity in the left medial visual cortex began at 375 ms (and continued through the end of analysis window), which then extended along the ventral aspect of the occipital and temporal lobes (until ~650 ms post-stimulus onset). The differences in activity between Perceived and Not Perceived trials also highlighted decreases below baseline gamma power in the ventrolateral regions of prefrontal cortex, including the pars orbitalis and frontal pole, which began ~150 ms post stimulus onset and continued (with brief periods of failing to reach significance) until the end of the analysis window. These decreases were noted mostly in the right hemisphere, though small clusters of significant decreases occurred in the left pars orbitalis, as well as left orbital frontal regions (including a portion of medial orbital frontal cortex), left medial superior frontal gyrus, and left superior precentral gyrus.

#### 3.2.1 Region of interest timecourses

The ROI timecourses (Figure 5) summarize the findings observed in the Perceived – Not Perceived cortical surface maps (Figure 4). On the top row of Figure 5B, we show the ROIs with slightly earlier signals: the early auditory regions, the caudal middle frontal gyrus, and the thalamus (timing of earliest changes indicated by vertical colored lines, see Figure 5B legend). The early auditory regions showed an early surge of activity shortly after the stimulus onset, but this activity remained robust for an extended period of time. In contrast, the right caudal middle frontal gyrus and the thalamus show early activity that drops relatively quickly. The bottom row of timecourses show regions with slightly delayed peaks of activity: the anterior insula, the inferior frontal gyrus (IFG) pars opercularis and the supramarginal gyrus. All three regions had timecourses that, aside from a slight delay relative to the early auditory regions, followed a similar pattern to those same auditory regions, peaking ~250 ms after stimulus onset and decreasing back to baseline by ~750 ms following the stimulus. However, activity in the anterior insula and IFG pars opercularis—both part of the salience network—was notably less robust than that observed in the supramarginal gyrus. Finally, several regions displayed baselines that were offset from zero: left early auditory and supramarginal regions had prestimulus activity that was above zero, and the right IFG pars opercularis showed a negative baseline period (though none of these timepoint reached significance; see Discussion). Clear hemispheric asymmetries already described in the brain maps (Figure 4) can also be seen in the ROI timecourses for Perceived – Not Perceived data (Figure 5B), including larger overall increases in the left early auditory cortex and left supramarginal gyrus, and larger increases in the right caudal MFG.

## 4. Discussion

Here we show through direct human recordings that widespread cortical and subcortical networks are rapidly involved in conscious auditory perception, whereas identical not perceived stimuli elicit signals mainly confined to early auditory cortex. We developed a novel auditory threshold task that was designed to be an auditory correlate of a previously reported visual paradigm used in the lab ^4^. This new paradigm provides robust behavioral results that allow auditory stimuli, presented under identical physical circumstances, to result in approximately equal numbers of stimuli that are perceived and not perceived. By using this task in conjunction with large scale recordings from patients undergoing intracranial electroencephalography and through measuring neural activity by quantifying power in broadband gamma frequencies, we showed that a rapid progression of neural events closely followed a consciously perceived auditory stimulus; and that the scope of events associated with not-perceived stimuli is limited almost entirely to primary and other nearby early auditory cortex. Our findings expand the switch-and-wave model previously put forward by Herman, Smith ^4^, and support more recent work done conducted with scalp EEG, fMRI and pupillometry ^5^ by adding two additional components: detect and pulse ^47, 52, 56^. In this expanded model ^6^, consciously perceived stimuli elicit a series of neural events: (1) ***detect***: an initial activation of early sensory and other areas critical in the detection of incoming sensory stimuli (such as the frontal eye fields); (2) a ***pulse*** of activity originating from subcortical regions such as the thalamus; (3) a monophasic ***wave*** of activity that propagates from sensory regions to higher-order and association cortical regions and (4) the ***switch***ing off of default-mode related regions.

Examination of the activity in Perceived trials and in the difference between Perceived and Not Perceived trials over time showed activation of the canonical auditory pathways ^16, 17, 57–62^. This begins with a detection signal in the early auditory regions and then propagates outwards in a wave of activity to higher order regions. The earliest activity in early auditory regions happened within 100 ms of the onset of the target in both Perceived and Not Perceived trials. The timing of this early sensory activation, even in the absence of perception, mirrors findings in previous studies across sensory modalities ^4, 47, 63^; and indeed, is also supported by findings at the single-neuron level in non-human primates ^64–66^. Although the early auditory activity was present in both Perceived and Not Perceived trials, it is important to note that this early sensory activity was *greater* when the target was perceived, also consistent with previous findings ^4, 66^.

Although the onset of this early sensory activity was consistent with the visual version of the task ^4–5^, the dynamics of activation differed. Whereas in the visual task, gamma power in the primary visual areas showed an increase (<200 ms), followed by a decrease below baseline (200–600 ms), followed by a secondary increase (>600 ms), thus leading us to include the primary visual areas as a component of the ‘switch’, here we saw a sustained increase in early auditory regions. This increase in gamma power persisted until approximately 800 ms after target onset; indeed, the early auditory activity persisted for the entirety of the wave of activity that spread to higher order and association regions. What leads to this marked difference between auditory and visual processing? Sustained activity in the auditory cortex following stimulus offset has been seen before for tasks with working memory demands (which might suggest that such activity is important for storing auditory information), but also in passive listening tasks ^67–71^. Sustained activity in the visual cortex is sometimes observed when there is a need for visual working memory ^72–75^, although transient decreases below baseline are also reported ^48, 76, 77^. It may be that because our previous task used a face stimulus, we would expect sustained activity in the fusiform gyrus (which, indeed, we did observe), whereas early visual regions may need to deactivate in order to be prepared for the next stimulus ^78^. Of course, there may be other reasons for differential activity in auditory and visual domains. Across the phylogenetic tree, auditory perception requires temporal integration ^79^, and this may require sustained auditory activity. Our observation that such activity is sustained— albeit in a more circumscribed manner—even in the absence of perception (1) supports the hypothesis that such activity is not driven by task demands or working memory and (2) suggests that primary sensory activity is insufficient for perception.

Activity in Not Perceived trials was constrained to this sustained activity in early auditory regions, but Perceived (and when comparing Perceived – Not Perceived) trials showed a wave of activity that propagated outward from early auditory regions and along the canonical auditory pathways. The auditory processing pathways are generally split into two distinct processing streams: a ventral pathway, critical for sound perception and identification; and a dorsal pathway, important for timing, localization, and audio-motor responses, including speech ^16, 17, 57–62^. We observed activity in both branches. The most pronounced activity in frontal regions was consistent with activity along the ventral auditory pathway—specifically, in areas of the inferior frontal gyrus associated with the ventrolateral prefrontal cortex. This region is the terminus of the ventral auditory pathway; and because our task involved identifying specific target sounds, it makes sense that this pathway would be activated. The wave of activity extending outward from early auditory regions also took a postero-dorsal path, consistent with the dorsal pathway, with substantial increases in activity observed in the supramarginal gyrus. Although our task did not involve locating a sound, it did require participants to (1) identify *nameable* stimuli and (2) carry out a subsequent motor response. The activity that we noted in the supramarginal gyrus may be due to these components of the task as opposed to the actual perception of the stimulus.

Moving away from regions canonically associated with auditory responses, our analyses also revealed early activity in caudal middle frontal gyrus—an area consistent with the location of the frontal eye fields (FEF). The FEF was described initially for its role in controlling saccadic eye movements ^80^, but has subsequently been recognized to play an important role in in selective visual attention ^81–84^, visual detection and the accumulation of visual sensory evidence ^47, 52, 85^, and even the processing of auditory stimuli when there is a spatial component ^86^. Its rapid activation after a sensory stimulus, as well as its multimodal sensitivities, suggest that far from being a solely motor-planning region, it may in fact play an important role in sensory processing and perception ^87^. Our findings support a sensory detection role for the FEF: its increase in gamma power is almost simultaneous with that of early auditory processing regions. Of note is that we observe this significant increase *only* when the participant perceived the auditory target. This is at odds with the activity found in early auditory regions: while both Perceived and Not Perceived trials elicit activity in the auditory cortex (albeit, to differing extents), the caudal middle frontal gyrus shows a significant increase solely for Perceived trials (and when comparing the difference between Perceived and Not Perceived trials), indicating that while sensory detection may occur in both sensory regions and the FEF, their roles are likely not identical. This differentiation may in part be reflected in previous findings in single neurons within the FEF, which show activity that reflects the accumulation of sensory evidence, as well as being influenced by the eventual behavioral choice of monkeys performing a visual discrimination task ^85^. Additionally, to our knowledge, these results represent the first demonstration of FEF activity in an auditory task that does not involve a spatial processing component.

We also observed a pulse of activity from a central region of the thalamus, likely including the intralaminar nuclei, associated with a perceived stimulus (Perceived trials; and Perceived-Not Perceived trials). This pulse began very shortly after the initial ‘detect’ signal in the early auditory regions and caudal middle frontal gyrus, and persisted until approximately 450 ms after stimulus onset. Previous work has shown that the intralaminar thalamus may be a key subcortical structure for arousal and consciousness ^88, 89^; and recent work in our lab has described an evoked potential in the thalamus (thalamic awareness potential [TAP]) and a BOLD signal consistent with TAP using the previously described visual analog of the current auditory task ^5^. Our findings lend further evidence to a role of subcortical structures in conscious perception.

Additionally, we note that there is an increase in activity in the salience network and a decrease in the default mode network. The is consistent with our findings in visual paradigms ^4, 5, 48^. In the salience network, we find robust increases of activity in the insula, particularly the anterior insula, in Perceived (and Perceived-Not Perceived) trials. This is not unexpected, as the anterior insula has been routinely implicated in the perceptual processing of auditory (and visual) stimuli ^90^. We observed a decrease in default mode network regions— specifically, the inferior frontal pars orbitalis and orbital frontal regions (Xu, Yuan, and Lei, 2016)—when comparing Perceived-Not Perceived trials. Such a decrease is unsurprising, as the default mode has been shown to deactivate in response to task demands, internal planning, and working memory ^48, 91–96^. Furthermore, similar decreases were observed in default mode regions in our visual studies ^4, 5^. However, of note is that default mode regions seem to deactivate earlier in the auditory paradigm than in the equivalent visual paradim, and although studies in both modalities show default mode network downregulation, the exact contributing areas are not fully overlapping. These differences maybe due to the modality itself, or because of differential experimental parameters (e.g., the distribution of electrodes).

Several of our observations beg further scrutiny. One such finding pertains to the prestimulus baselines observed in various regions of the brain. Although baselines across Perceived and Not Perceived trials were similar in most regions (Fig 5), several areas displayed activity that diverged from zero (though no timepoint in prestimulus in any area reached significance), meaning that Perceived and Not Perceived trials had differing baselines. These differences were in auditory or auditory adjascent areas: early auditory regions (left hemisphere), supramarginal gyrus (left hemisphere), and the pars opercularis (right hemisphere). Although many studies have established that prestimulus activity influences likelihood of perception ^38, 97–99^, future studies may be able to establish whether, and which, areas are causally tied to perception.

One last finding of note is that the early visual cortex showed late activation in Perceived and Perceived – Not Perceived trials. A similar finding was shown in the visual paradigm ^4^, although this was interpreted as a possible mechanism for maintaining or retrieving the memory of the visual target ^72–75^. However, given that a similar activation was observed in a non-visual task at approximately the same time suggests that visual memory maintenance and/or retrieval may not fully explain this late visual activity. Because a visual noise stimulus was used in the auditory study, it is possible that after an auditory target was perceived, attention was reallocated to vision; but further study with additional modalities are needed to fully understand the cause and meaning of this late emerging activity.

Our previous work from ^4^ proposed a “switch and wave” model of conscious perception which expanded upon “ignition” models of consciousness, in which widespread brain activity followed a consciously perceived event ^100, 101^. However, Kronemer, Aksen ^5^, other recent work ^47, 52, 56^ and the current study expand upon that switch and wave model. Here, we provide evidence for a more complete model ^6^, including the following steps: (1) *detection*—in early sensory regions and the FEF; (2) a *pulse*—from subcortical arousal regions such as the intralaminar thalamus; (3) a *switch*—decreases in default mode regions and (4) a *wave*—of activity that sweeps from hierarchically from early sensory areas into higher order association areas. Many of our findings here bear remarkable similarity to our findings in equivalent visual paradigms ^4, 5^, which suggests that there may be shared neural underpinnings of the detect/pulse/switch/wave model across sensory systems. However, it is notable that there are marked differences in the temporal dynamics of early sensory regions between visual and auditory modalities, which highlights the need for perceptual experiments across sensory modalities. Future investigation should also focus on the role of perceptual report: although having participants report their percepts to experiments is a mainstay of many perceptual paradigms, the act of reporting introduces additional processing (working memory, motor planning, etc) ^102, 103^. The future development of no-report paradigms used in conjunction with electrophysiological experiments across sensory modalities will help determine the spatiotemporal dynamics of conscious perception itself.

## Supporting information

Supplemental presentation 1

## Acknowledgments

We thank our subjects and their families for their patience and willingness to participate in this research. We thank the staff of the Yale Comprehensive Epilepsy Center for their help.

## Author contributions

Conceptualization: HB, KLCL

Formal analysis: KLCL, CM, NCF

Funding acquisition: HB

Investigation (data collection): KLCL, NCF, CM, SIK, MMG, LK, SF, JD, MA, NM, EY, EE, JH, SB, JY, AM, JG, ED, DS

Methodology (design): KLCL, SIK, AK, HK, HB

Project administration: HB, KLCL

Supervision: HB

Writing – original draft: KLCL, HB

Writing – review & editing: KLCL, HB, NCF, CM, SIK, AK, MMG, LK, SF, JD, MA, AAA, HK, NM, EY, EE, JH, SB, JY, AM, KW, JG, ED, DS

## References

1. Koch C. The quest for consciousness: a neurobiological approach. Denver, Colo.: Roberts and Co.; 2004. xviii, 429 p. p.

2. Del Cul A, Baillet S, Dehaene S. Brain dynamics underlying the nonlinear threshold for access to consciousness. Plos Biology. 2007;5(10):e260.

3. Tononi G, Boly M, Massimini M, Koch C. Integrated information theory: from consciousness to its physical substrate. Nat Rev Neurosci. 2016;17(7):450–61.

4. Herman WX, Smith RE, Kronemer SI, Watsky RE, Chen WC, Gober LM, et al. A Switch and Wave of Neuronal Activity in the Cerebral Cortex During the First Second of Conscious Perception. Cereb Cortex. 2019;29(2):461–74.

5. Kronemer SI, Aksen M, Ding J, Ryu JH, Xin Q, Ding Z, et al. Brain networks in human conscious visual perception. bioRxiv. 2021:2021.10.04.462661.

6. Blumenfeld H. Brain Mechanisms of Conscious Awareness: Detect, Pulse, Switch, and Wave. Neuroscientist. 2021:https://doi.org/10.1177/10738584211049378.

7. Bregman A. Auditory Scene Analysis: The Perceptual Organization of Sound. Cambridge, MA: MIT Press; 1990.

8. McDermott J. The cocktail party problem. Curr Biol. 2010;19:R1024–R7.

9. Shinn-Cunningham BG. Object-based auditory and visual attention. Trends Cogn Sci. 2008;12(5):182–6.

10. Sussman ES, Bregman AS, Wang WJ, Khan FJ. Attentional modulation of electrophysiological activity in auditory cortex for unattended sounds within multistream auditory environments. Cognitive Affective & Behavioral Neuroscience. 2005;5(1):93–110.

11. Winkler I, Denham SL, Nelken I. Modeling the auditory scene: predictive regularity representations and perceptual objects. Trends Cogn Sci. 2009;13(12):532–40.

12. Christison-Lagay KL, Gifford AM, Cohen YE. Neural correlates of auditory scene analysis and perception. International journal of psychophysiology. 2015;95(2):238–45.

13. Micheyl C, Carlyon RP, Gutschalk A, Melcher JR, Oxenham AJ, Rauschecker JP, et al. The role of auditory cortex in the formation of auditory streams. Hear Res. 2007;229(1-2):116–31.

14. Bizley JK, Cohen YE. The what, where and how of auditory-object perception. Nature Reviews Neuroscience. 2013;14(10):693–707.

15. Singh PG. Perceptual organization of compex-tone sequences: A tradeoff between pith and timbre? J Acoust Soc Am. 1987;82(3):886–9.

16. Kaas JH, Hackett TA. ‘What’ and ‘where’ processing in auditory cortex. Nat Neurosci. 1999;2(12):1045–7.

17. Romanski LM, Tian B, Fritz J, Mishkin M, Goldman-Rakic PS, Rauschecker JP. Dual streams of auditory afferents target multiple domains in the primate prefrontal cortex. Nat Neurosci. 1999;2(12):1131–6.

18. Griffiths TD. Sensory Systems: Auditory Action Streams? Curr Biol. 2008;18:R387–R8.

19. Razak KA. Systematic representation of sound locations in the primary auditory cortex. J Neurosci. 2011;31(39):13848–59.

20. Steinschneider M, Arezzo JC, Vaughan J, Herbert G. Tonotopic features of speech-evoked activity in primate auditory cortex. Brain Res. 1990;519:158–68.

21. Kim S, Schwalje AT, Liu AS, Gander PE, McMurray B, Griffiths TD, et al. Pre- and post-target cortical processes predict speech-in-noise performance. Neuroimage. 2021;228:117699.

22. Bhaya-Grossman I, Chang EF. Speech Computations of the Human Superior Temporal Gyrus. Annual Review of Psychology. 2022;73(1):null.

23. Steinschneider M, Nourski KV, Kawasaki H, Oya H, Brugge JF, Howard MA, III. Intracranial Study of Speech-Elicited Activity on the Human Posterolateral Superior Temporal Gyrus. Cerebral Cortex. 2011;21(10):2332–47.

24. Christison-Lagay K, Cohen Y. The contribution of primary auditory cortex to auditory categorization in behaving monkeys. Frontiers in Neuroscience. 2018;12:601.

25. Fishman YI, Reser DH, Arezzo JC, Steinschneider M. Neural correlates of auditory stream segregation in primary auditory cortex of the awake monkey. Hear Res. 2001;151:167–87.

26. Micheyl C, Hunter C, Oxenham AJ. Auditory stream segregation and the perception of across-frequency synchrony. J Exp Psychol Hum Percept Perform. 2010;36(4):1029–39.

27. Micheyl C, Kreft H, Shamma SA, Oxenham AJ. Temporal coherence versus harmonicity in auditory stream formation. J Acoust Soc Am. 2013;133(3):EL188.

28. Javitt DC, Steinschneider M, Schroeder CE, Vaughan HG, Jr., Arezzo JC. Detection of stimulus deviance within primate primary auditory cortex: intracortical mechanisms of mismatch negativity (MMN) generation. Brain Res. 1994;667(2):192–200.

29. Näatänen R, Paavilainen P, Rinne T, Alho K. The mismatch negativity (MMN) in basic research of central auditory processing: A review. Clin Neurophysiol. 2007;118:2544–90.

30. Yabe H, Tervaniemi M, Reinikainen K, Näatänen R. Temporal window of integration revealed by MMN to sound omission. Neuroreport. 1997;8:1971–4.

31. Gifford III GW, MacLean KA, Hauser MD, Cohen YE. The neurophysiology of functionally meaningful categories: macaque ventrolateral prefrontal cortex plays a critical role in spontaneous categorization of species-specific vocalizations. Journal of Cognitive Neuroscience. 2005;17:1471–82.

32. Hillyard SA, Squires KC, Bauer JW, Lindsay PH. Evoked potential correlates of auditory signal detection. Science. 1971;172(3990):1357–60.

33. Squires KC, Hillyard SA, Lindsay PH. Vertex potentials evoked during auditory signal detection: Relation to decision criteria. Percept Psychophys. 1973;14(2):265–72.

34. Sanchez G, Hartmann T, Fuscà M, Demarchi G, Weisz N. Decoding across sensory modalities reveals common supramodal signatures of conscious perception. Proceedings of the National Academy of Sciences. 2020;117(13):7437–46.

35. Sussman ES, Horváth J, Winkler I, Orr M. The role of attention in the formation of auditory streams. Percept Psychophys. 2007;69(1):136–52.

36. Sadaghiani S, Hesselmann G, Kleinschmidt A. Distributed and antagonistic contributions of ongoing activity fluctuations to auditory stimulus detection. J Neurosci. 2009;29(42):13410–7.

37. Leske S, Ruhnau P, Frey J, Lithari C, Müller N, Hartmann T, et al. Prestimulus Network Integration of Auditory Cortex Predisposes Near-Threshold Perception Independently of Local Excitability. Cerebral cortex (New York, NY: 1991). 2015;25(12):4898–907.

38. Herrmann B, Henry MJ, Haegens S, Obleser J. Temporal expectations and neural amplitude fluctuations in auditory cortex interactively influence perception. NeuroImage. 2016;124:487–97.

39. Zoefel B, Heil P. Detection of Near-Threshold Sounds is Independent of EEG Phase in Common Frequency Bands. Frontiers in psychology. 2013;4(262).

40. Mukamel R, Gelbard H, Arieli A, Hasson U, Fried I, Malach R. Coupling between neuronal firing, field potentials, and FMRI in human auditory cortex. Science. 2005;309(5736):951–4.

41. Manning JR, Jacobs J, Fried I, Kahana MJ. Broadband shifts in local field potential power spectra are correlated with single-neuron spiking in humans. J Neurosci. 2009;29(43):13613–20.

42. Miller KJ, Honey CJ, Hermes D, Rao RP, Dennijs M, Ojemann JG. Broadband changes in the cortical surface potential track activation of functionally diverse neuronal populations. Neuroimage. 2013.

43. Kleiner M, Brainard DH, Pelli D. What’s new in Psychtoolbox-3? Perception. 2007;36:1–16.

44. Pelli DG. The VideoToolbox software for visual psychophysics: transforming numbers into movies. Spat Vis. 1997;10(4):437–42.

45. Brainard DH. The Psychophysics Toolbox. Spatial Vision. 1997;10(4):433–6.

46. Khalaf A, Kronemer SI, Christison-Lagay K, Kwon H, Li J, Wu K, et al. Early neural activity changes associated with stimulus detection during visual conscious perception. Cerebral Cortex. 2022.

47. Kwon H, Kronemer SI, Christison-Lagay KL, Khalaf A, Li J, Ding JZ, et al. Early cortical signals in visual stimulus detection. Neuroimage. 2021;244:118608.

48. Li J, Kronemer SI, Herman WX, Kwon H, Ryu JH, Micek C, et al. Default mode and visual network activity in an attention task: Direct measurement with intracranial EEG. Neuroimage. 2019;201:116003.

49. Ray S, Maunsell JH. Different origins of gamma rhythm and high-gamma activity in macaque visual cortex. PLoS Biol. 2011;9(4):e1000610.

50. Price D. MNI2FS: Surface Rendering of MNI Space Volumes for MATLAB github 2017 [Available from: https://github.com/dprice80/mni2fs.

51. Maris E, Oostenveld R. Nonparametric statistical testing of EEG- and MEG-data. J Neurosci Methods. 2007;164(1):177–90.

52. Khalaf A, Kronemer SI, Christison-Lagay K, Kwon H, Li J, Wu K, et al. Early neural activity changes associated with stimulus detection during visual conscious perception. Cereb Cortex. 2022.

53. Groppe DM, Urbach TP, Kutas M. Mass univariate analysis of event-related brain potentials/fields I: a critical tutorial review. Psychophysiology. 2011;48(12):1711–25.

54. Tzourio-Mazoyer N, Landeau B, Papathanassiou D, Crivello F, Etard O, Delcroix N, et al. Automated anatomical labeling of activations in SPM using a macroscopic anatomical parcellation of the MNI MRI single-subject brain. Neuroimage. 2002;15(1):273–89.

55. Vernet M, Quentin R, Chanes L, Mitsumasu A, Valero-Cabré A. Frontal eye field, where art thou? Anatomy, function, and non-invasive manipulation of frontal regions involved in eye movements and associated cognitive operations. Frontiers in Integrative Neuroscience. 2014;8(66).

56. Li R, Ryu JH, Vincent P, Springer M, Kluger D, Levinsohn EA, et al. The pulse: transient fMRI signal increases in subcortical arousal systems during transitions in attention. Neuroimage. 2021;232:117873.

57. Rauschecker JP. Ventral and dorsal streams in the evolution of speech and language. Frontiers in evolutionary neuroscience. 2012;4:7.

58. Cohen YE, Bennur S, Christison-Lagay K, Gifford AM, Tsunada J. Functional Organization of the Ventral Auditory Pathway. Adv Exp Med Biol. 2016;894:381–8.

59. Kaas JH, Hackett TA. Subdivisions of auditory cortex and processing streams in primates. Proc Natl Acad Sci. 2000;97(22):1179–11799.

60. Hackett TA. Information flow in the auditory cortical network. Hear Res. 2011;271(1-2):133–46.

61. Rauschecker JP, Scott SK. Maps and streams in the auditory cortex: nonhuman primates illuminate human speech processing. Nat Neurosci. 2009;12(6):718–24.

62. Rauschecker JP, Tian B. Mechanisms and streams for processing of “what” and “where” in auditory cortex. Proc Natl Acad Sci U S A. 2000;97(22):11800–6.

63. Palva S, Linkenkaer-Hansen K, Naatanen R, Palva JM. Early neural correlates of conscious somatosensory perception. Journal of Neuroscience. 2005;25(21):5248–58.

64. Schmolesky MT, Wang Y, Hanes DP, Thompson KG, Leutgeb S, Schall JD, et al. Signal Timing Across the Macaque Visual System. Journal of Neurophysiology. 1998;79(6):3272–8.

65. Christison-Lagay KL, Bennur S, Cohen YE. Contribution of spiking activity in the primary auditory cortex to detection in noise. Journal of neurophysiology. 2017;118(6):3118–31.

66. Vugt Bv, Dagnino B, Vartak D, Safaai H, Panzeri S, Dehaene S, et al. The threshold for conscious report: Signal loss and response bias in visual and frontal cortex. Science. 2018;360(6388):537–42.

67. Coffey EBJ, Arseneau-Bruneau I, Zhang X, Baillet S, Zatorre RJ. Oscillatory Entrainment of the Frequency-following Response in Auditory Cortical and Subcortical Structures. J Neurosci. 2021;41(18):4073–87.

68. Huang Y, Matysiak A, Heil P, König R, Brosch M. Persistent neural activity in auditory cortex is related to auditory working memory in humans and nonhuman primates. Elife. 2016;5.

69. Brosch M, Selezneva E, Scheich H. Formation of associations in auditory cortex by slow changes of tonic firing. Hear Res. 2011;271(1-2):66–73.

70. Nolden S, Grimault S, Guimond S, Lefebvre C, Bermudez P, Jolicoeur P. The retention of simultaneous tones in auditory short-term memory: a magnetoencephalography study. Neuroimage. 2013;82:384–92.

71. Linke AC, Cusack R. Flexible information coding in human auditory cortex during perception, imagery, and STM of complex sounds. J Cogn Neurosci. 2015;27(7):1322–33.

72. Thakral PP, Slotnick SD, Schacter DL. Conscious processing during retrieval can occur in early and late visual regions. Neuropsychologia. 2013;51(3):482–7.

73. Barceló F, Suwazono S, Knight RT. Prefrontal modulation of visual processing in humans. Nat Neurosci. 2000;3(4):399–403.

74. Premereur E, Janssen P, Vanduffel W. FEF-microstimulation causes task-dependent modulation of occipital fMRI activity. Neuroimage. 2013;67:42–50.

75. Agam Y, Hyun JS, Danker JF, Zhou F, Kahana MJ, Sekuler R. Early neural signatures of visual shortterm memory. Neuroimage. 2009;44(2):531–6.

76. Hindi Attar C, Andersen SK, Müller MM. Time course of affective bias in visual attention: convergent evidence from steady-state visual evoked potentials and behavioral data. Neuroimage. 2010;53(4):1326–33.

77. Li Y, Lou B, Gao X, Sajda P. Post-stimulus endogenous and exogenous oscillations are differentially modulated by task difficulty. Front Hum Neurosci. 2013;7:9.

78. Dux PE, Marois R. The attentional blink: a review of data and theory. Attention, perception & psychophysics. 2009;71(8):1683–700.

79. Itoh K, Nejime M, Konoike N, Nakamura K, Nakada T. Evolutionary Elongation of the Time Window of Integration in Auditory Cortex: Macaque vs. Human Comparison of the Effects of Sound Duration on Auditory Evoked Potentials. Frontiers in Neuroscience. 2019;13(630).

80. Ferrier D. Experimental researches in cerebral physiology and pathology. British medical journal. 1873;1(643):457.

81. de Haan B, Morgan PS, Rorden C. Covert orienting of attention and overt eye movements activate identical brain regions. Brain Res. 2008;1204:102–11.

82. Paneri S, Gregoriou GG. Top-Down Control of Visual Attention by the Prefrontal Cortex. Functional Specialization and Long-Range Interactions. Frontiers in Neuroscience. 2017;11(545).

83. Buschman TJ, Miller EK. Serial, Covert Shifts of Attention during Visual Search Are Reflected by the Frontal Eye Fields and Correlated with Population Oscillations. Neuron. 2009;63(3):386–96.

84. Gregoriou GG, Gotts SJ, Zhou H, Desimone R. Long-range neural coupling through synchronization with attention. In: Srinivasan N, editor. Progress in Brain Research. 176: Elsevier; 2009. p. 35–45.

85. Fan Y, Gold JI, Ding L. Frontal eye field and caudate neurons make different contributions to reward-biased perceptual decisions. Elife. 2020;9.

86. Smith DT, Jackson SR, Rorden C. Repetitive transcranial magnetic stimulation over frontal eye fields disrupts visually cued auditory attention. Brain Stimul. 2009;2(2):81–7.

87. Kirchner H, Barbeau EJ, Thorpe SJ, Régis J, Liégeois-Chauvel C. Ultra-Rapid Sensory Responses in the Human Frontal Eye Field Region. The Journal of Neuroscience. 2009;29(23):7599–606.

88. Redinbaugh MJ, Phillips JM, Kambi NA, Mohanta S, Andryk S, Dooley GL, et al. Thalamus Modulates Consciousness via Layer-Specific Control of Cortex. Neuron. 2020;106(1):66–75 e12.

89. Schiff ND. Central Lateral Thalamic Nucleus Stimulation Awakens Cortex via Modulation of Cross-Regional, Laminar-Specific Activity during General Anesthesia. Neuron. 2020;106(1):1–3.

90. Protas M, editor Role of the Insula in Visual and Auditory Perception 2018.

91. Smallwood J, Brown K, Baird B, Schooler JW. Cooperation between the default mode network and the frontal-parietal network in the production of an internal train of thought. Brain Res. 2012;1428:60–70.

92. Miller KJ, Weaver KE, Ojemann JG. Direct electrophysiological measurement of human default network areas. Proc Natl Acad Sci U S A. 2009;106(29):12174–7.

93. Singh KD, Fawcett IP. Transient and linearly graded deactivation of the human default-mode network by a visual detection task. Neuroimage. 2008;41(1):100–12.

94. Ossandón T, Jerbi K, Vidal JR, Bayle DJ, Henaff MA, Jung J, et al. Transient suppression of broadband gamma power in the default-mode network is correlated with task complexity and subject performance. J Neurosci. 2011;31(41):14521–30.

95. Weissman DH, Roberts KC, Visscher KM, Woldorff MG. The neural bases of momentary lapses in attention. Nat Neurosci. 2006;9(7):971–8.

96. Li CS, Yan P, Bergquist KL, Sinha R. Greater activation of the “default” brain regions predicts stop signal errors. Neuroimage. 2007;38(3):640–8.

97. Rahn E, Başar E. Prestimulus EEG-activity strongly influences the auditory evoked vertex response: a new method for selective averaging. Int J Neurosci. 1993;69(1-4):207–20.

98. Kayser SJ, McNair SW, Kayser C. Prestimulus influences on auditory perception from sensory representations and decision processes. Proc Natl Acad Sci U S A. 2016;113(17):4842–7.

99. Ronconi L, Pincham HL, Cristoforetti G, Facoetti A, Szucs D. Shaping prestimulus neural activity with auditory rhythmic stimulation improves the temporal allocation of attention. NeuroReport. 2016;27(7):487–94.

100. Fisch L, Privman E, Ramot M, Harel M, Nir Y, Kipervasser S, et al. Neural “ignition”: enhanced activation linked to perceptual awareness in human ventral stream visual cortex. Neuron. 2009;64(4):562–74.

101. Dehaene S, Changeux JP. Experimental and theoretical approaches to conscious processing. Neuron. 2011;70(2):200–27.

102. Aru J, Bachmann T, Singer W, Melloni L. Distilling the neural correlates of consciousness. Neurosci Biobehav Rev. 2012;36(2):737–46.

103. Pitts MA, Metzler S, Hillyard SA. Isolating neural correlates of conscious perception from neural correlates of reporting one’s perception. Front Psychol. 2014;5(1078).

