## Supplemental presentation 1 for "The neural activity of auditory conscious perception"

### Gamma power activity changes:

### Full timelines

**Supplementary Presentation S1. Perceived, Not Perceived, and Perceived – Not Perceived changes displayed at all time points.** Dynamic changes in cortical broad-band gamma power for threshold auditory stimuli as in Figures 3 and 4 but showing the full range of time points and surface views. No significant changes are seen in baseline time bins before target sound onset (0 ms), after which strikingly different changes are seen in between Perceived and Not Perceived trials. N = 31 participants.

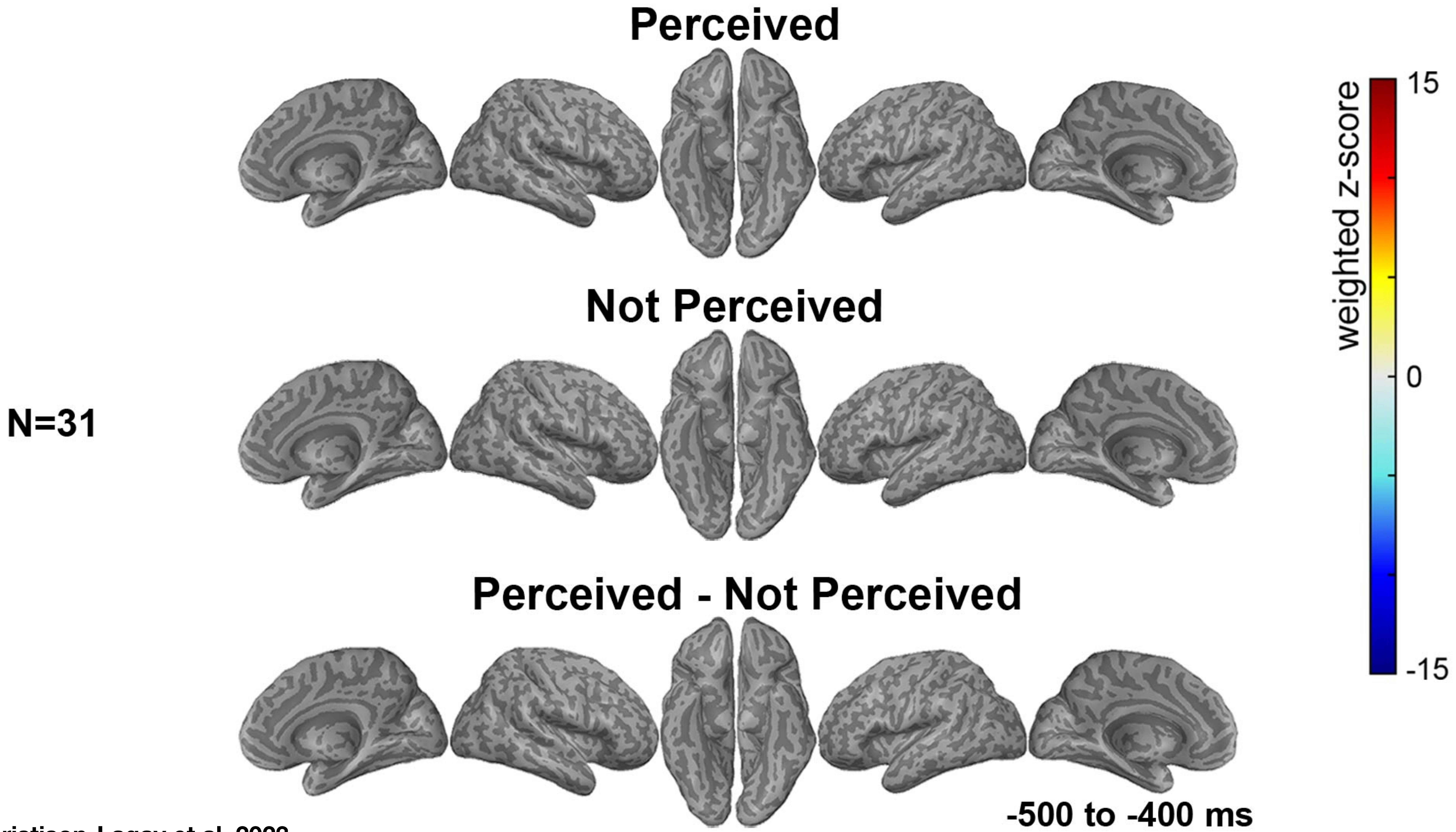

Christison-Lagay et al, 2023

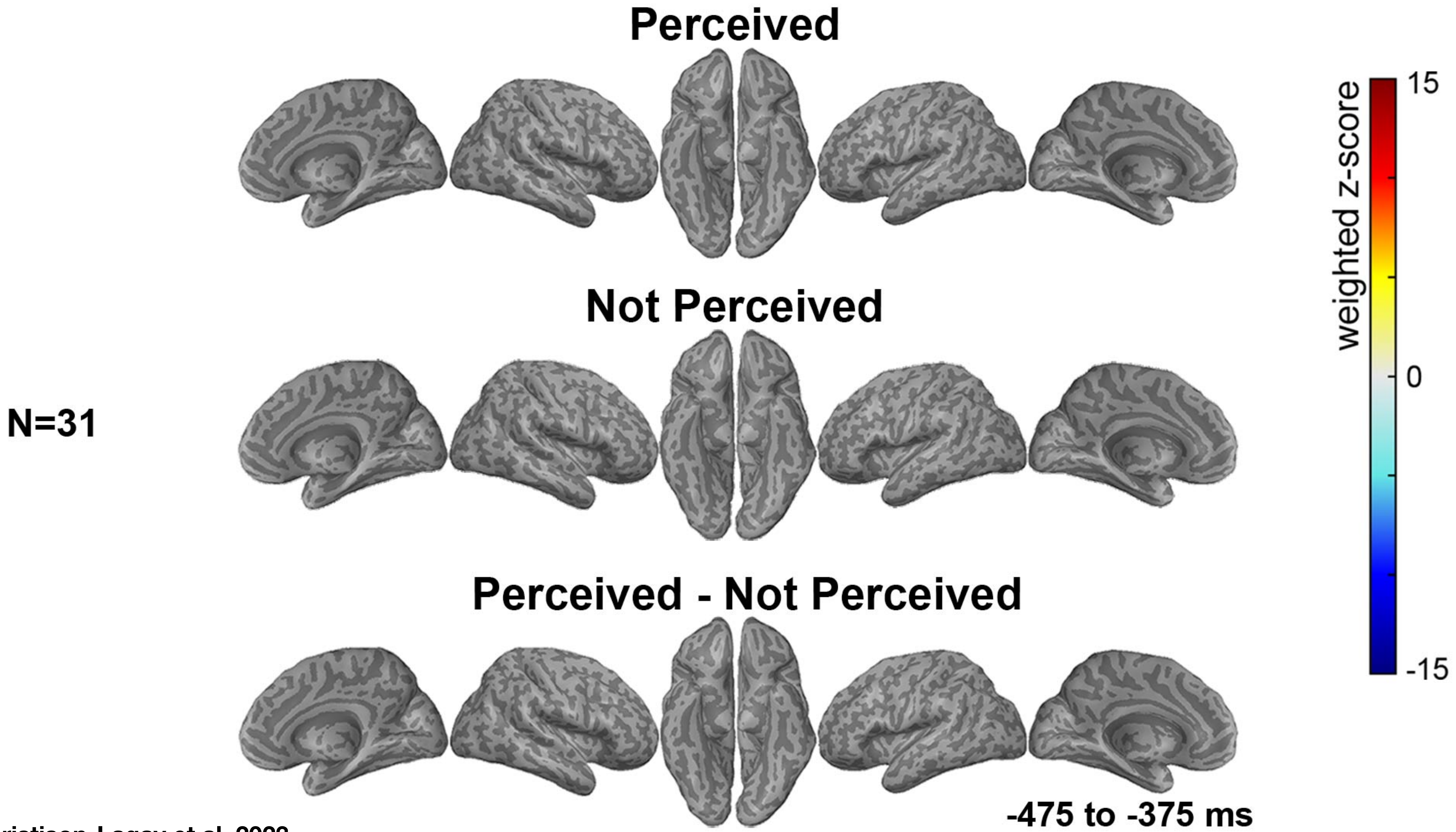


Christison-Lagay et al, 2023

N=31

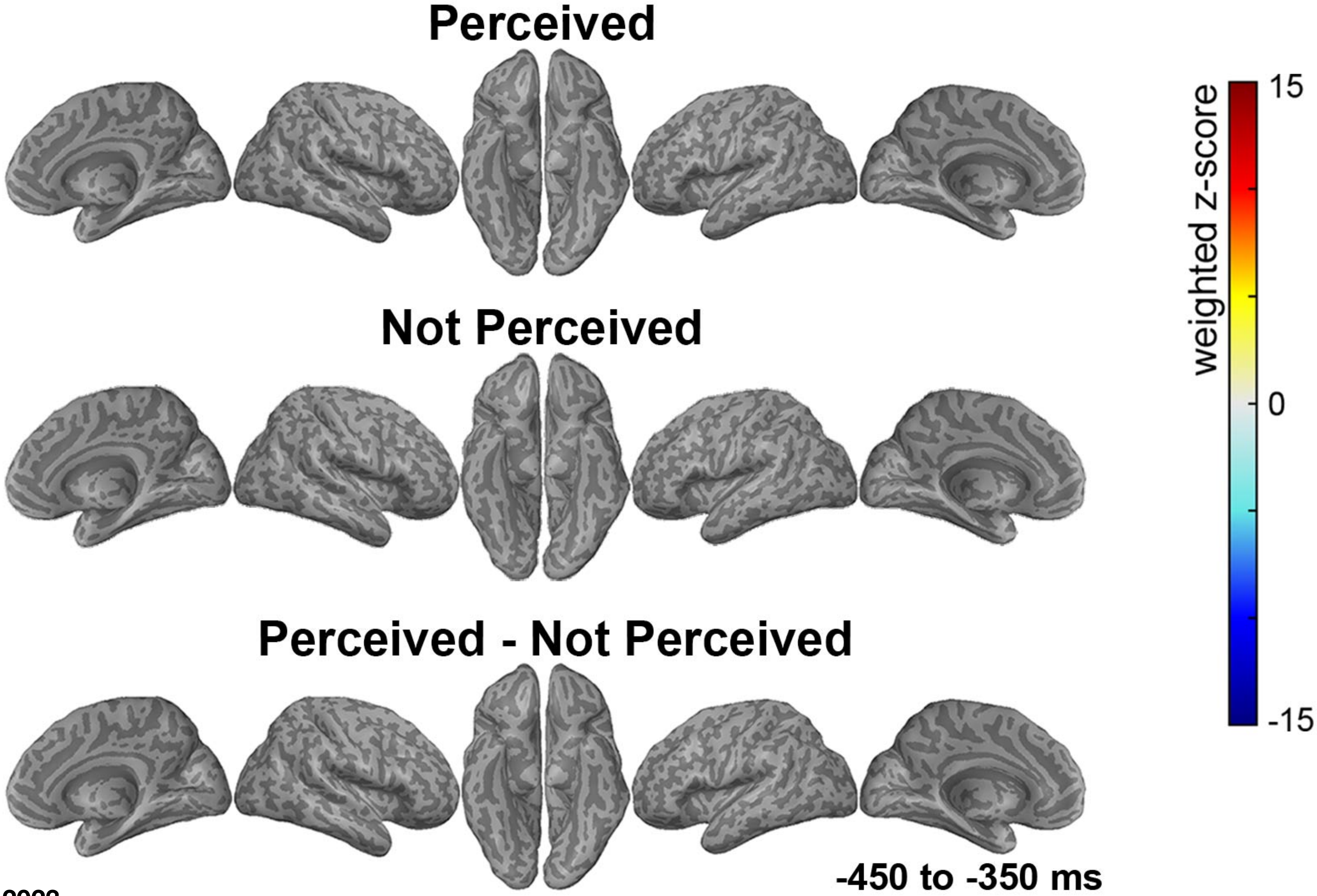


Christison-Lagay et al, 2023

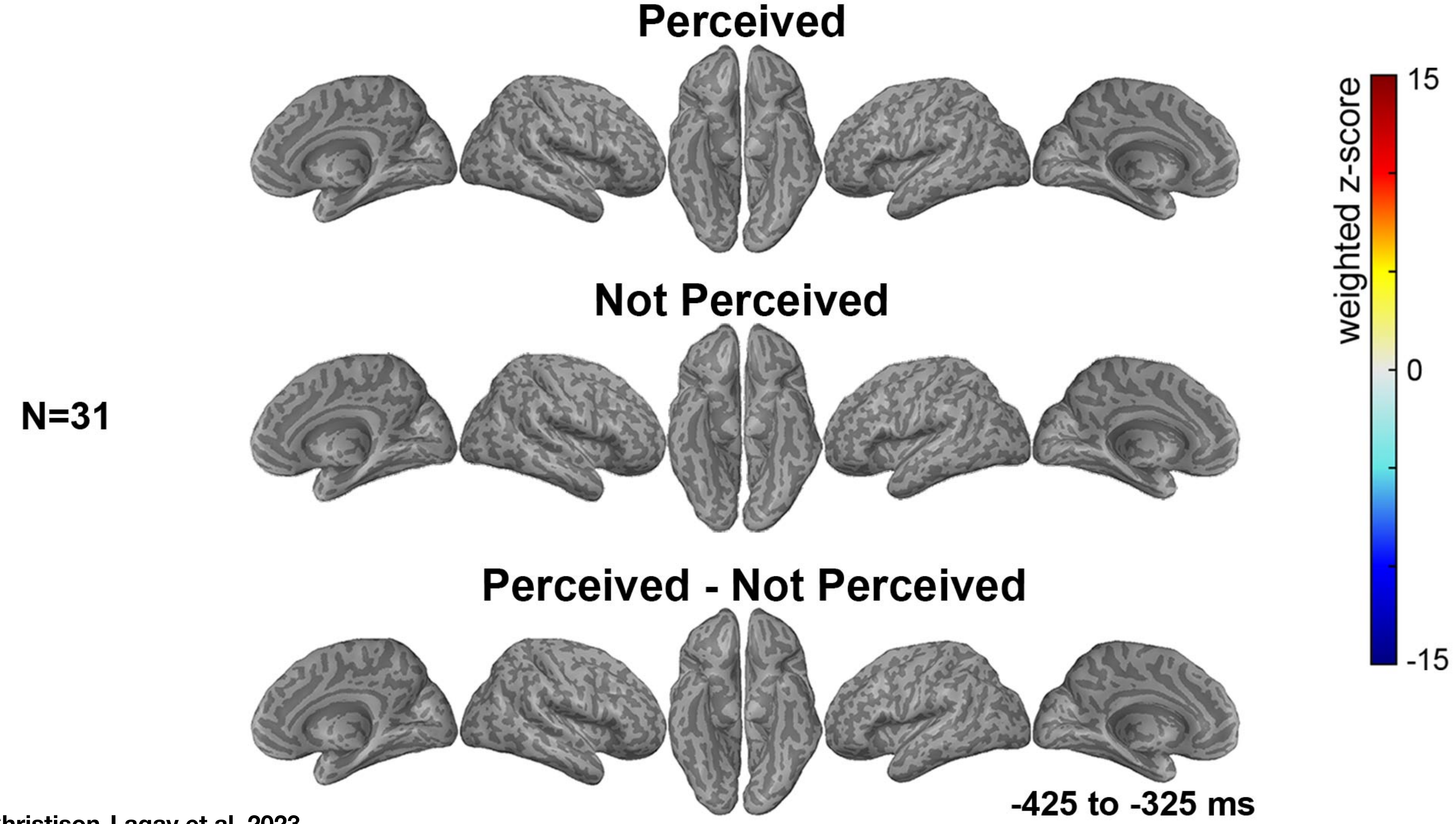

Christison-Lagay et al, 2023

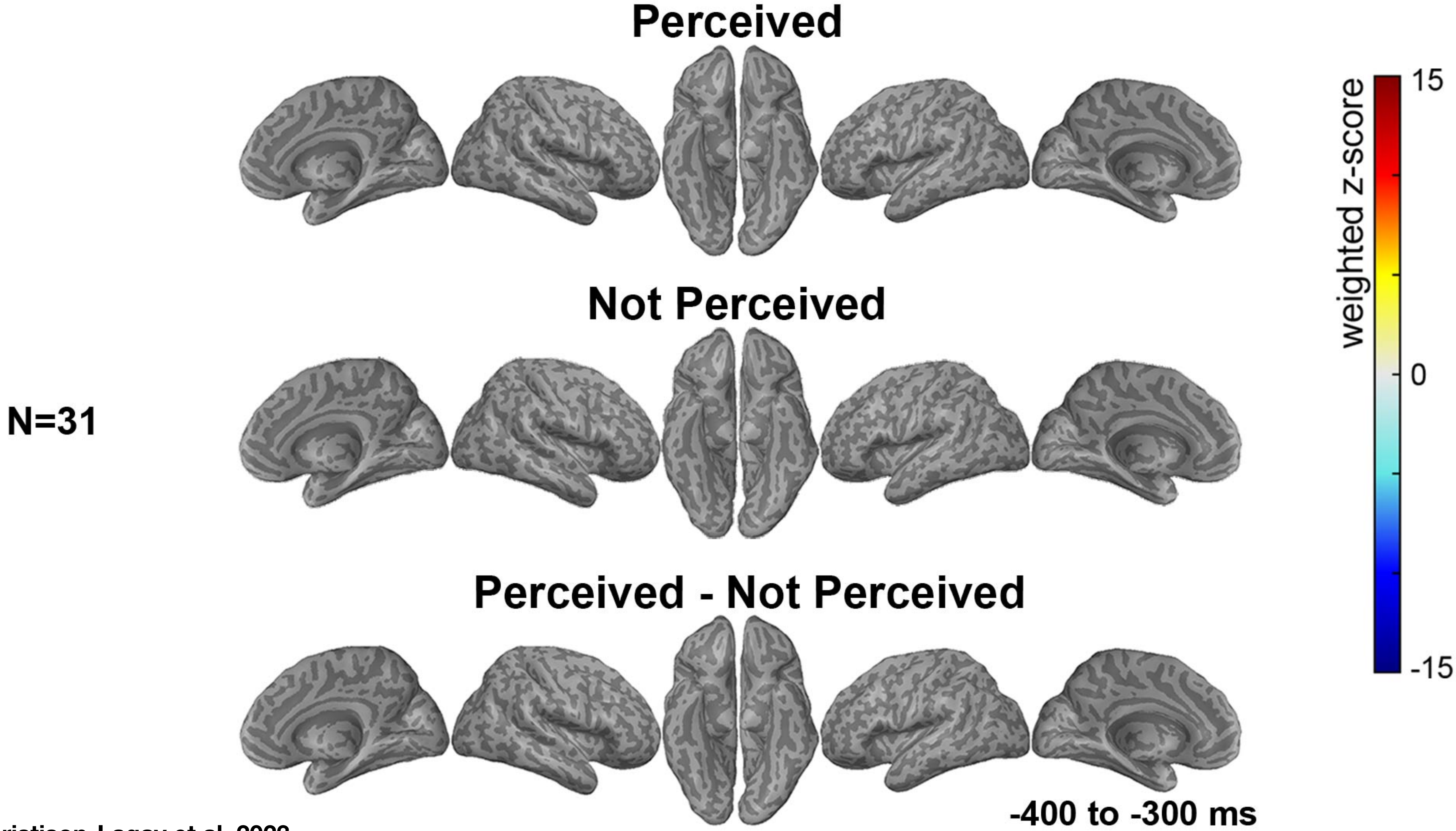

Christison-Lagay et al, 2023

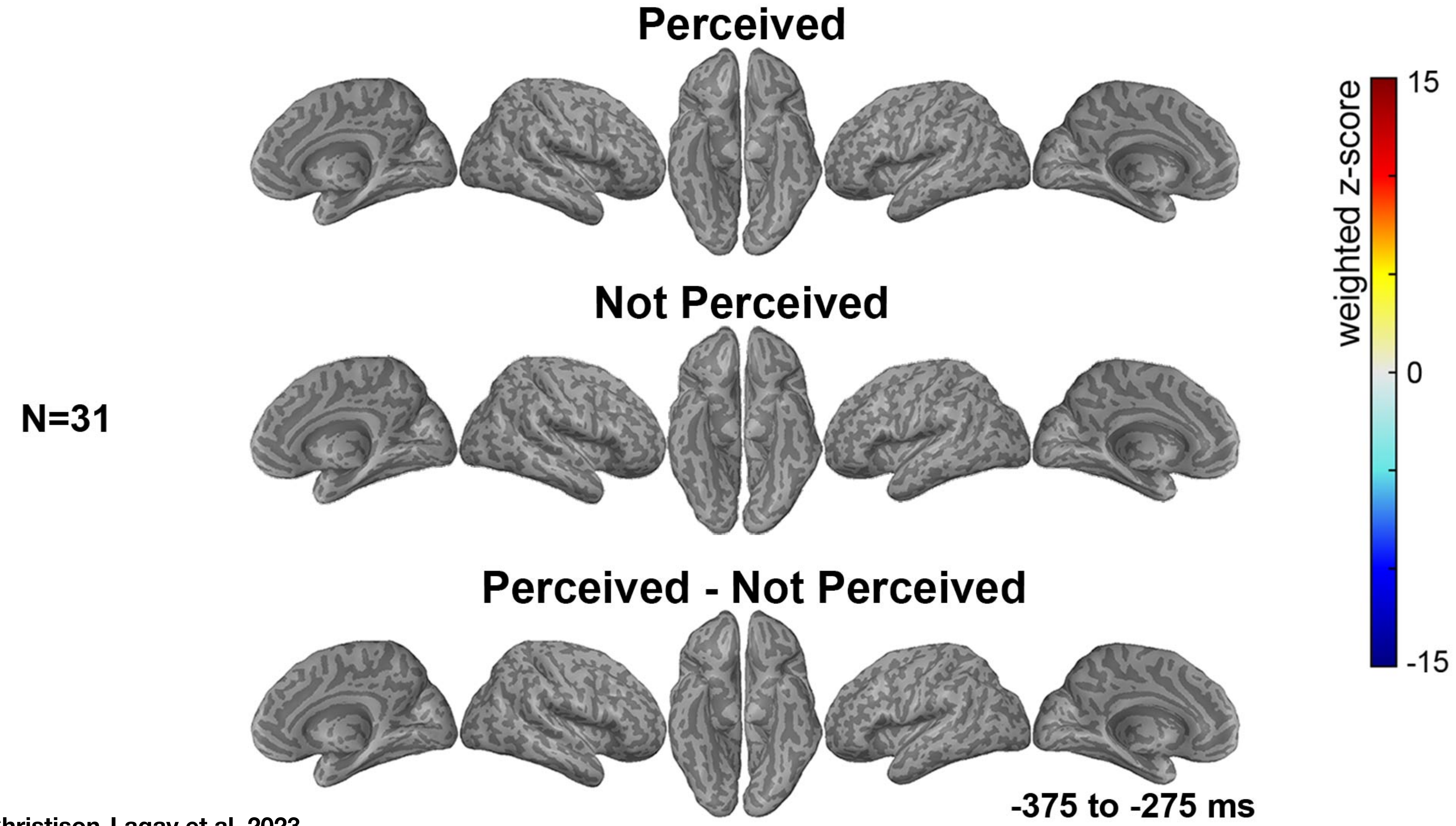


Christison-Lagay et al, 2023

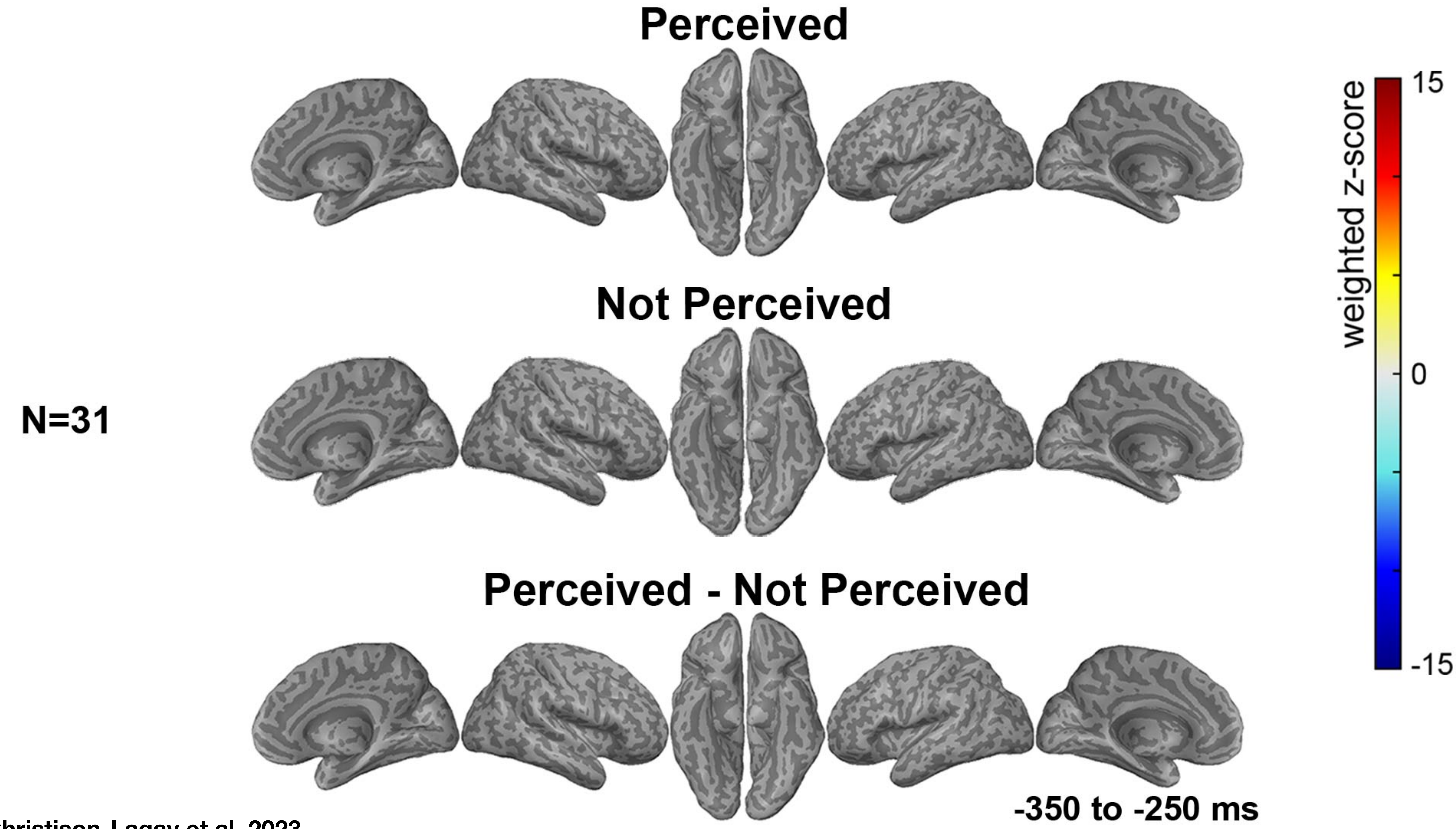


Christison-Lagay et al, 2023

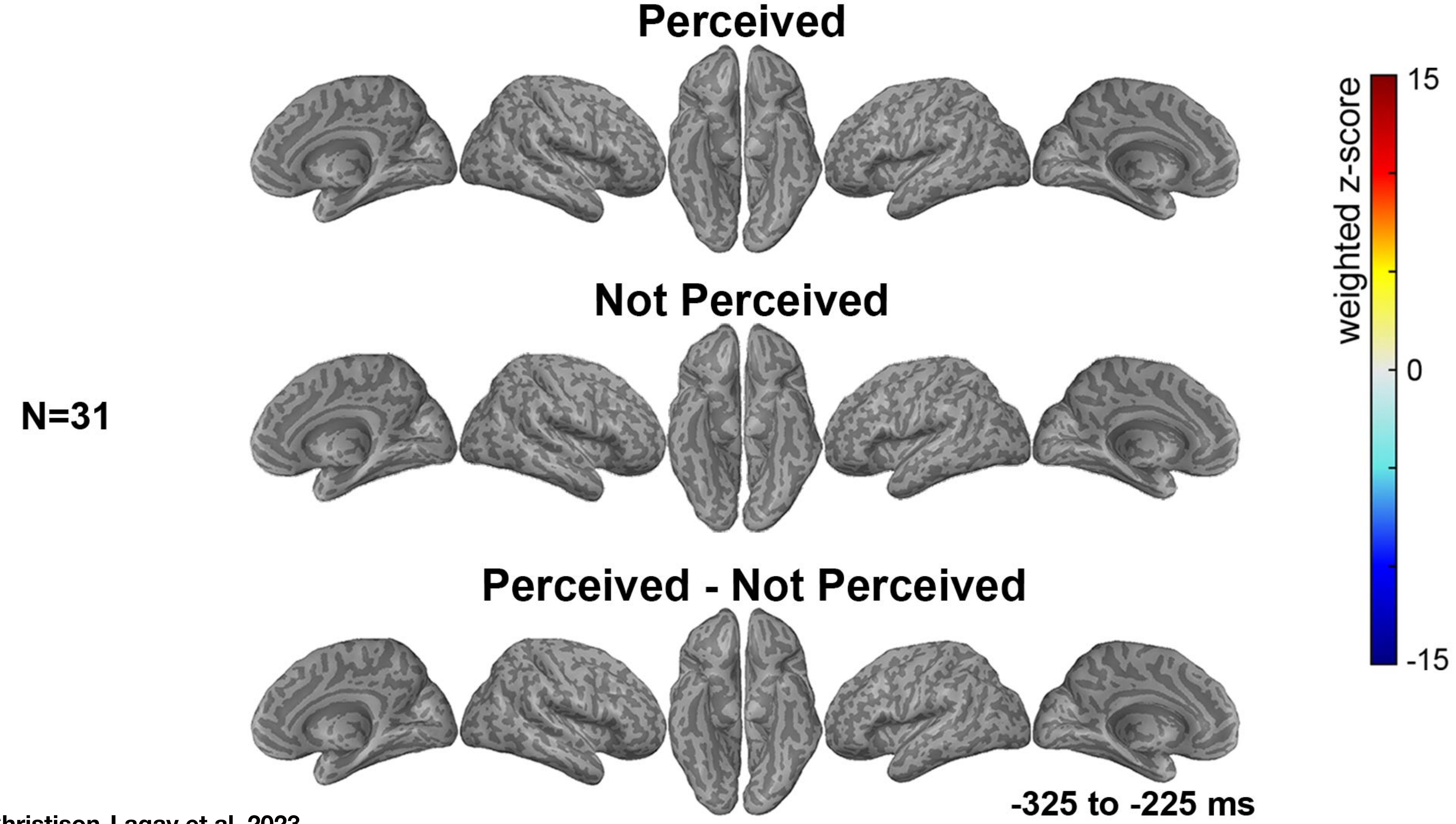


Christison-Lagay et al, 2023

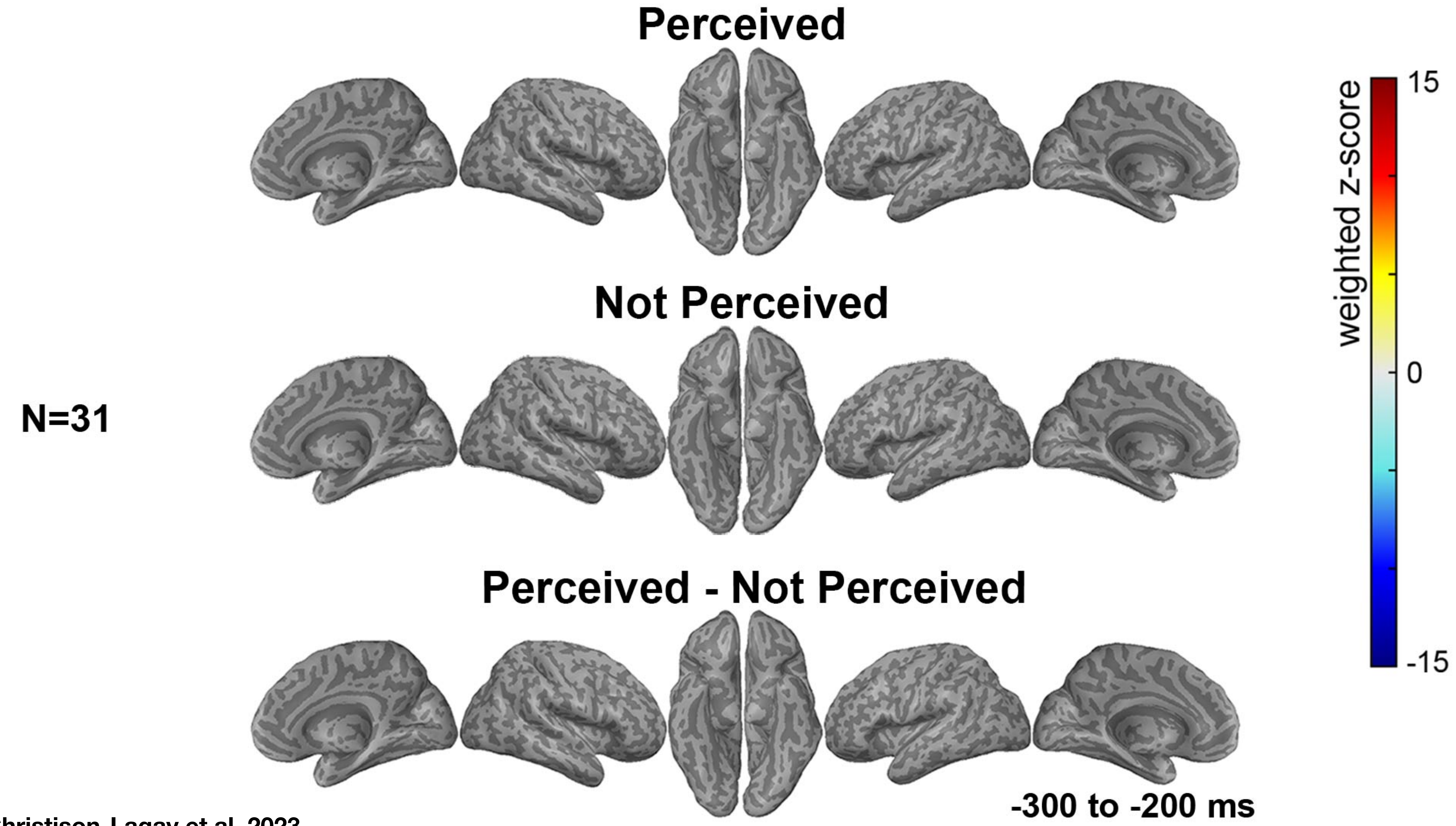


Christison-Lagay et al, 2023

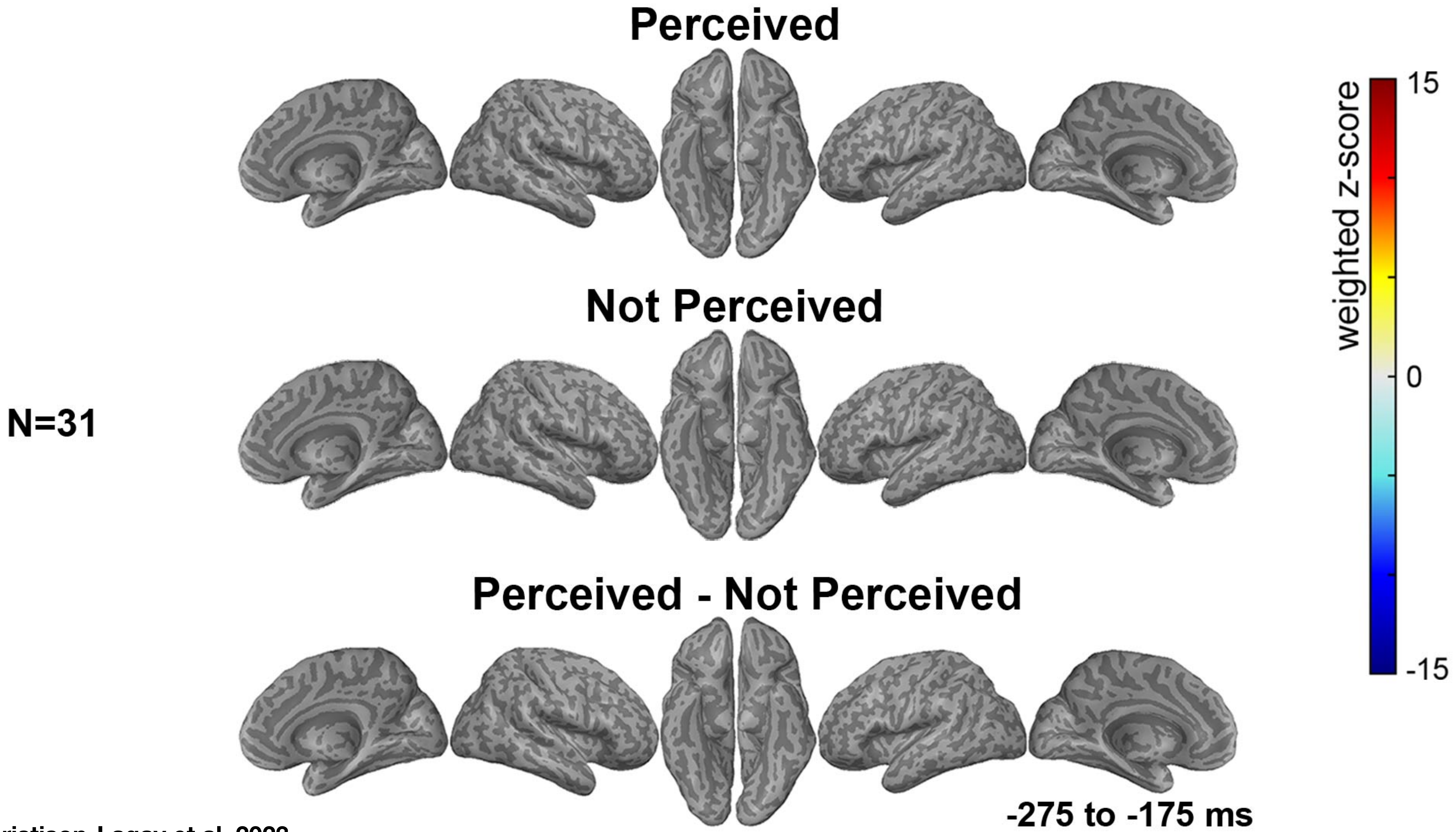


Christison-Lagay et al, 2023

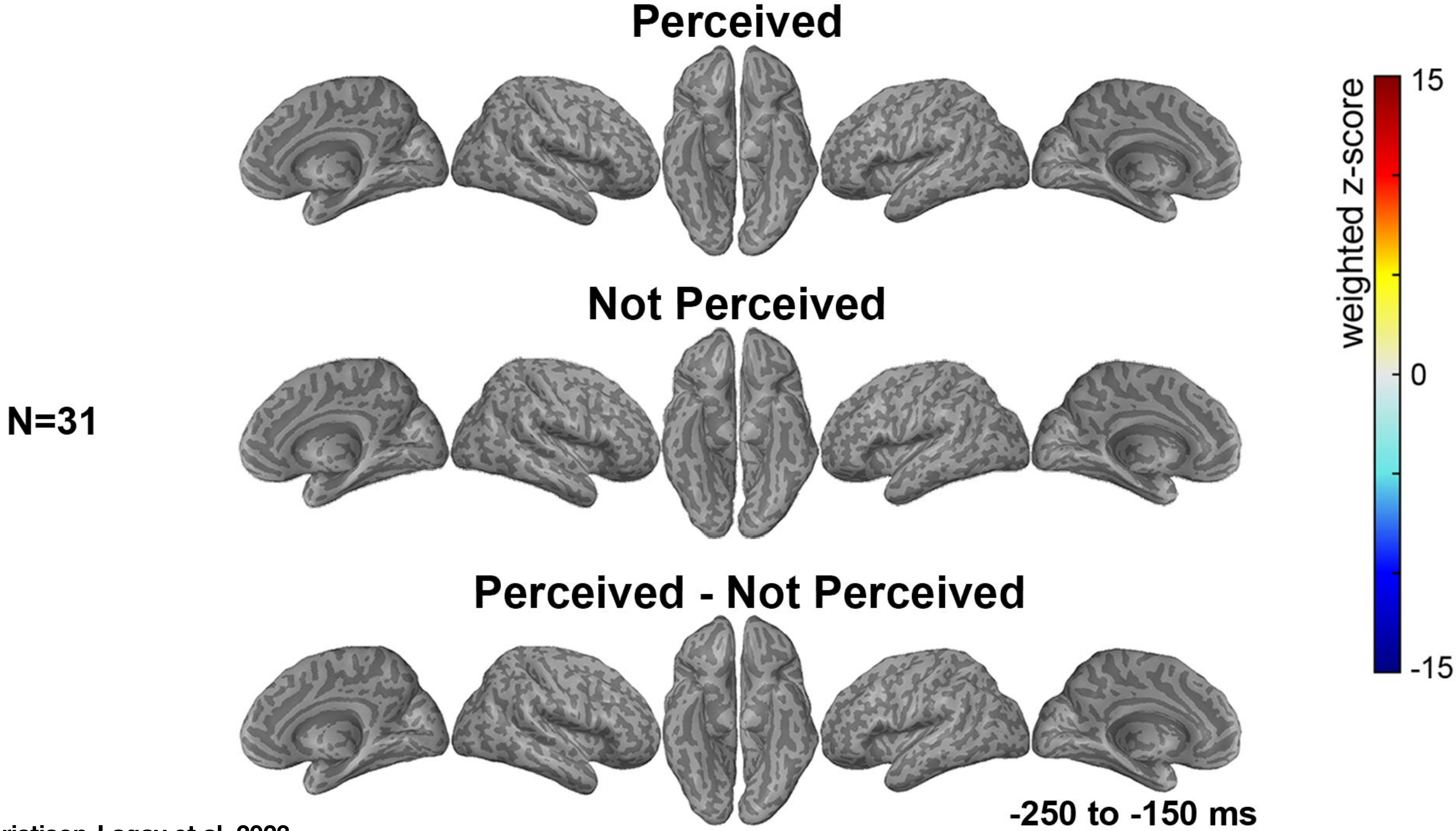


Christison-Lagay et al, 2023

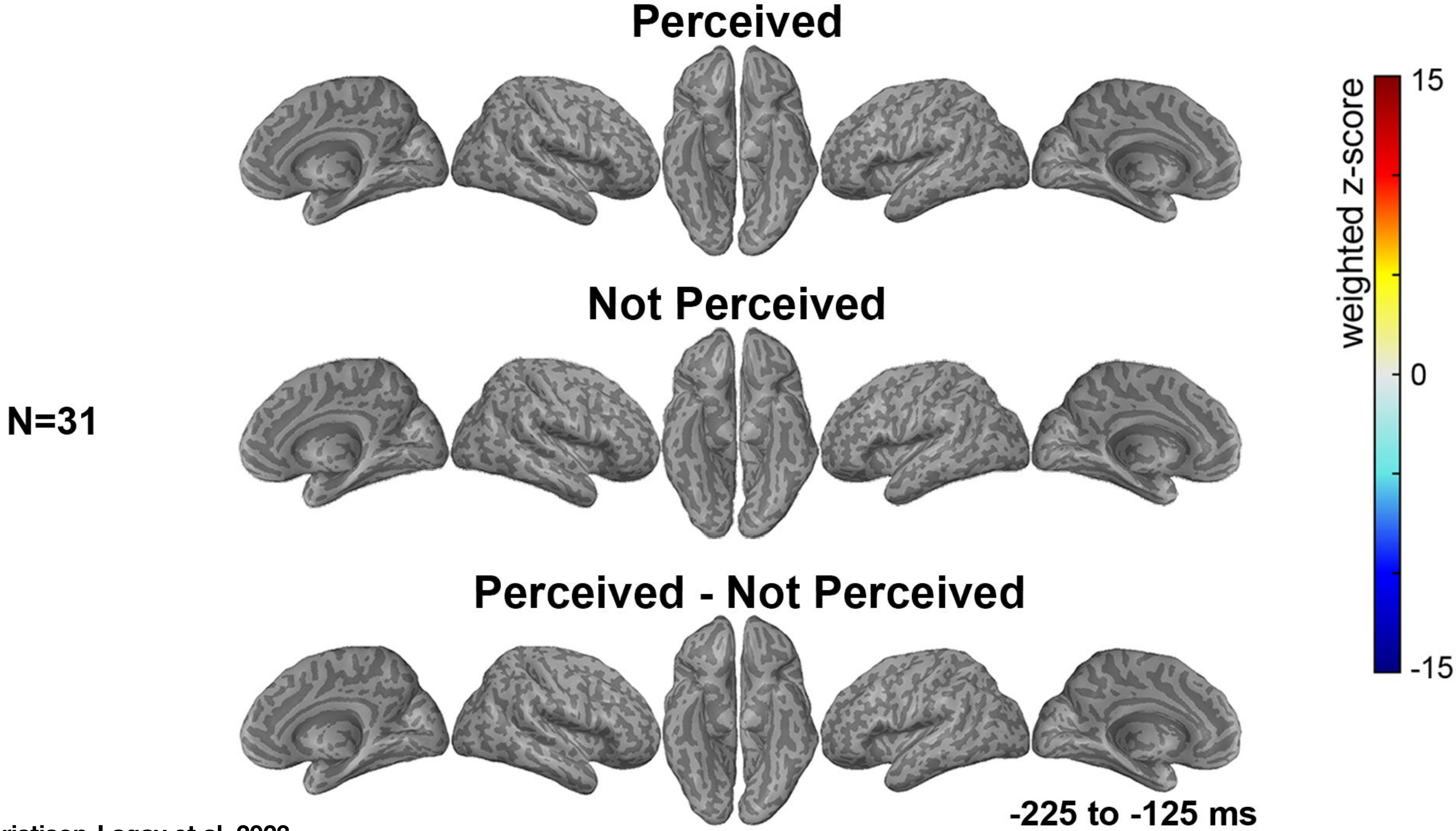


Christison-Lagay et al, 2023

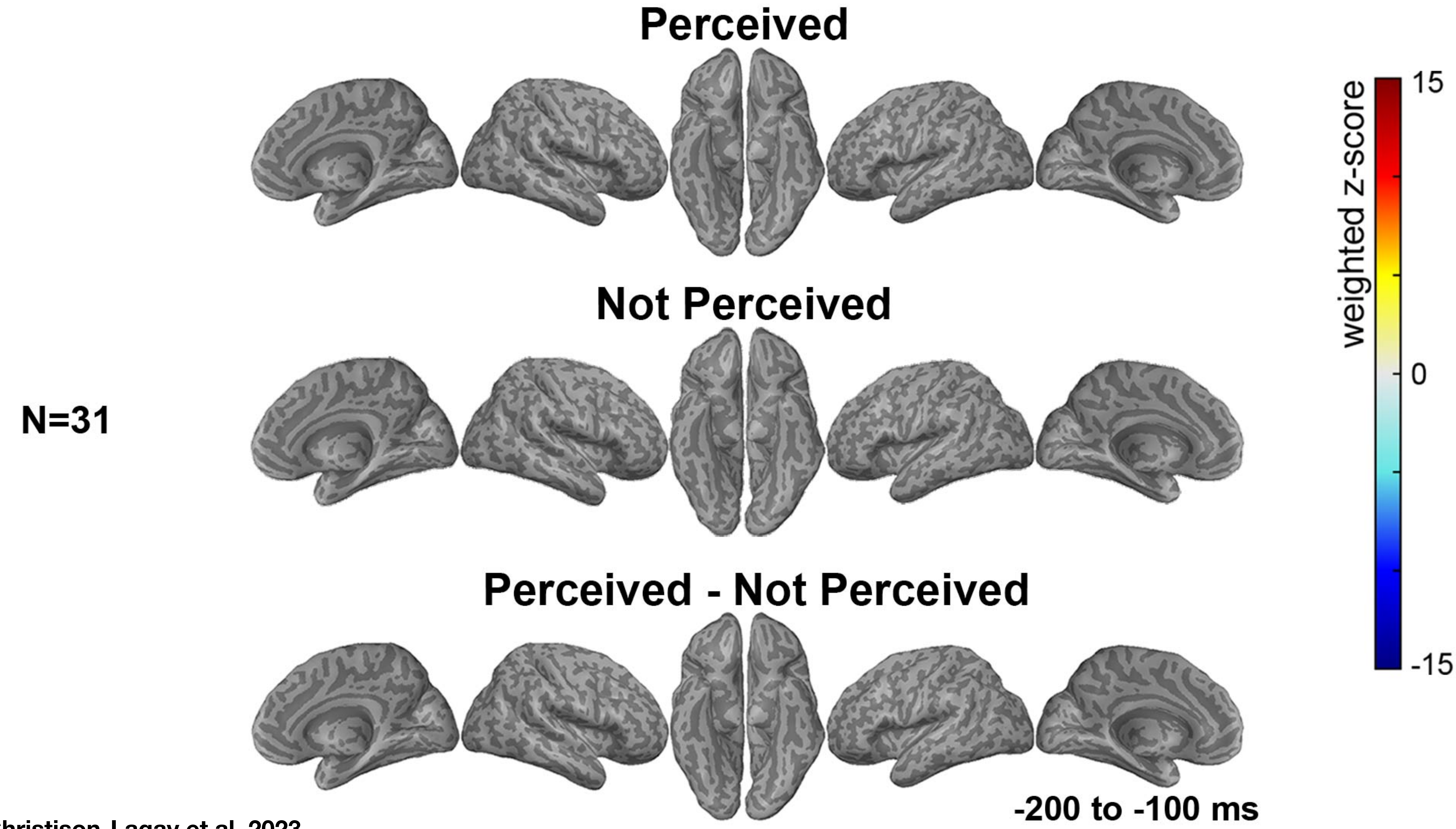


Christison-Lagay et al, 2023

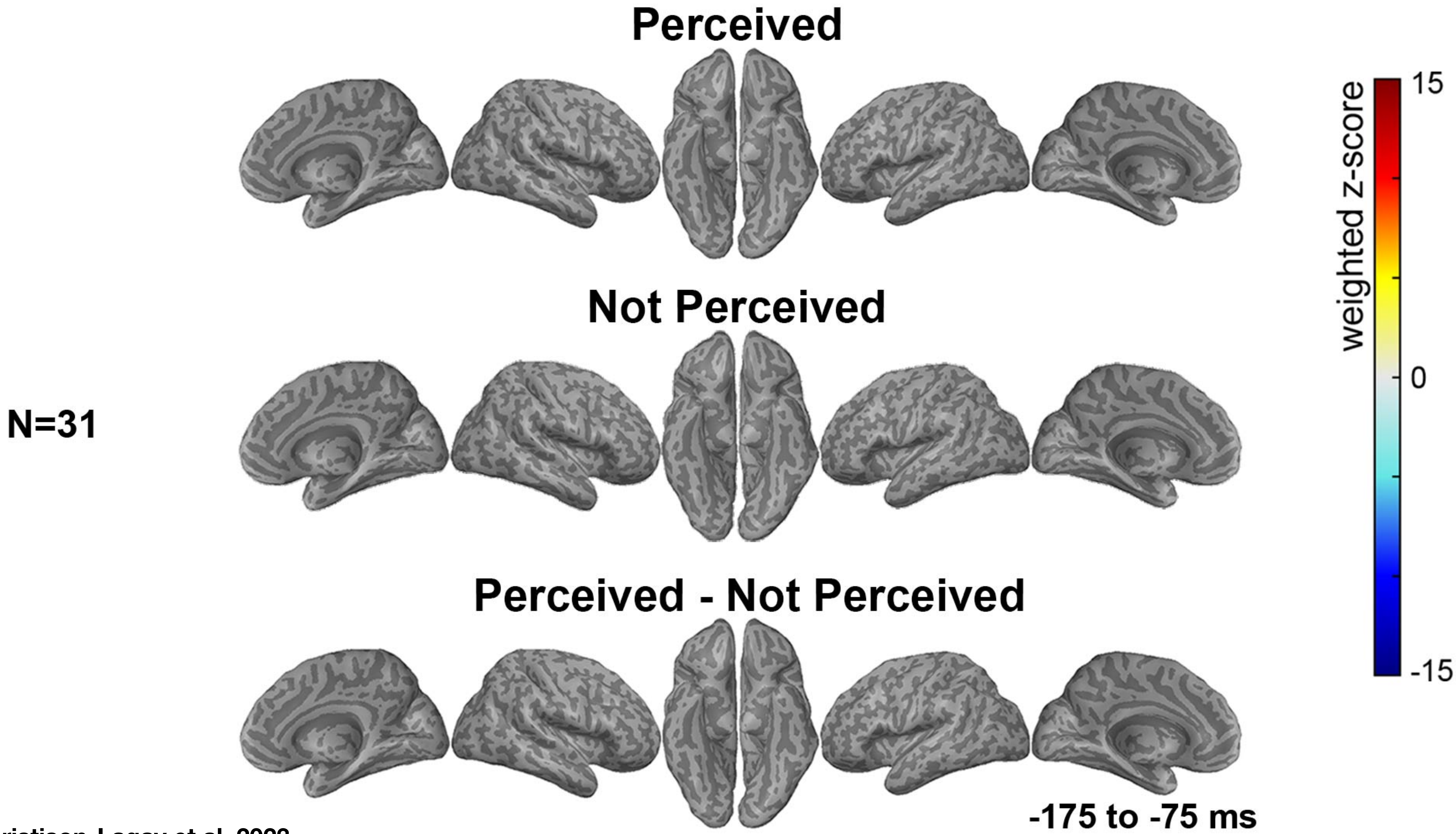


Christison-Lagay et al, 2023

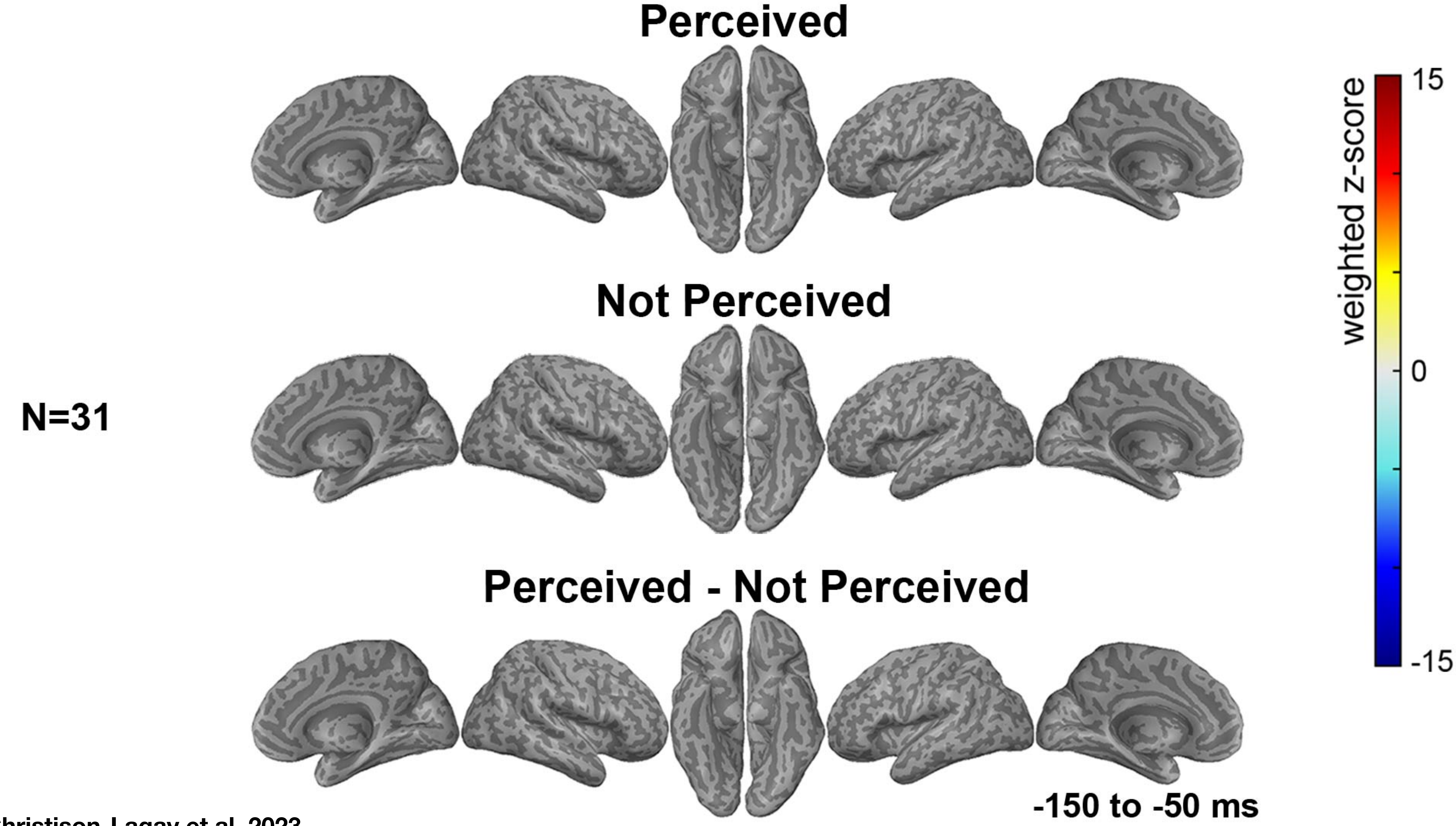

Christison-Lagay et al, 2023

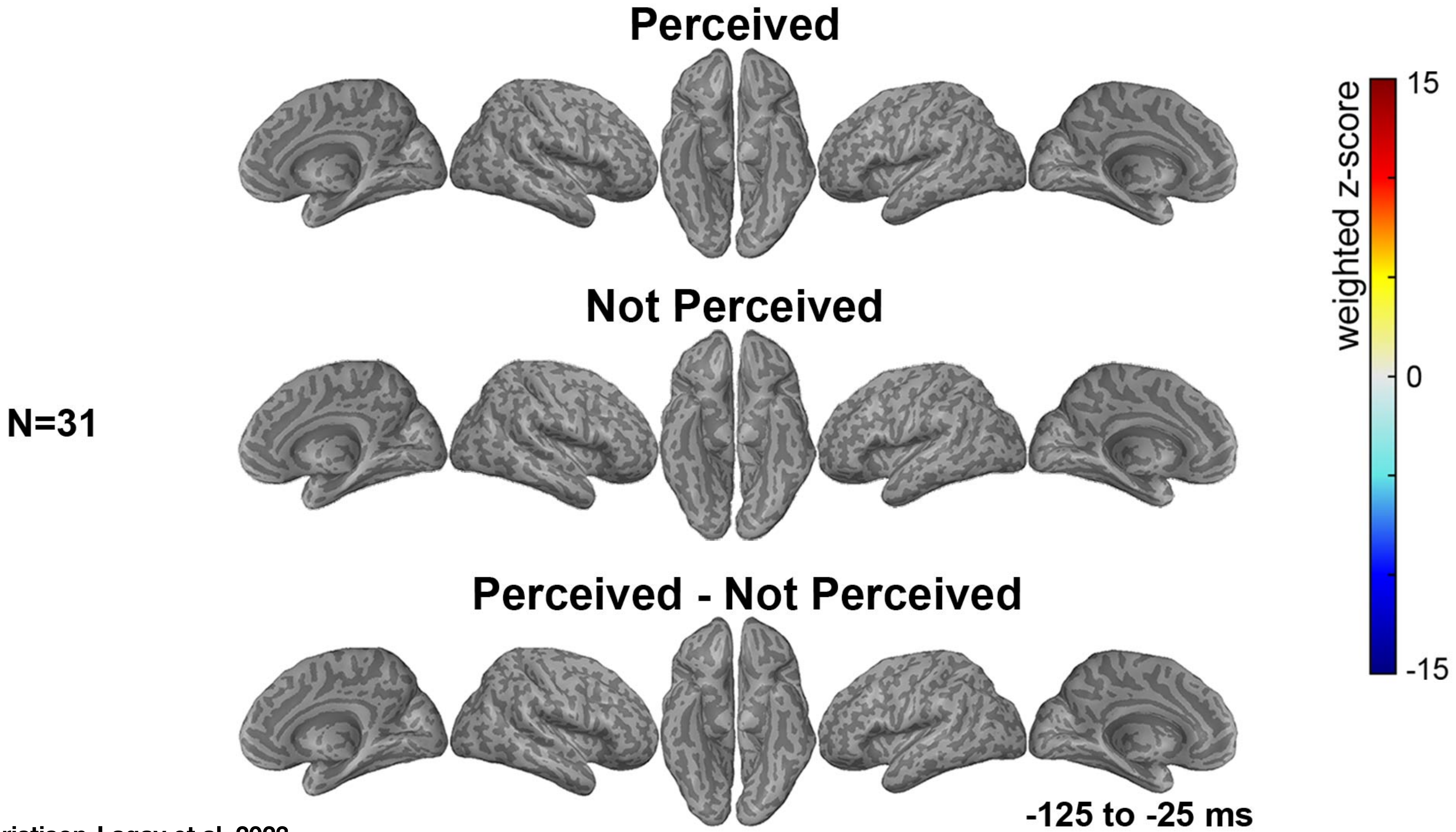


Christison-Lagay et al, 2023

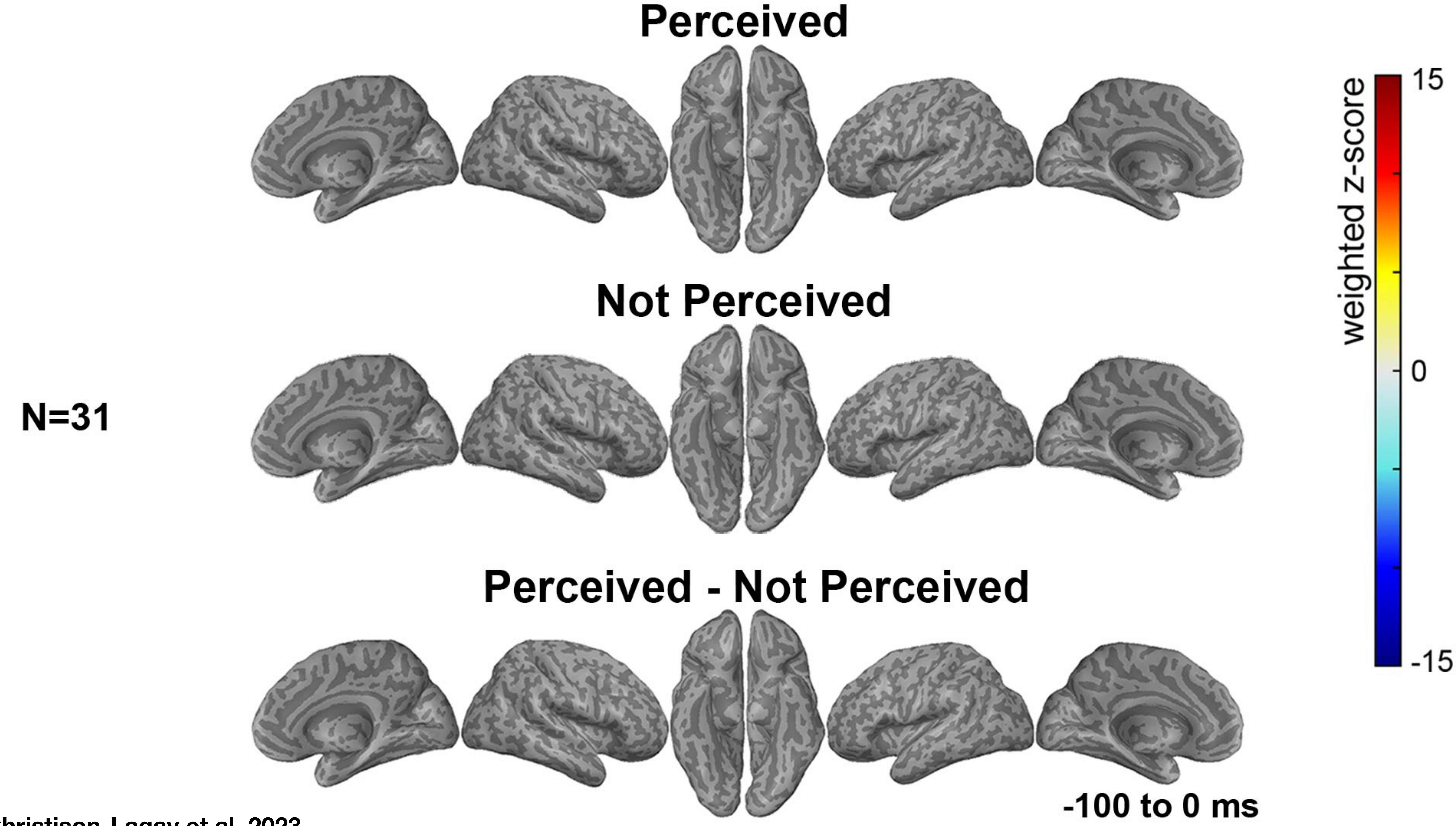


Christison-Lagay et al, 2023

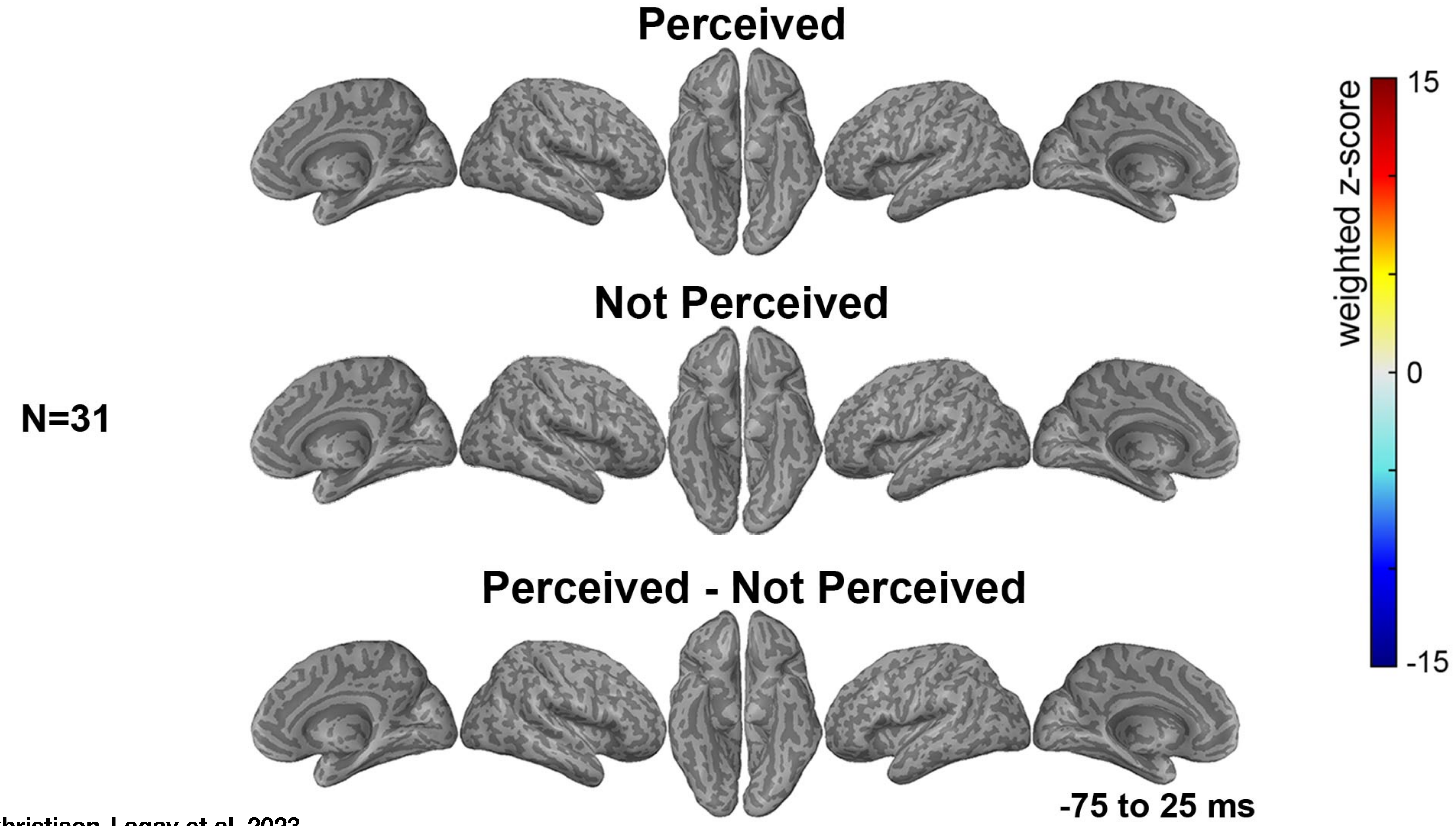


Christison-Lagay et al, 2023

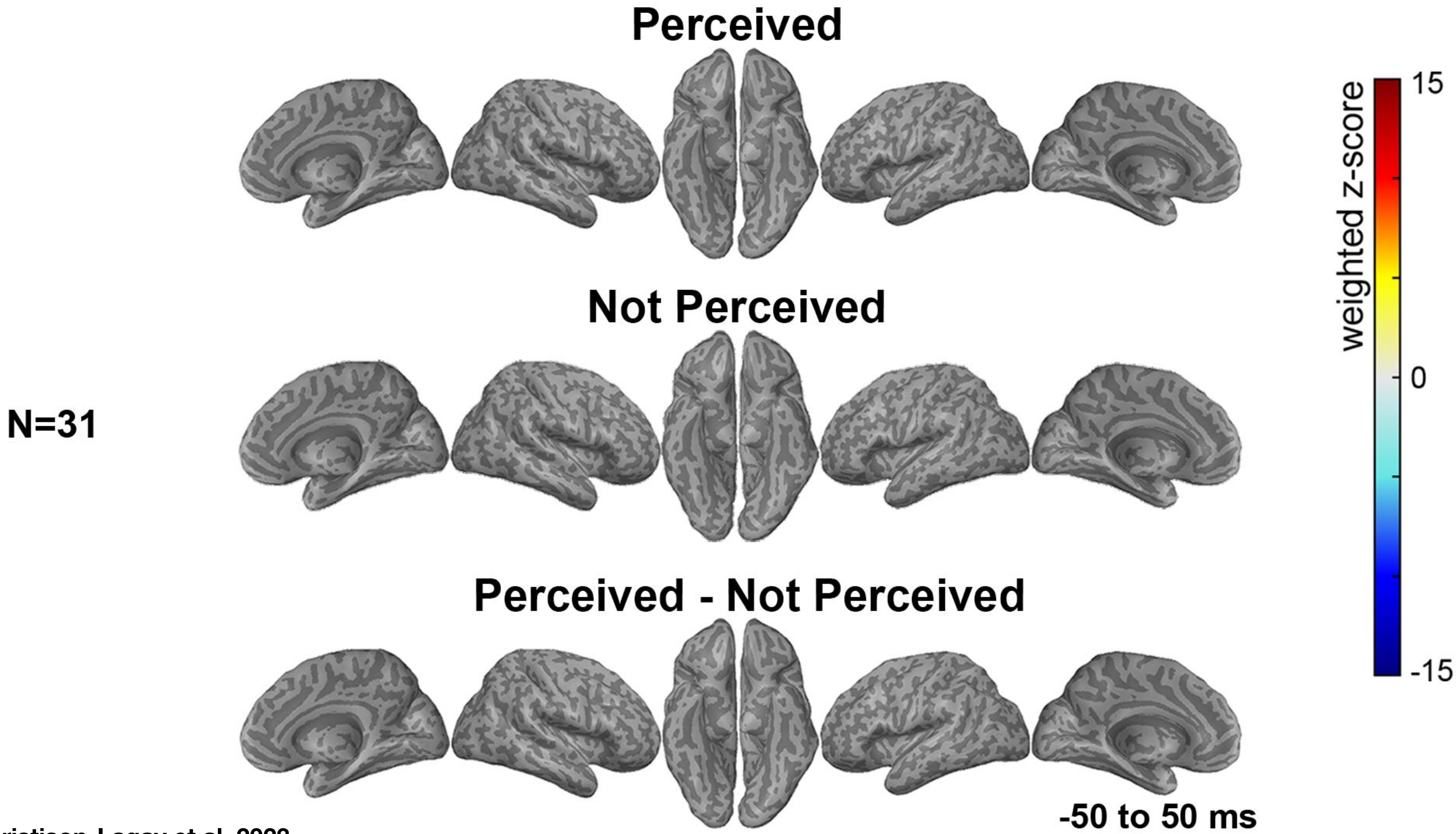


Christison-Lagay et al, 2023

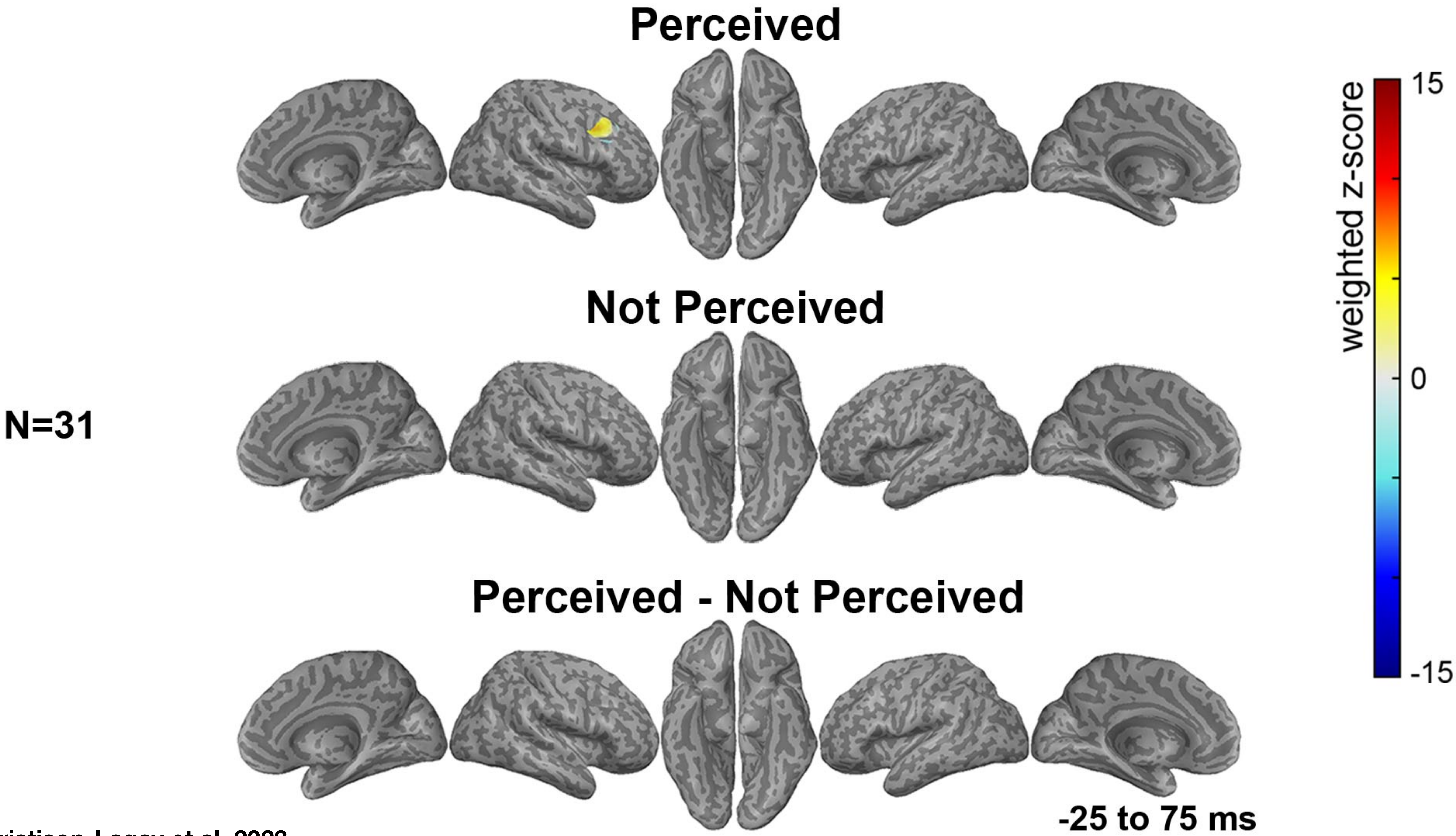


Christison-Lagay et al, 2023

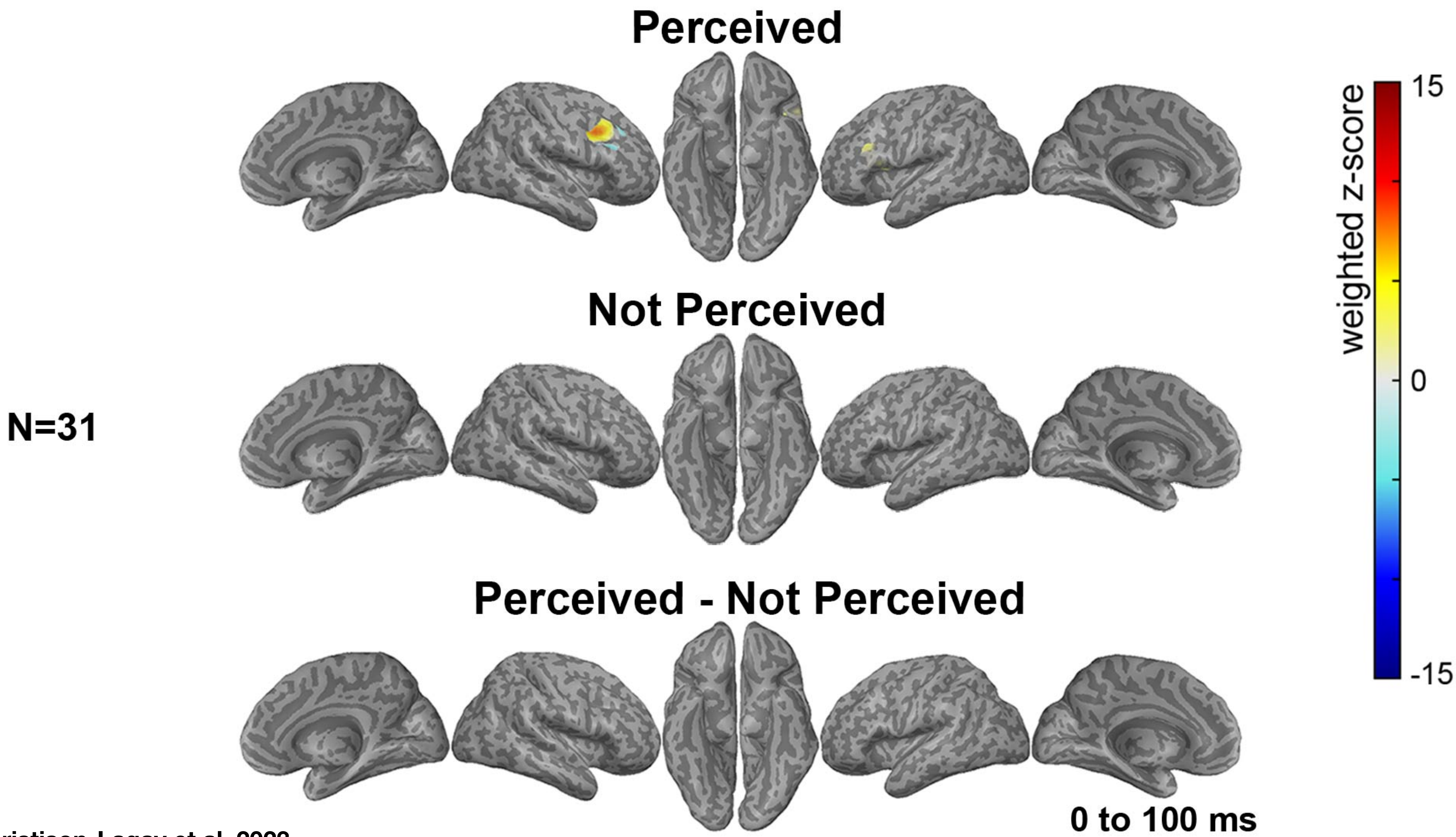


Christison-Lagay et al, 2023

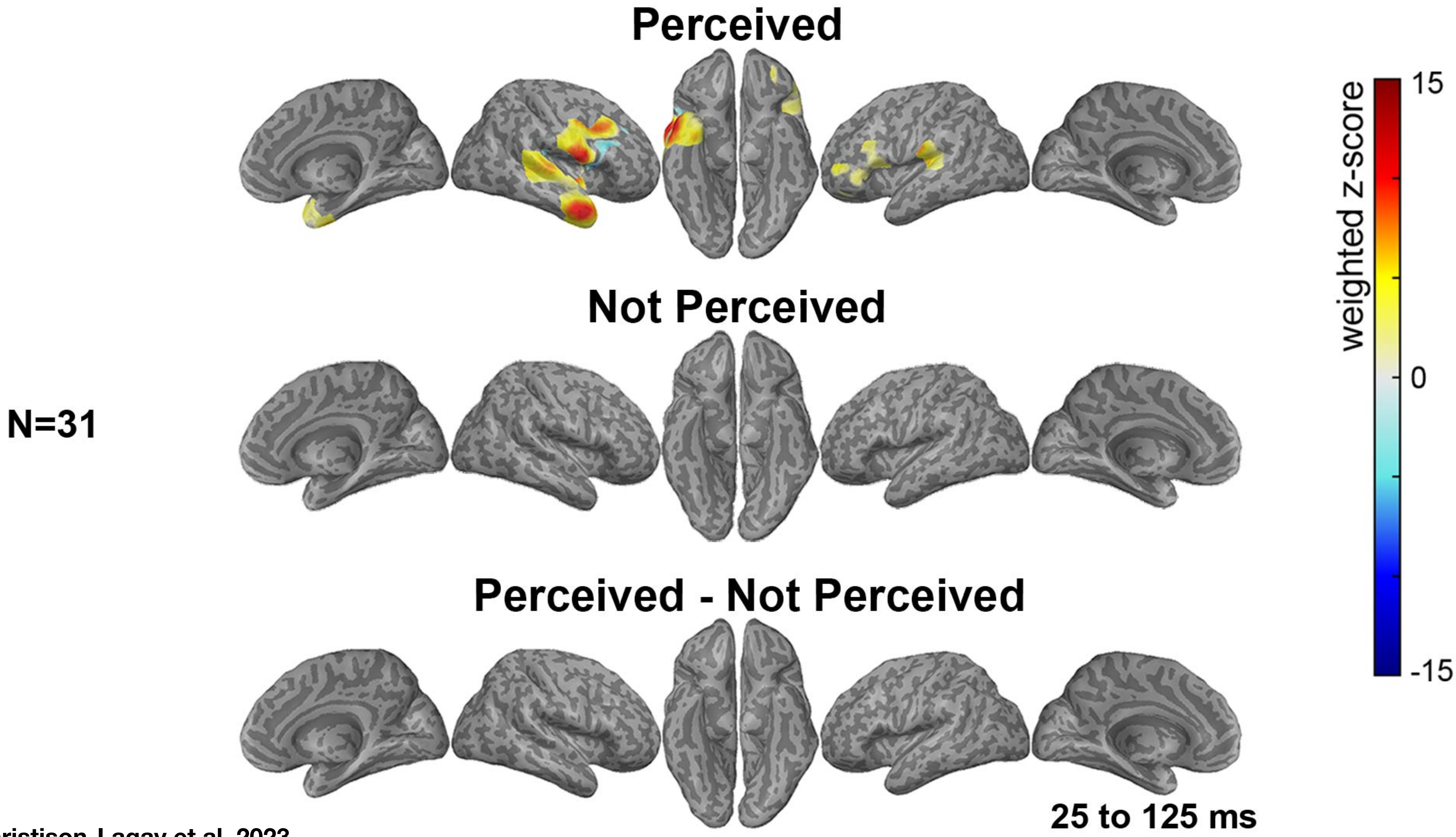


Christison-Lagay et al, 2023

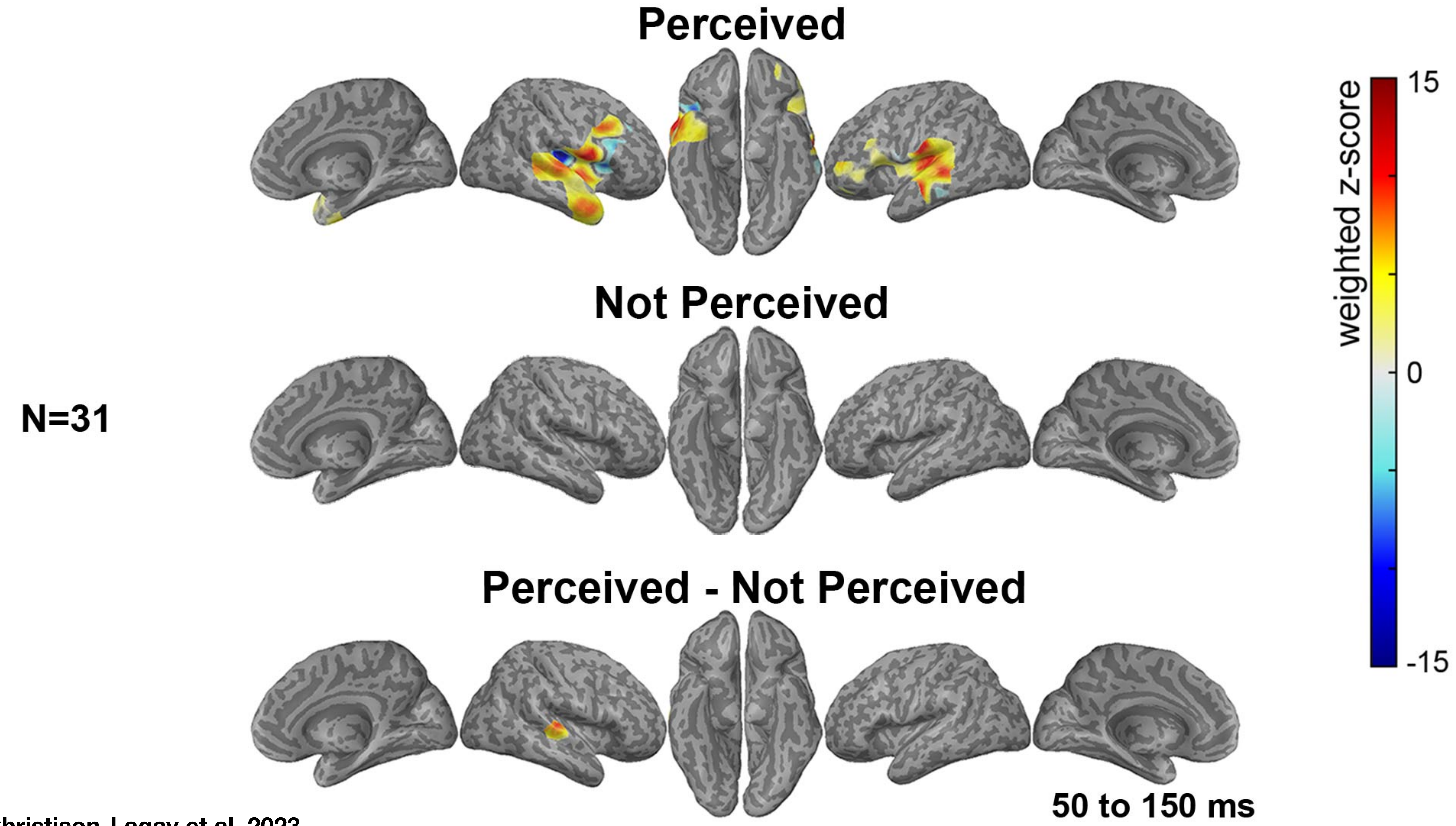


Christison-Lagay et al, 2023

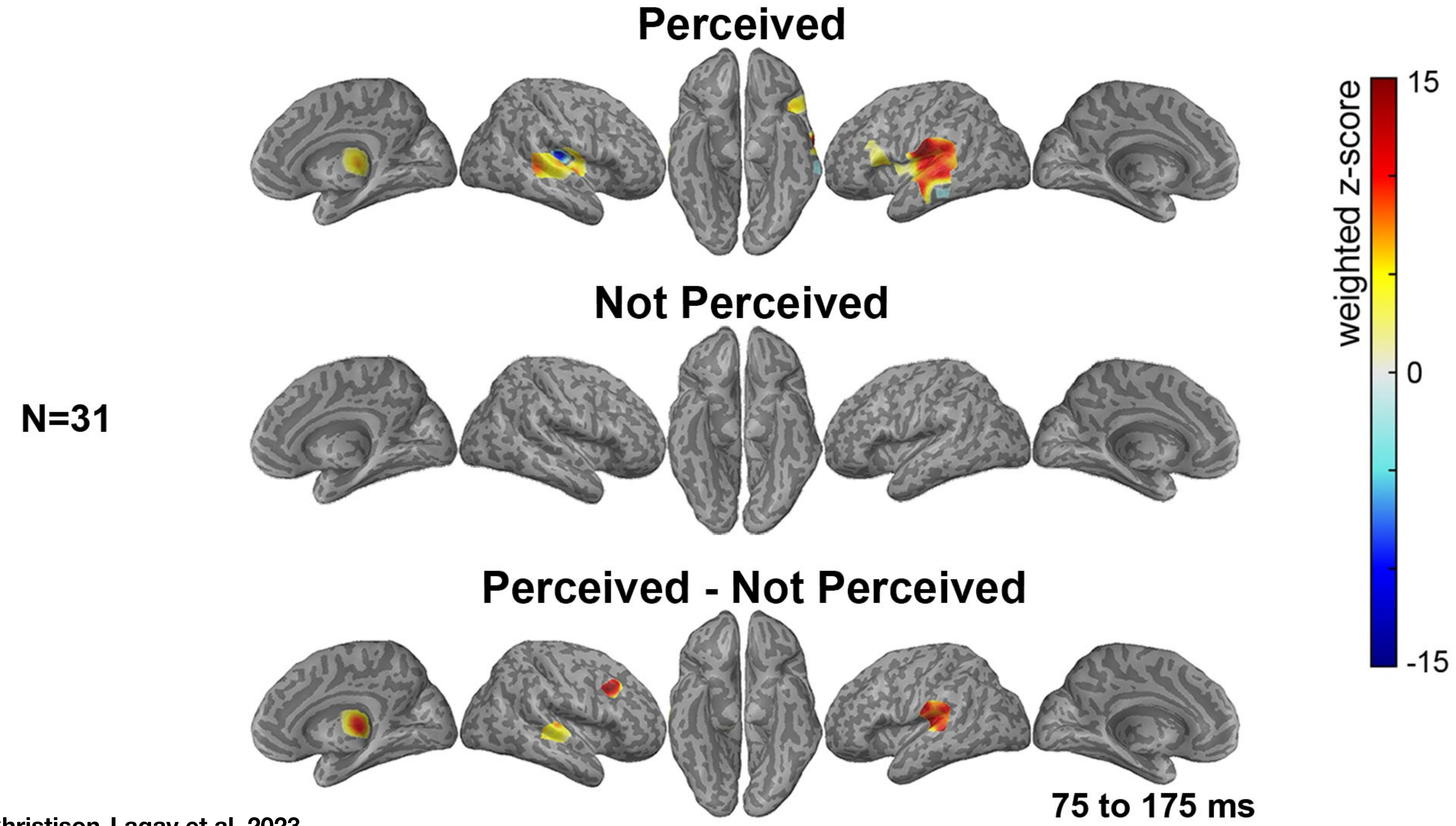


Christison-Lagay et al, 2023

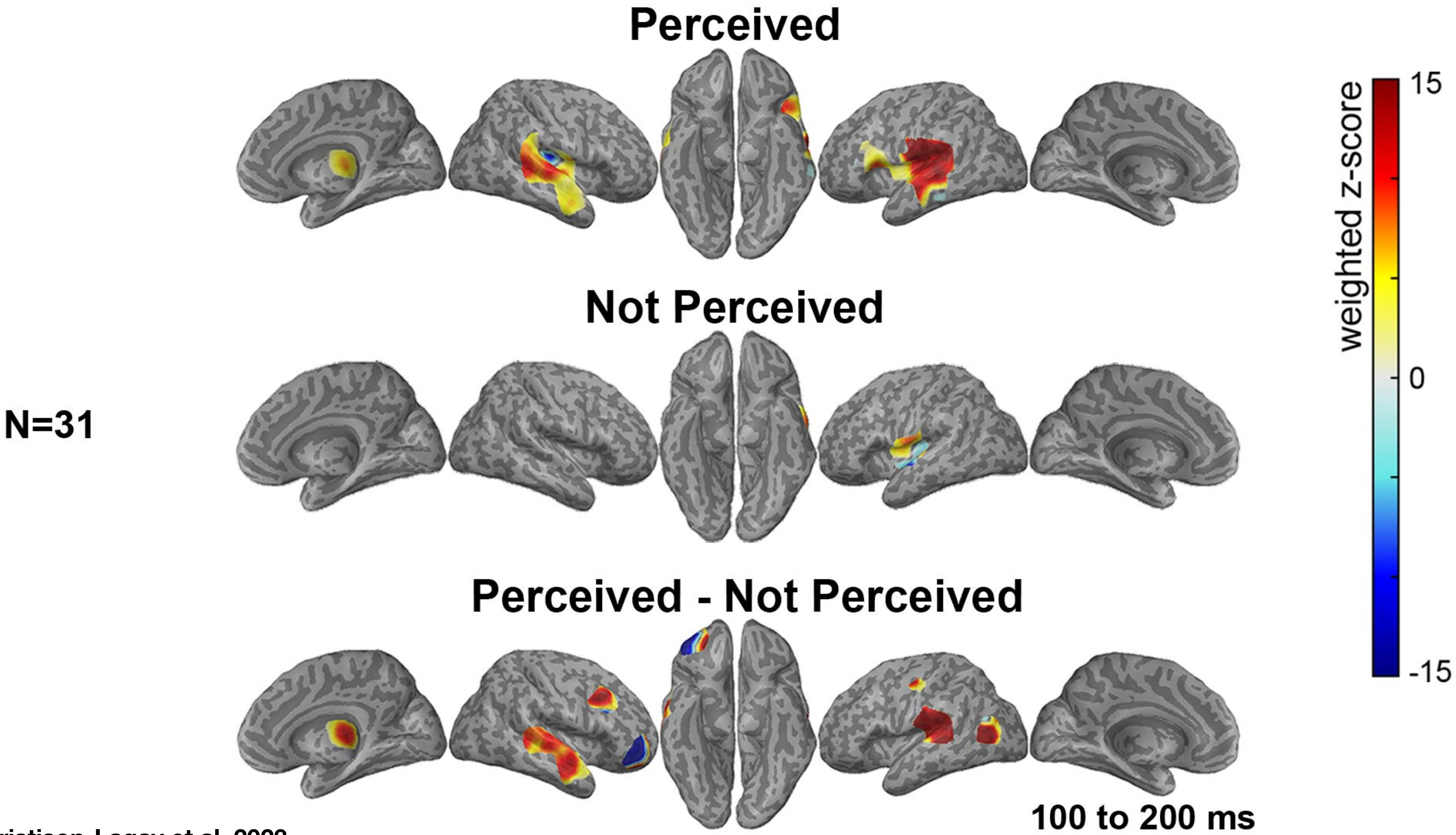


Christison-Lagay et al, 2023

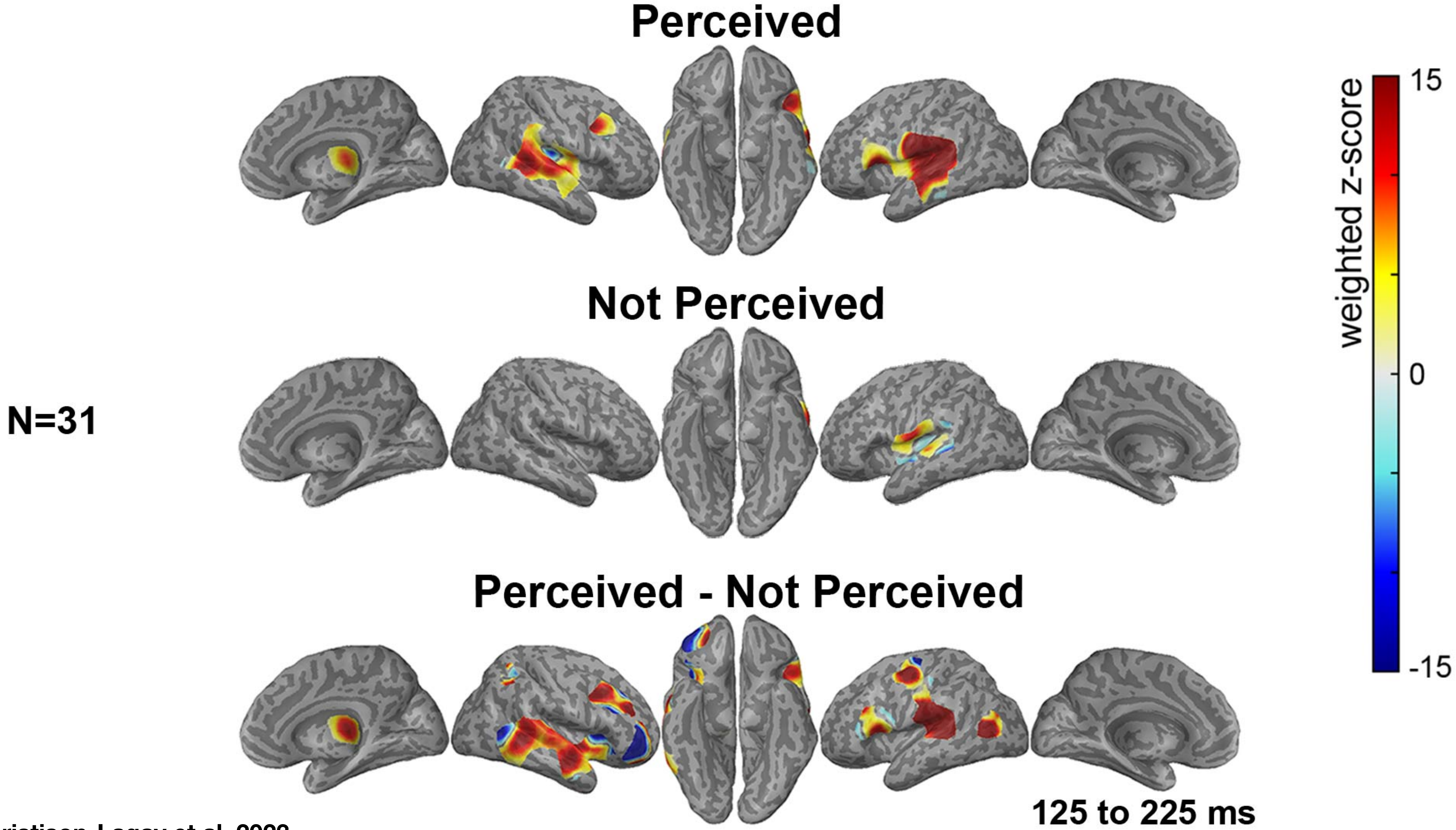


Christison-Lagay et al, 2023

N=31

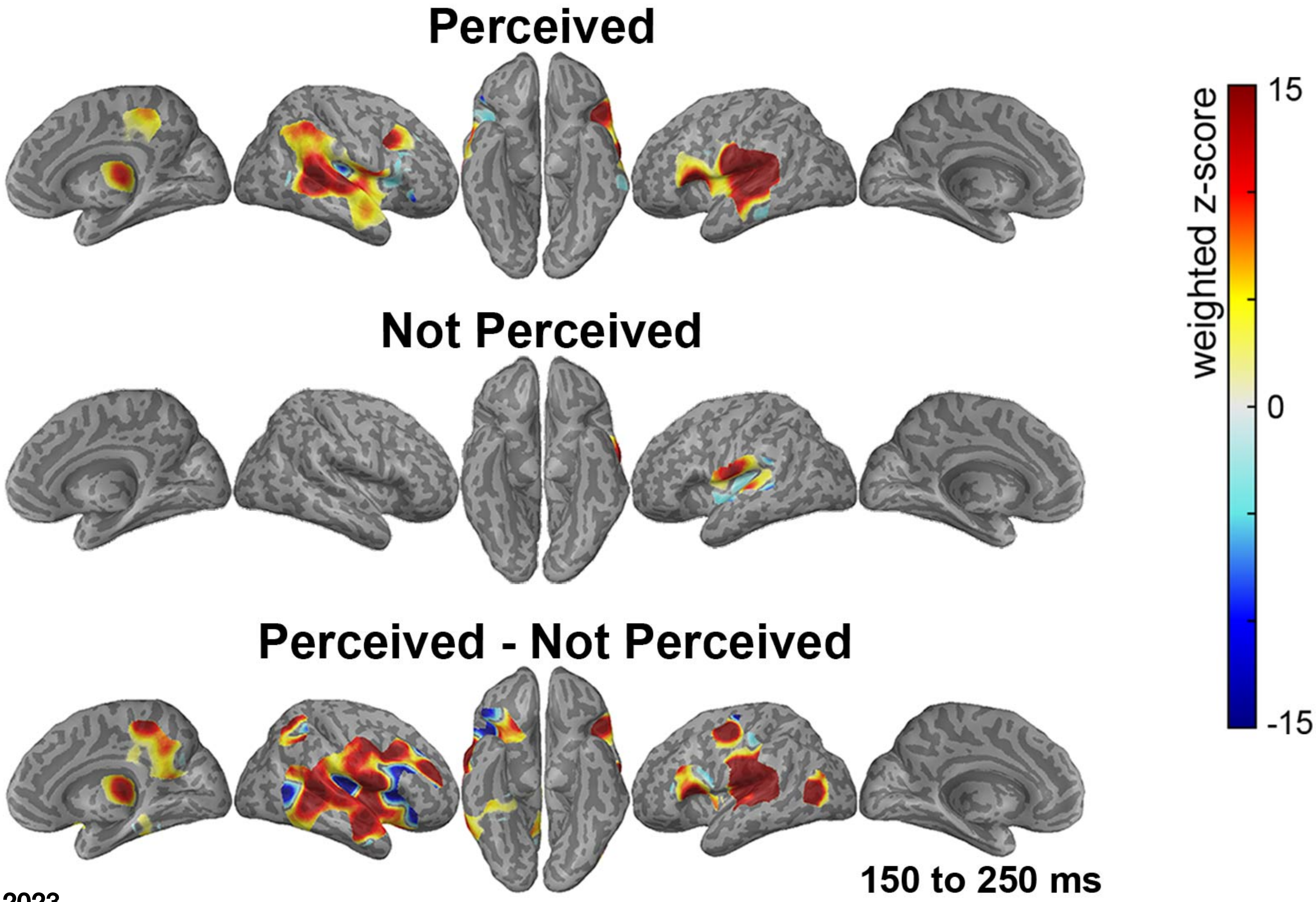


Christison-Lagay et al, 2023

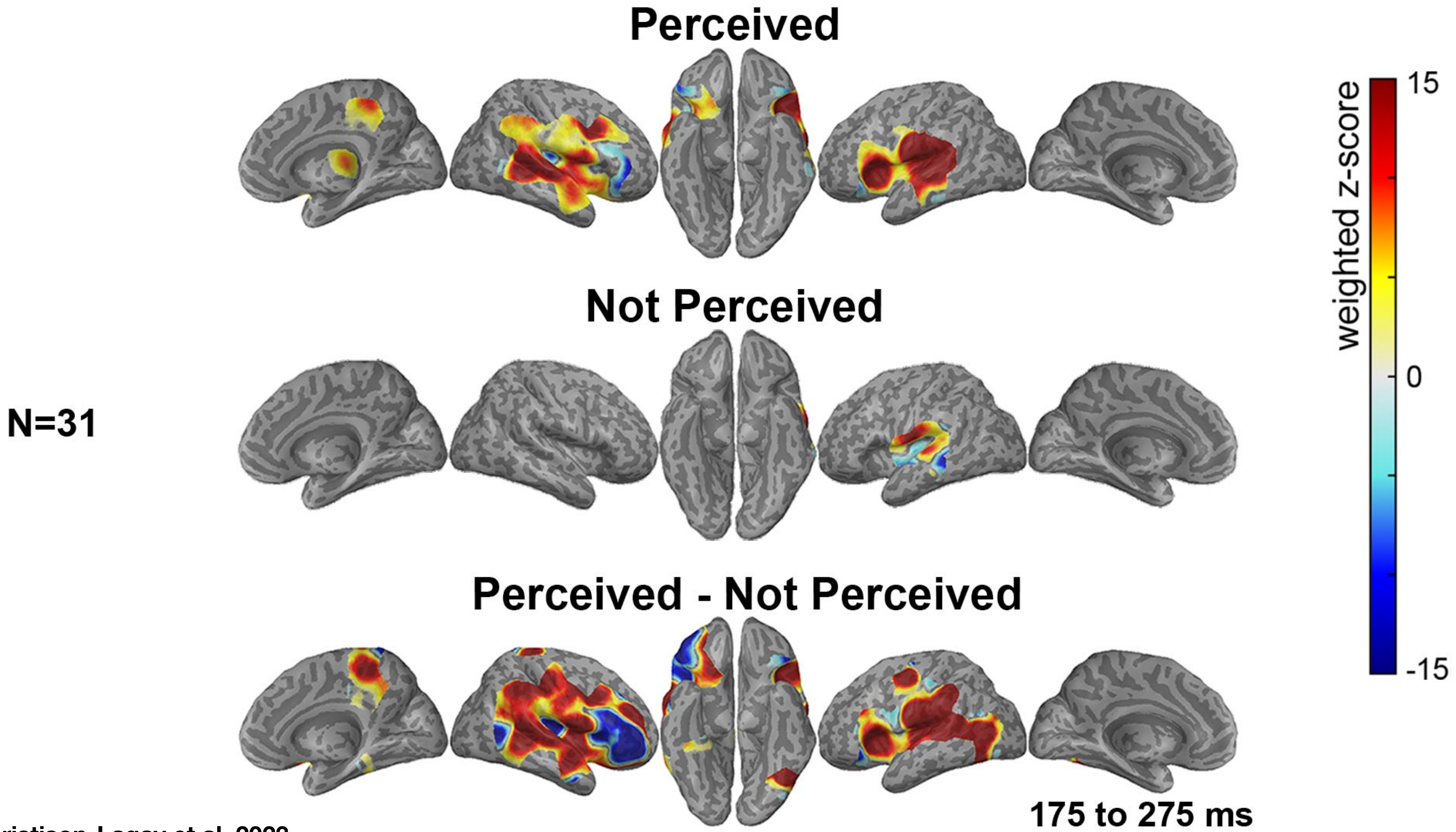


Christison-Lagay et al, 2023

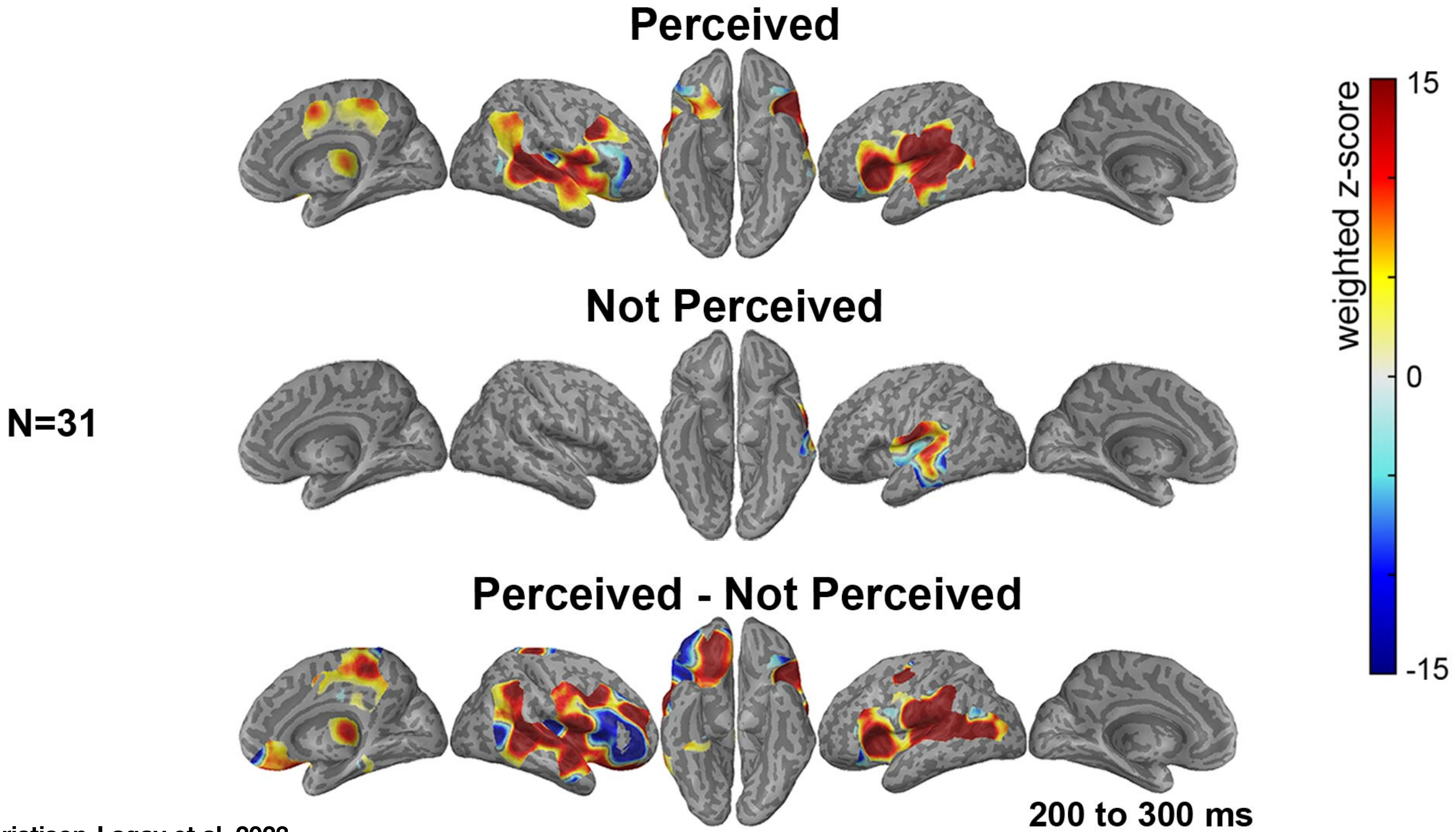


Christison-Lagay et al, 2023

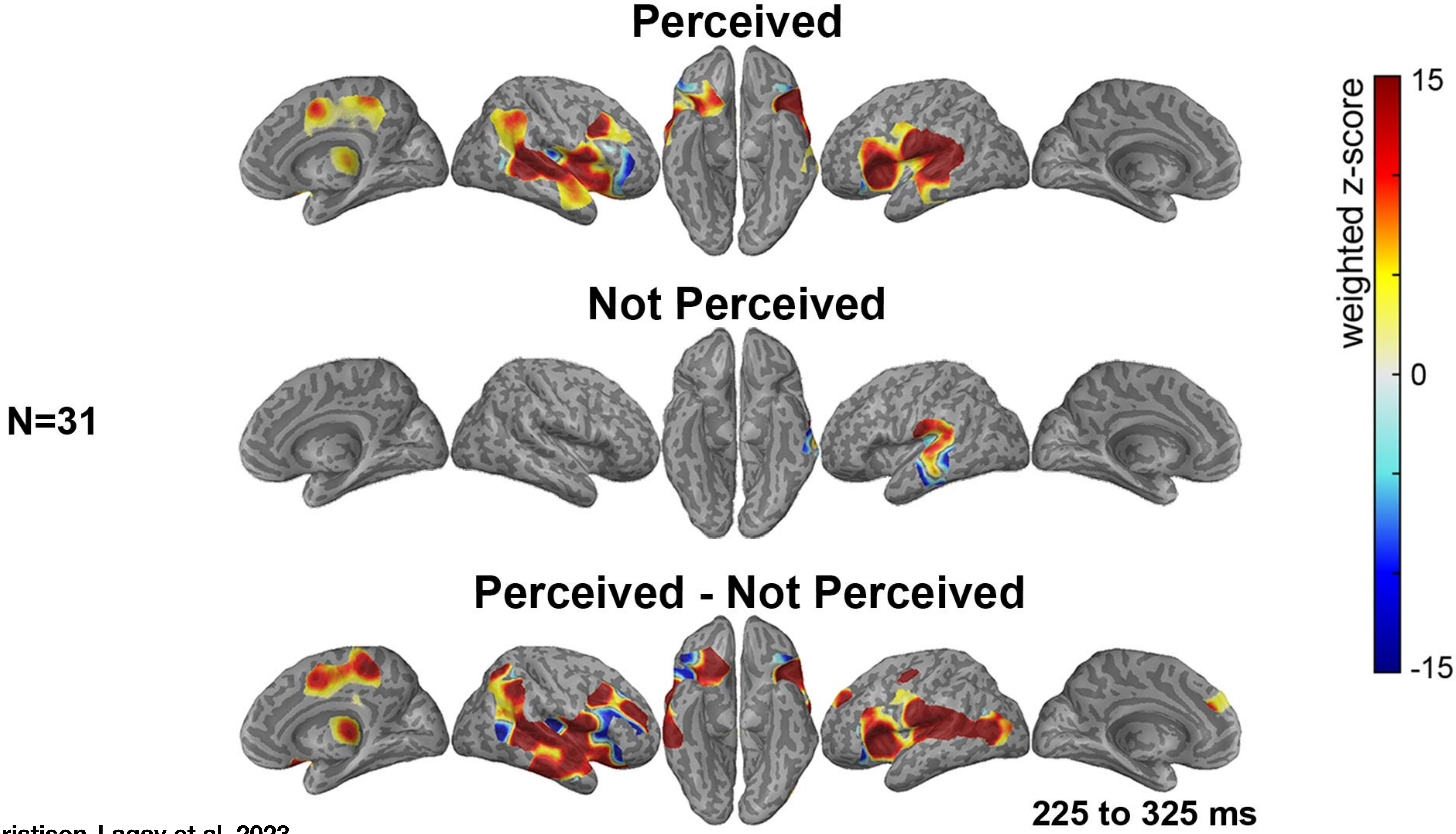


Christison-Lagay et al, 2023

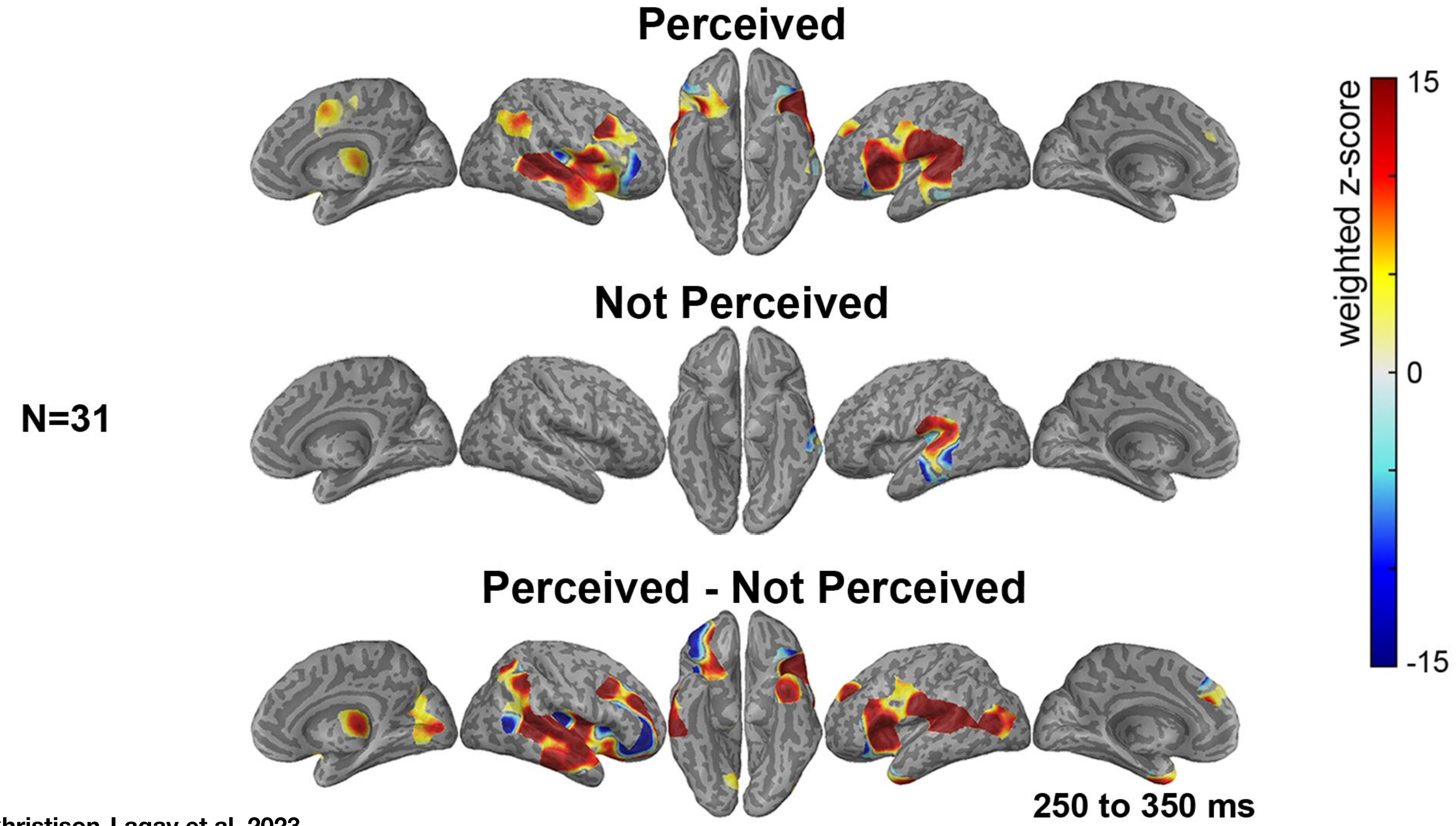


Christison-Lagay et al, 2023

N=31

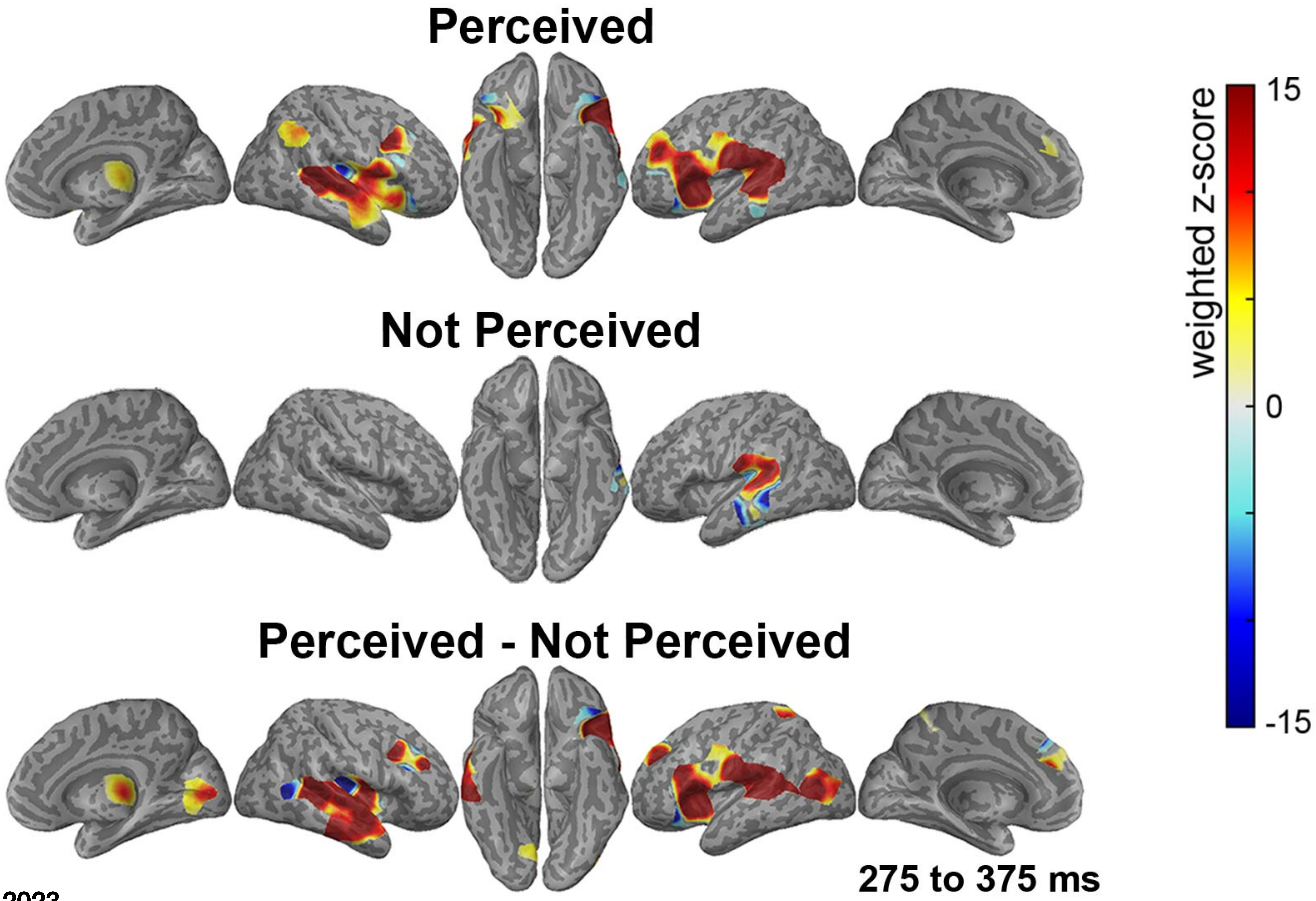


Christison-Lagay et al, 2023

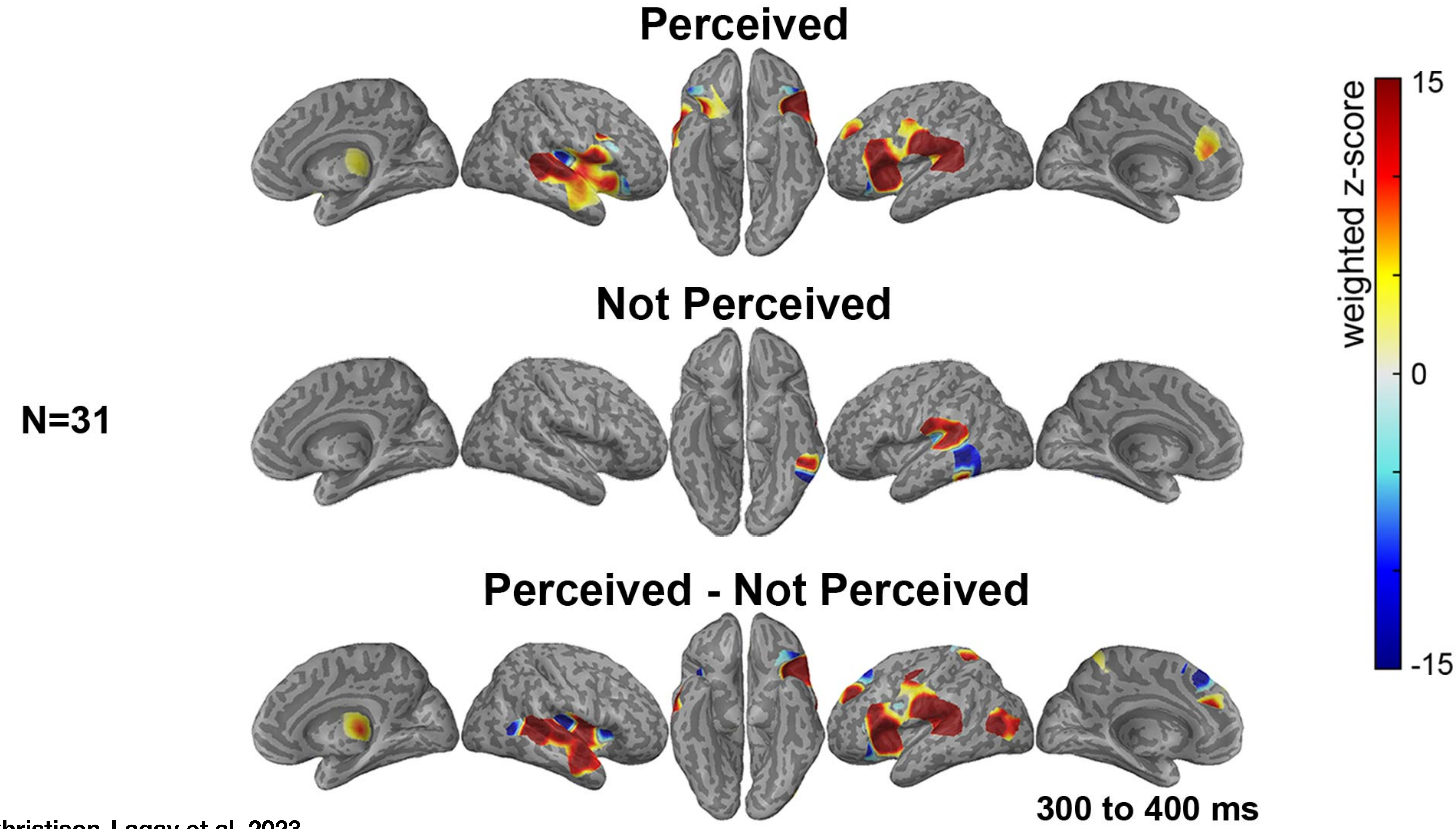

Christison-Lagay et al, 2023

N=31

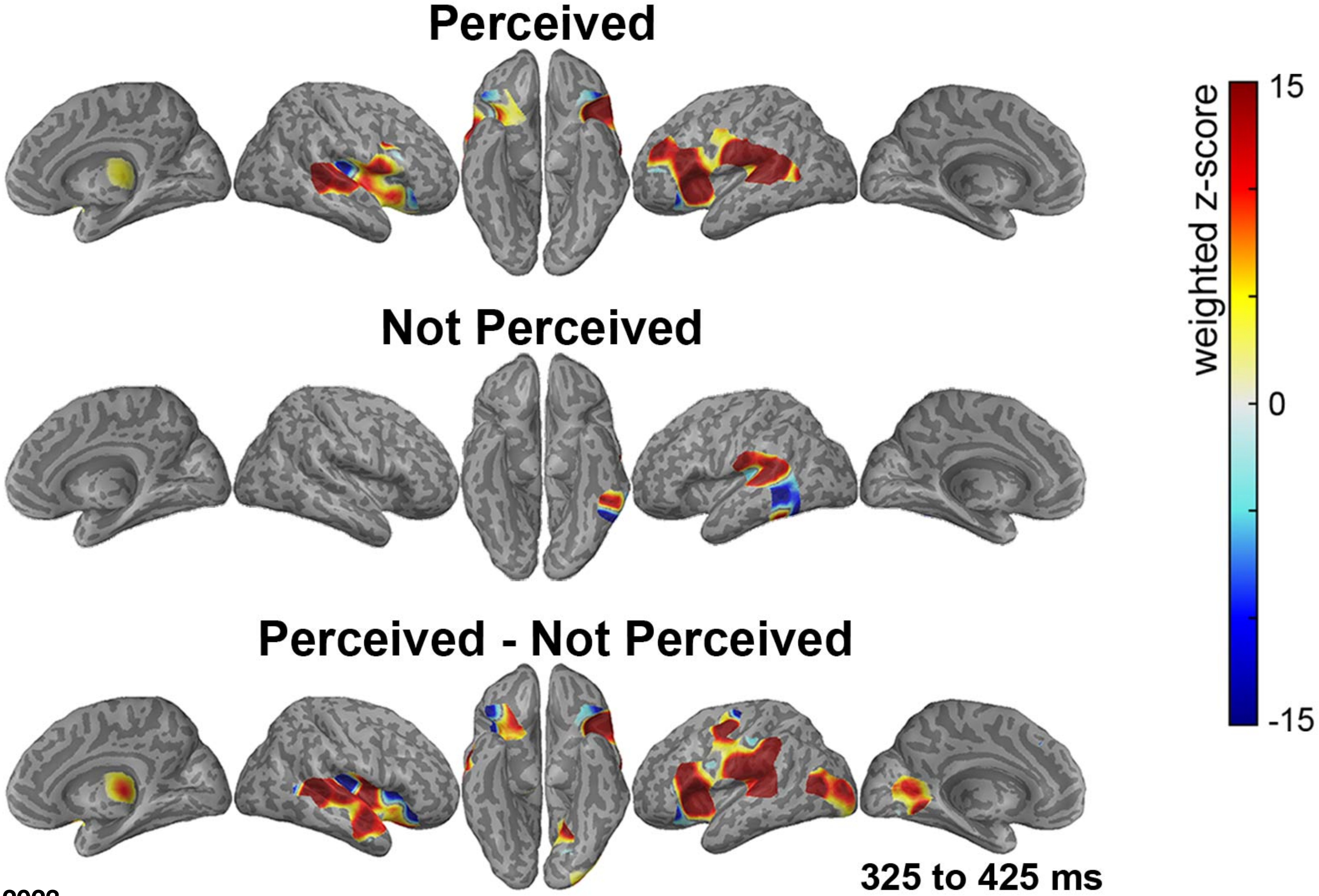


Christison-Lagay et al, 2023

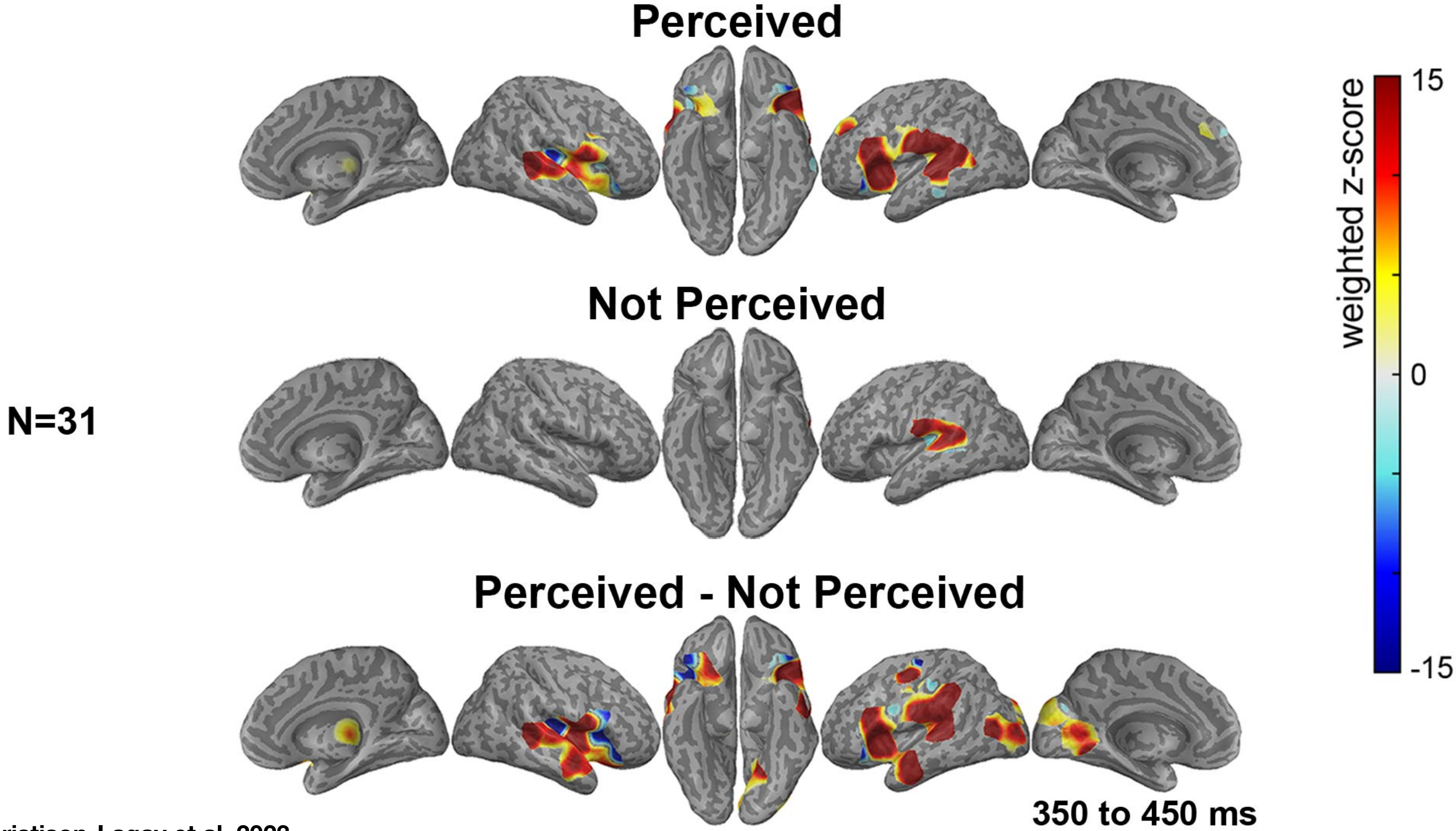


Christison-Lagay et al, 2023

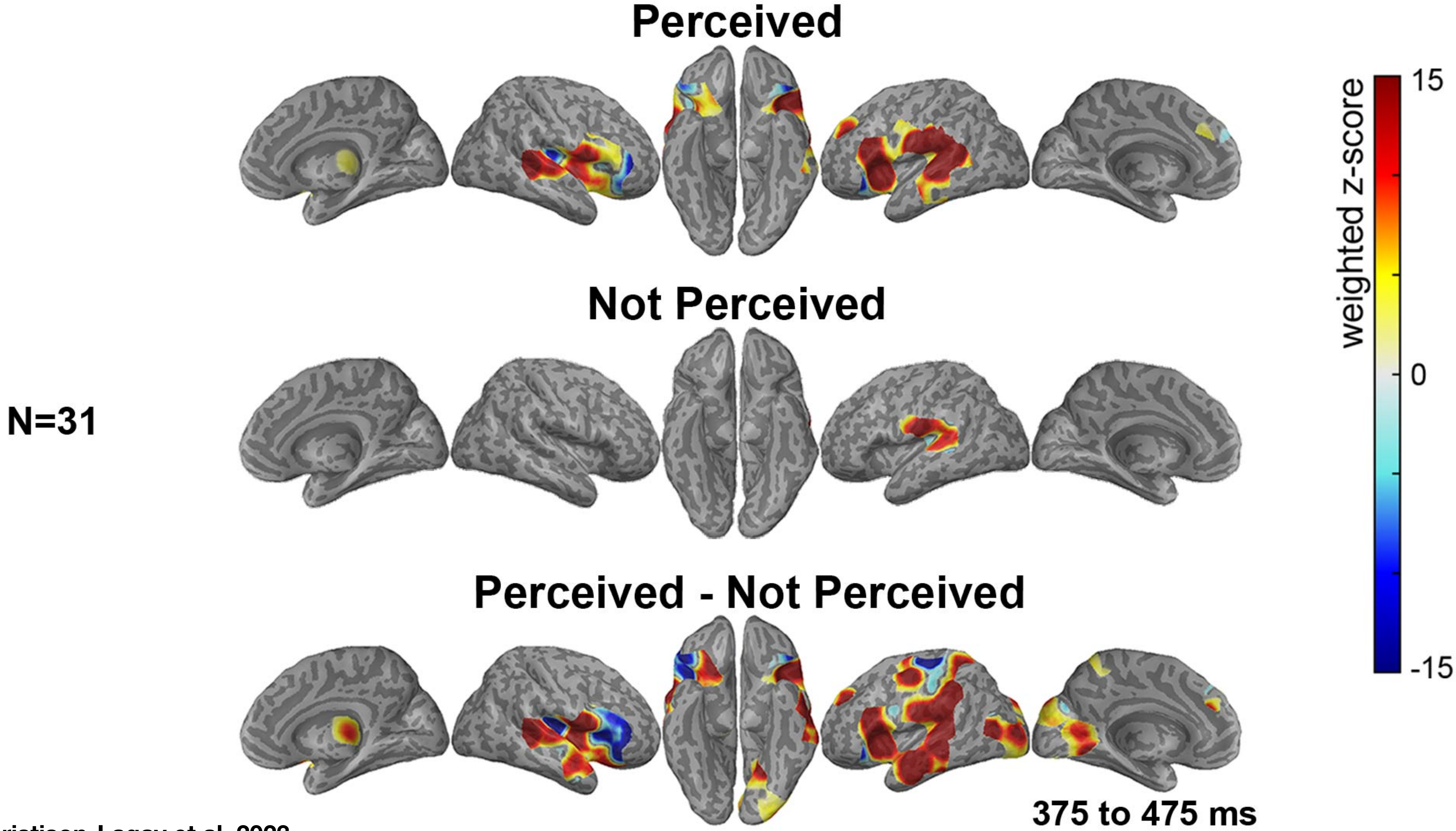


Christison-Lagay et al, 2023

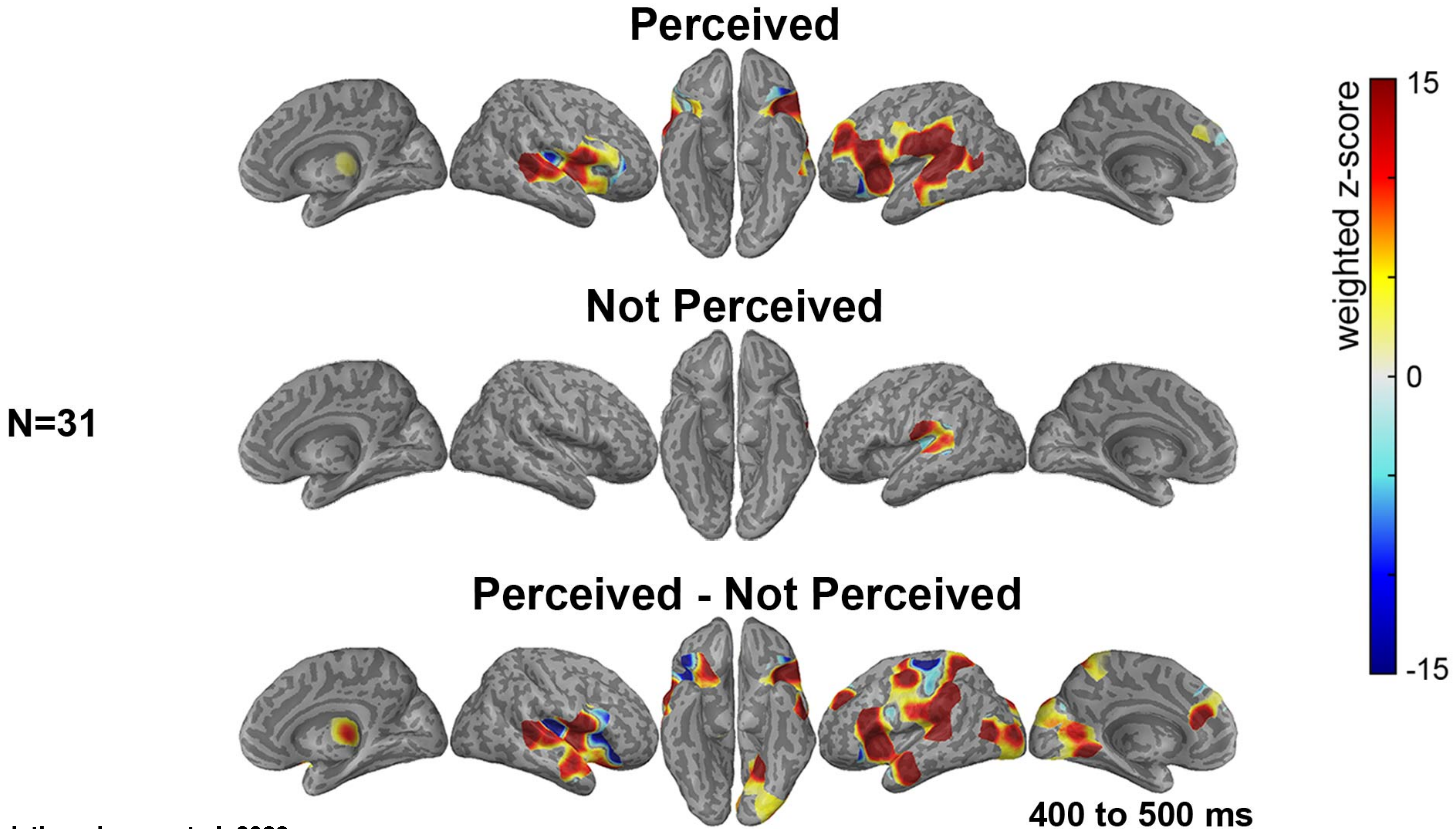


Christison-Lagay et al, 2023

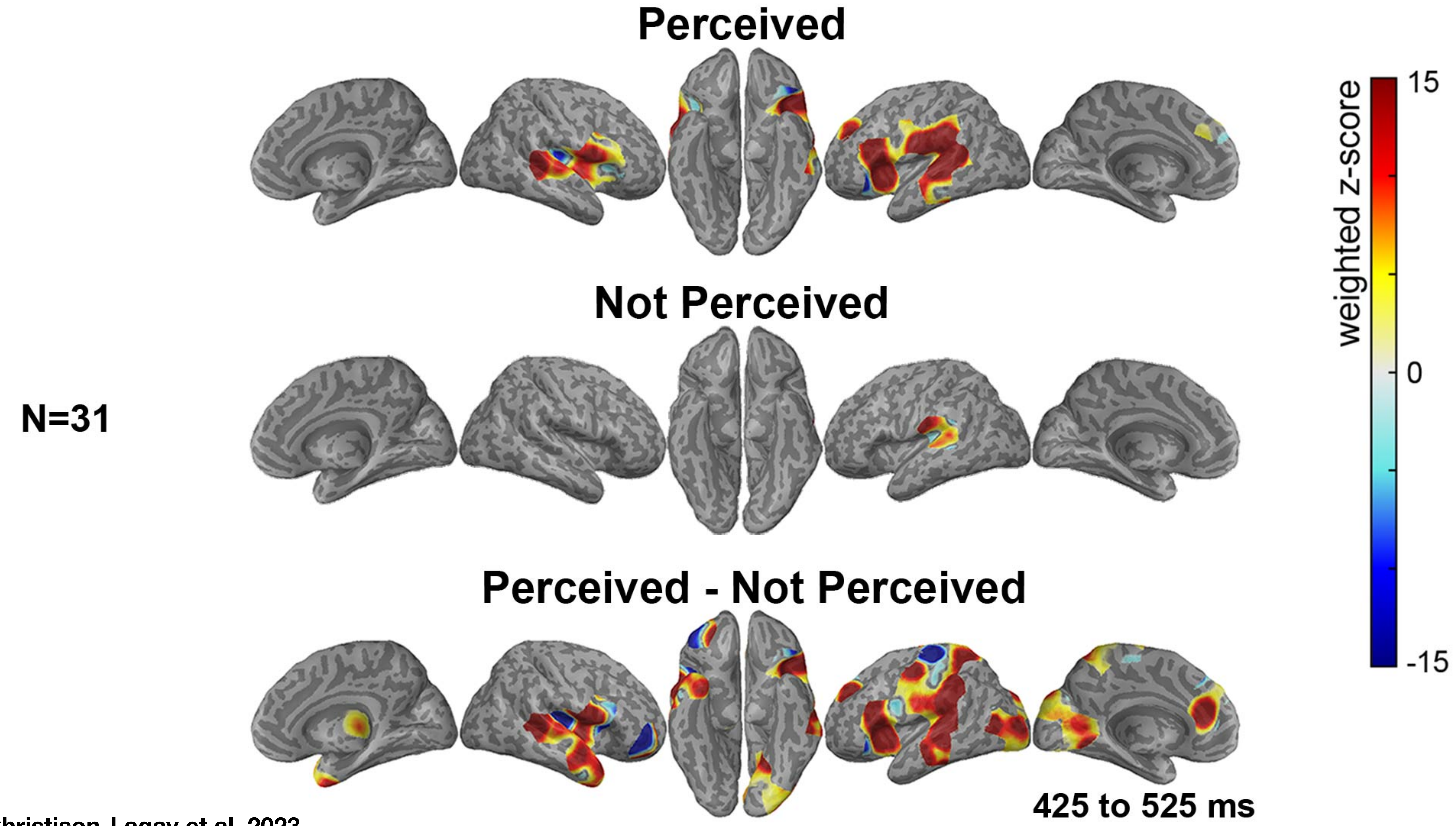


Christison-Lagay et al, 2023

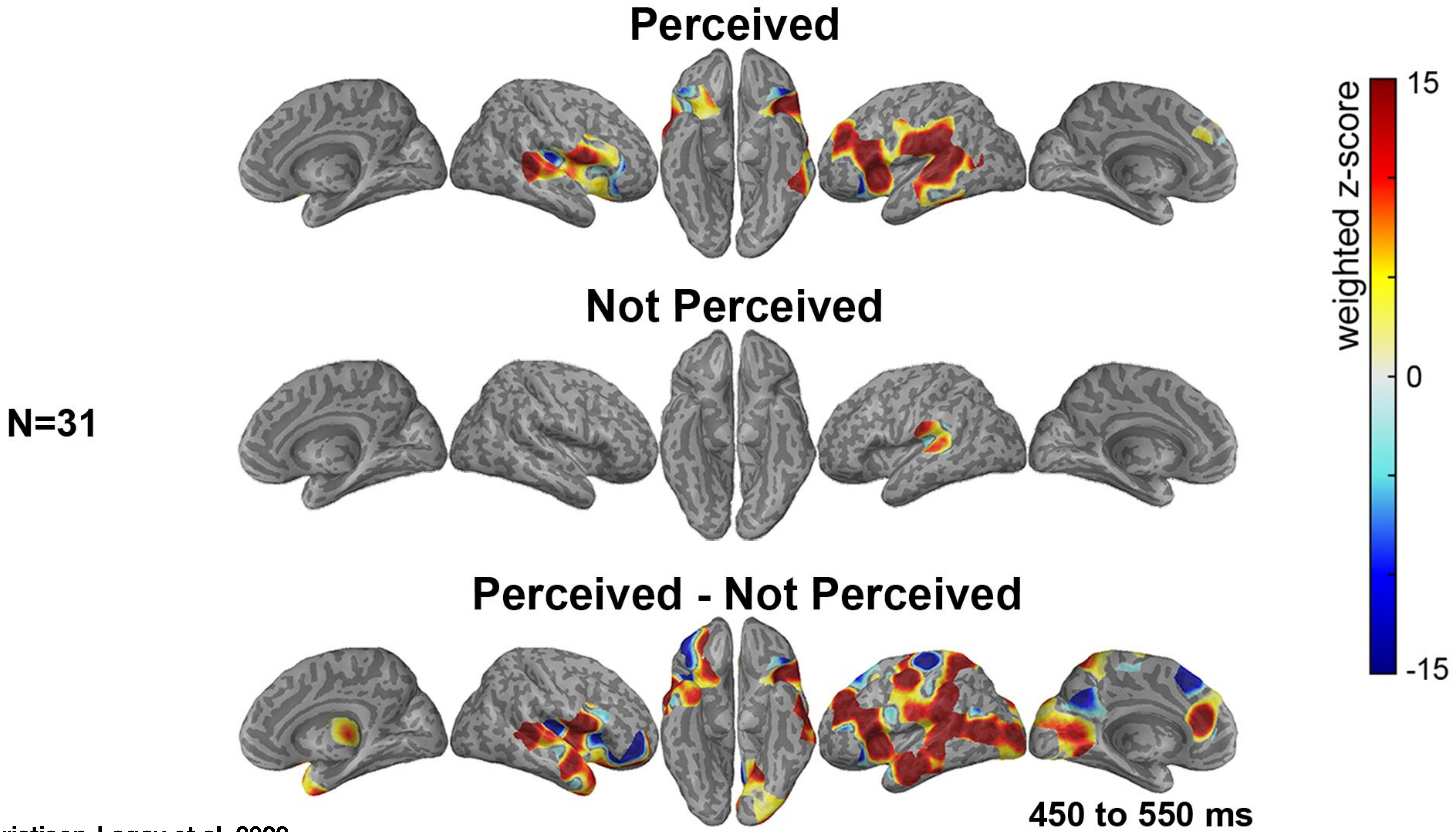


Christison-Lagay et al, 2023

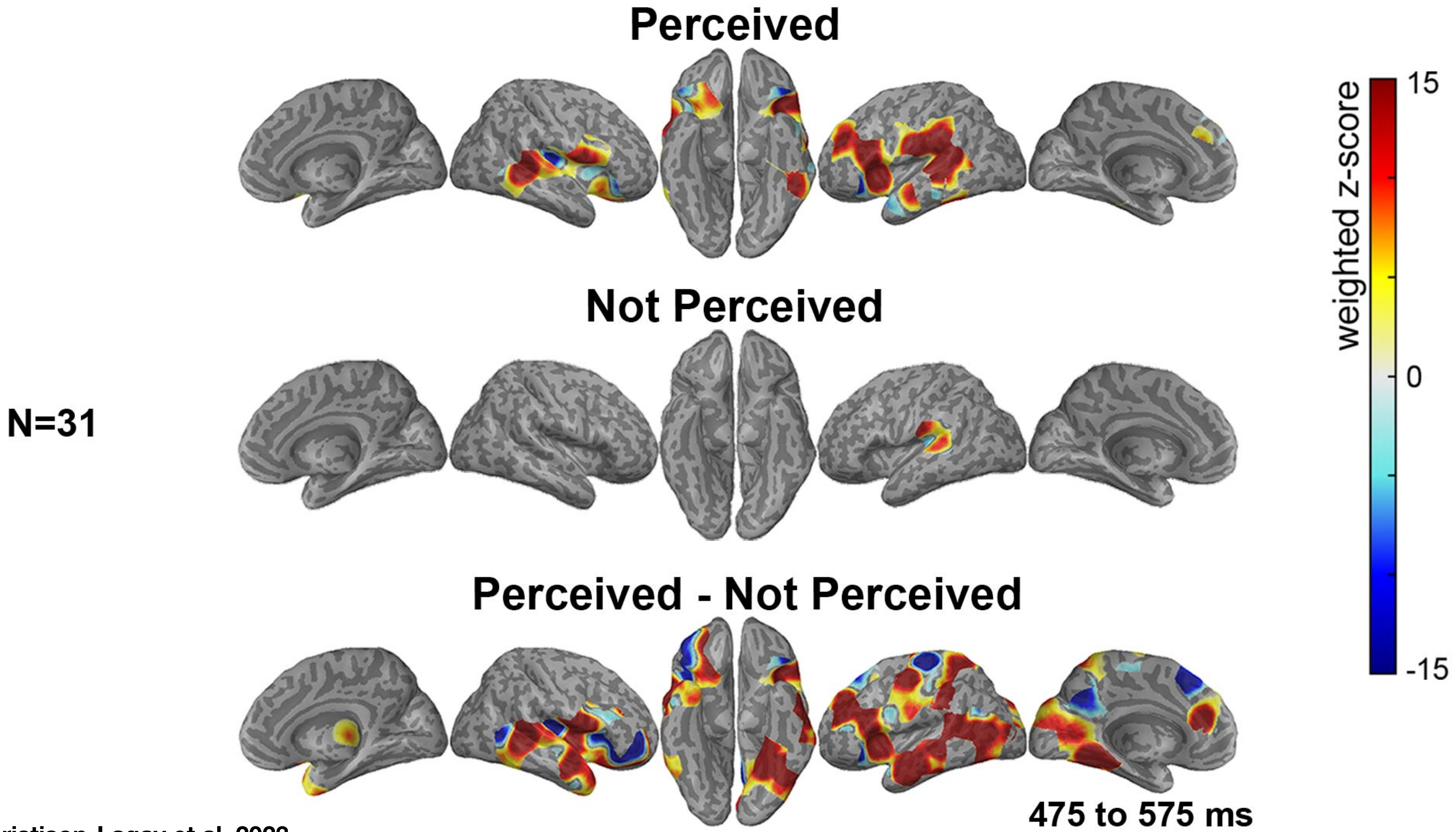


Christison-Lagay et al, 2023

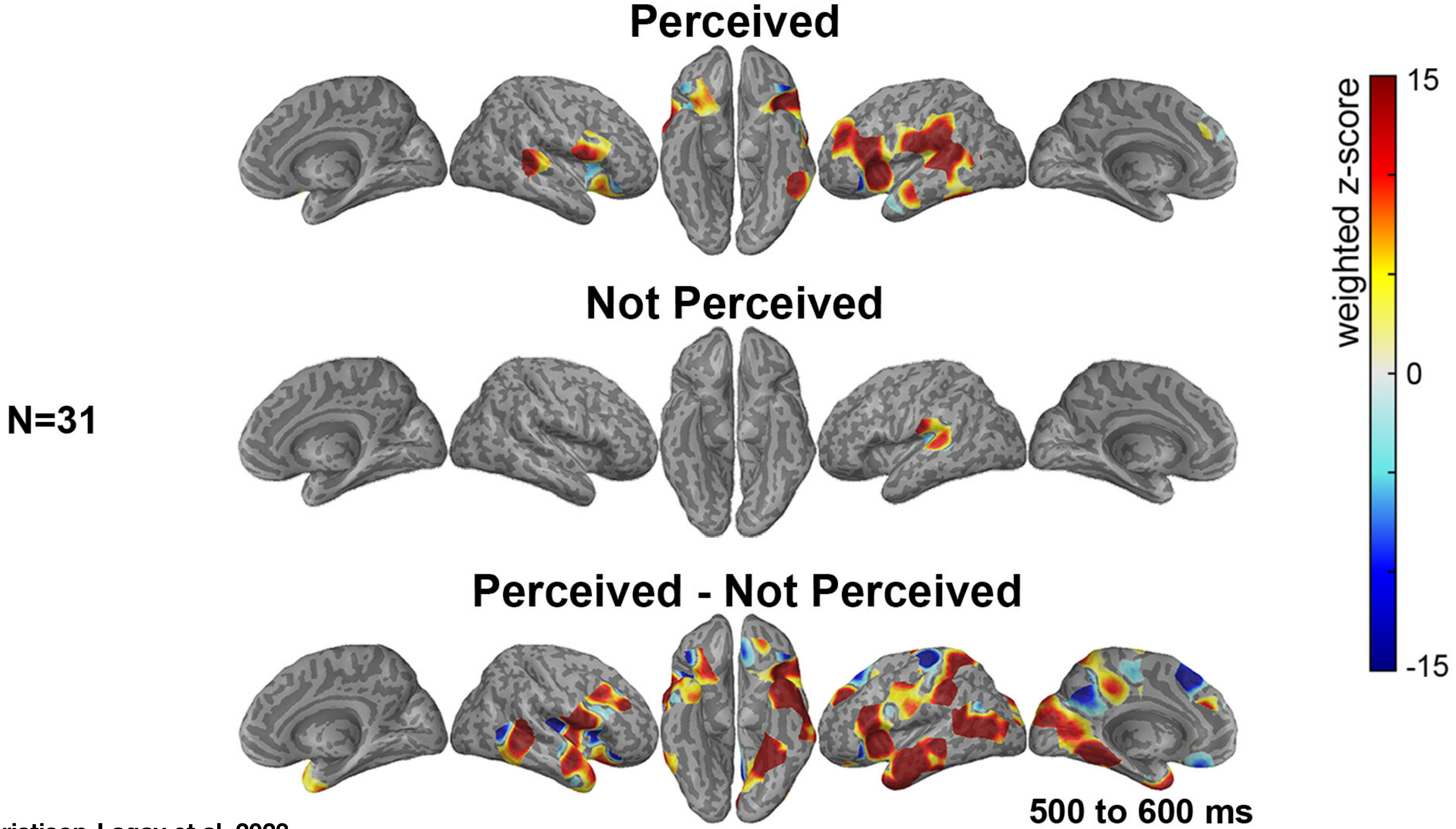


Christison-Lagay et al, 2023

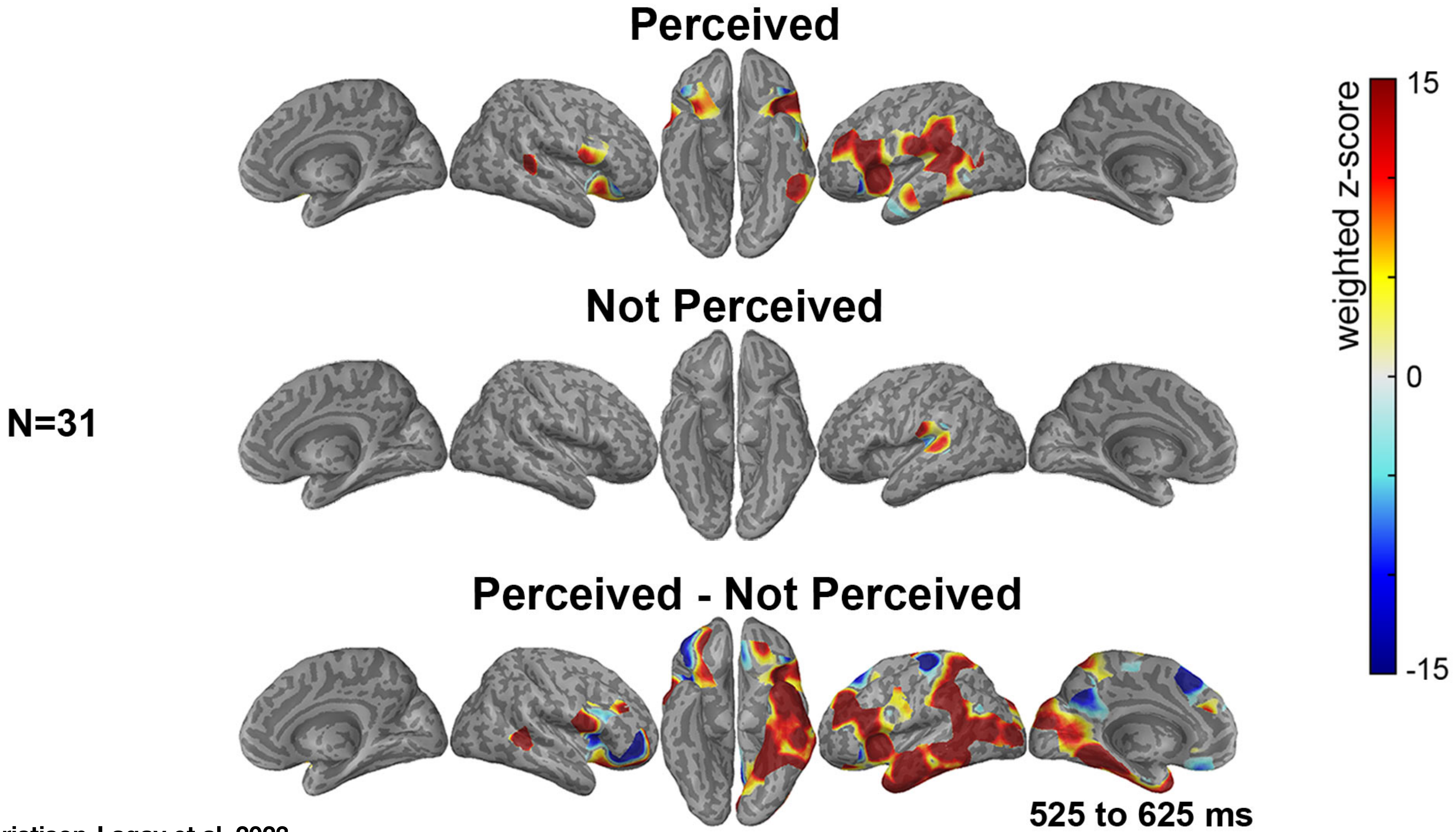


Christison-Lagay et al, 2023

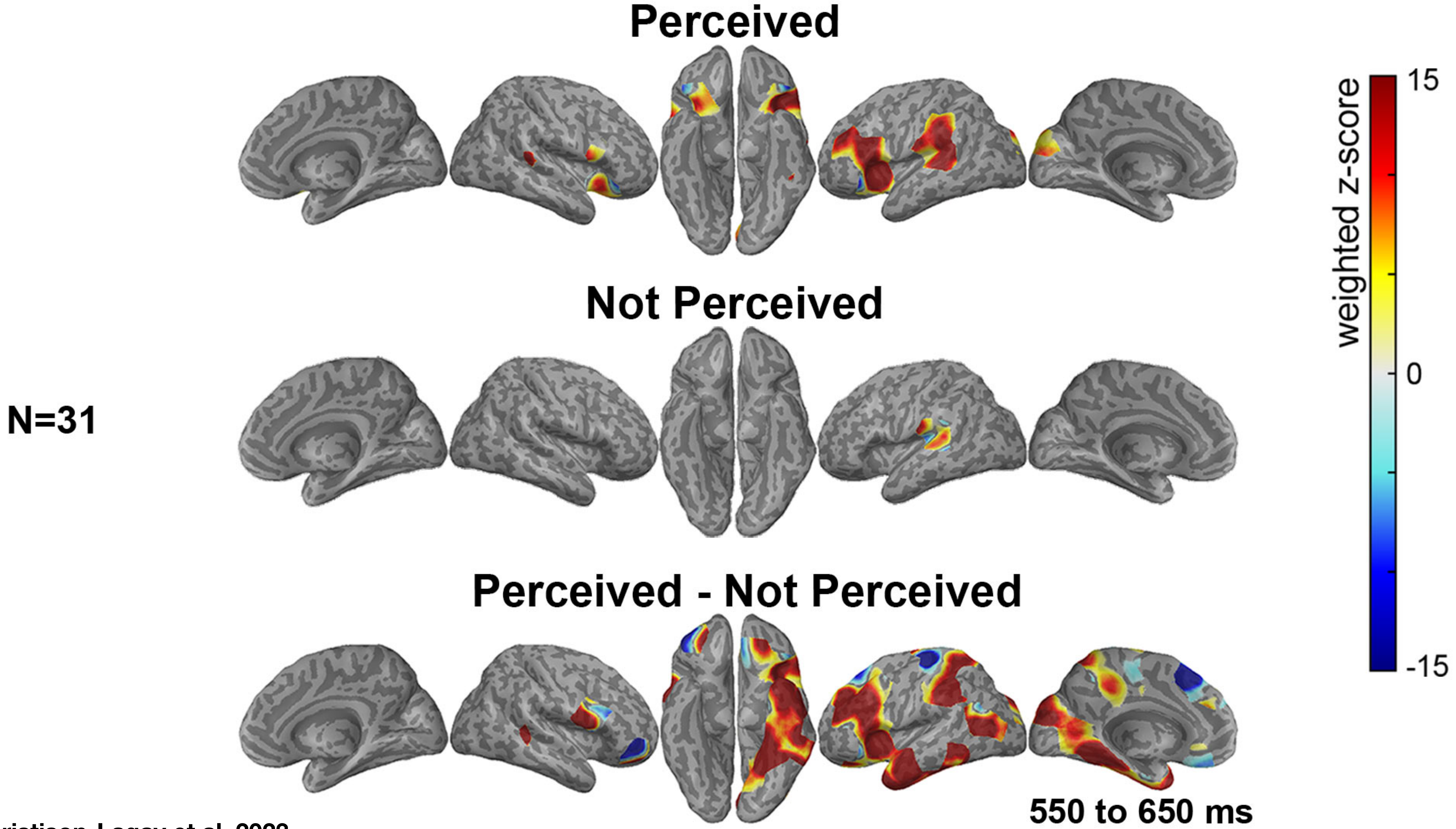


Christison-Lagay et al, 2023

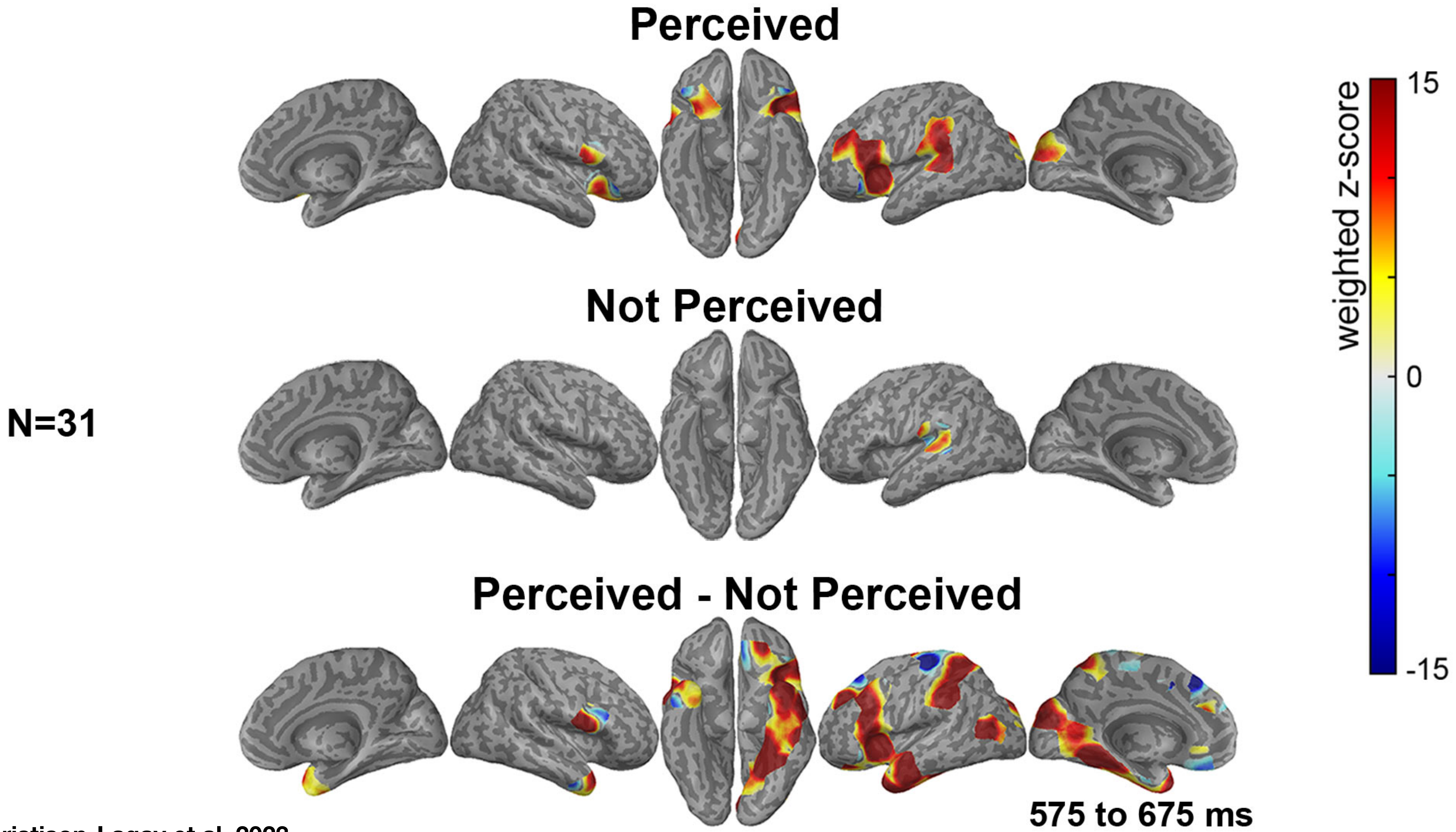

Christison-Lagay et al, 2023

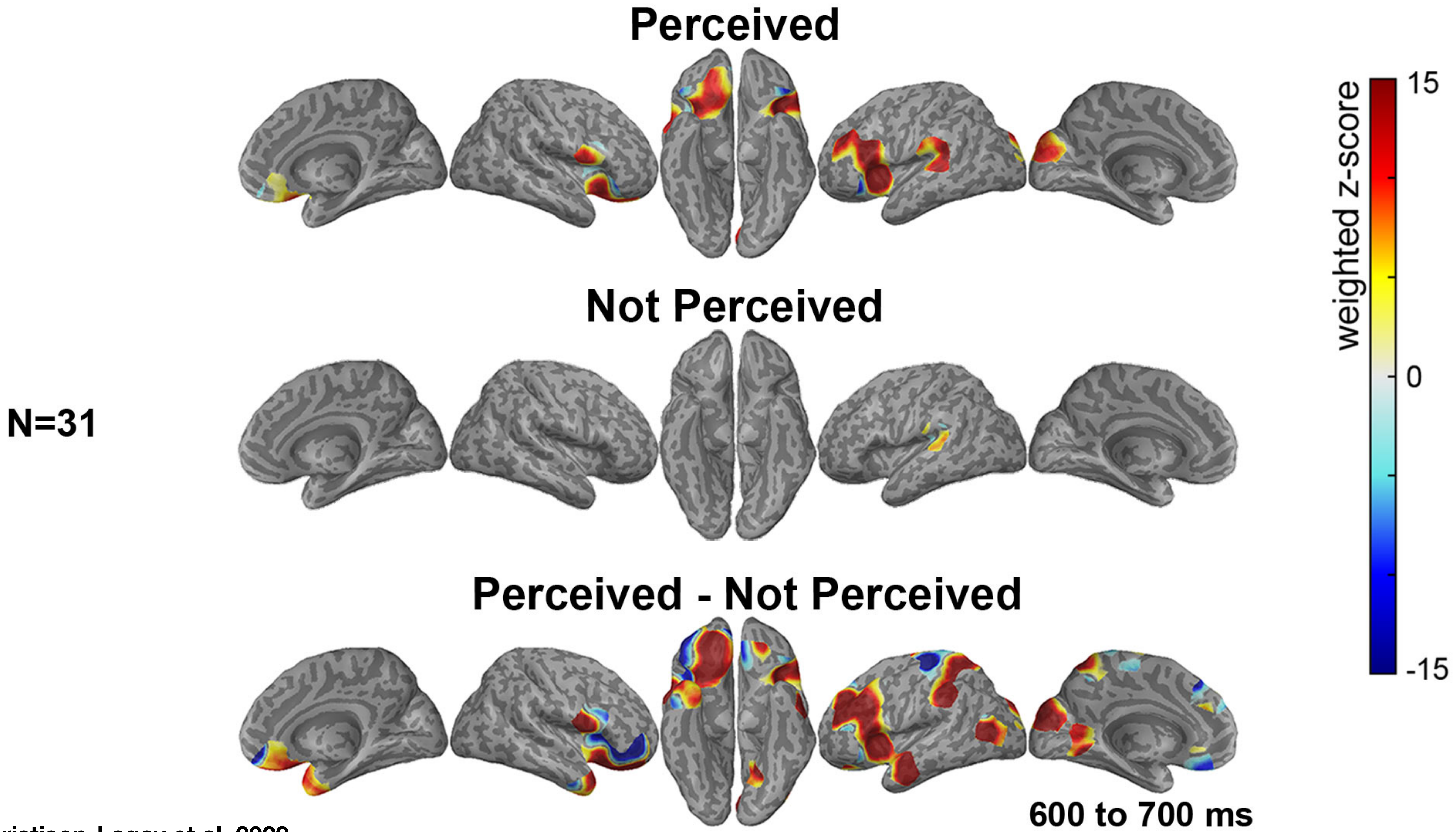

Christison-Lagay et al, 2023

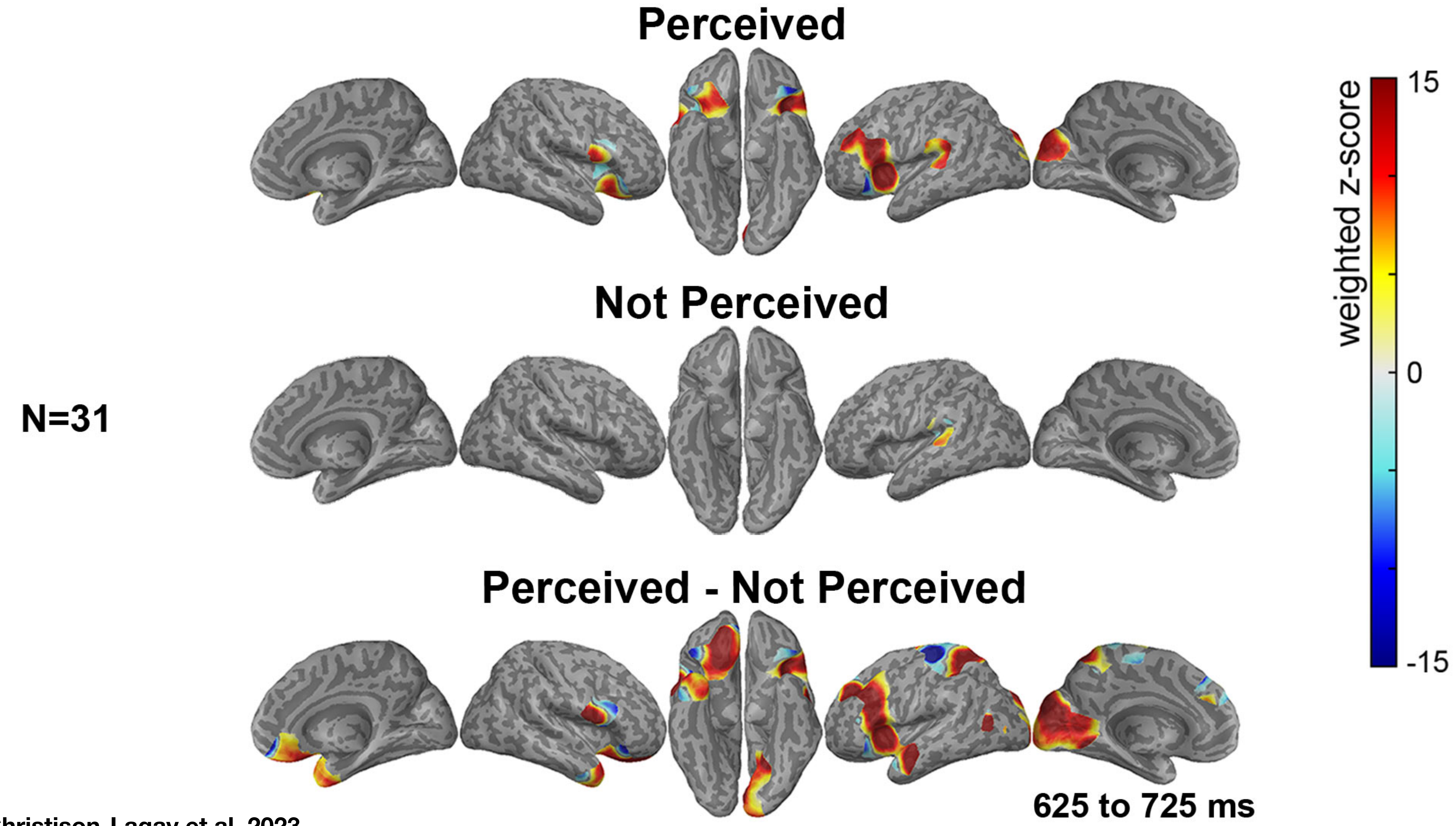


Christison-Lagay et al, 2023

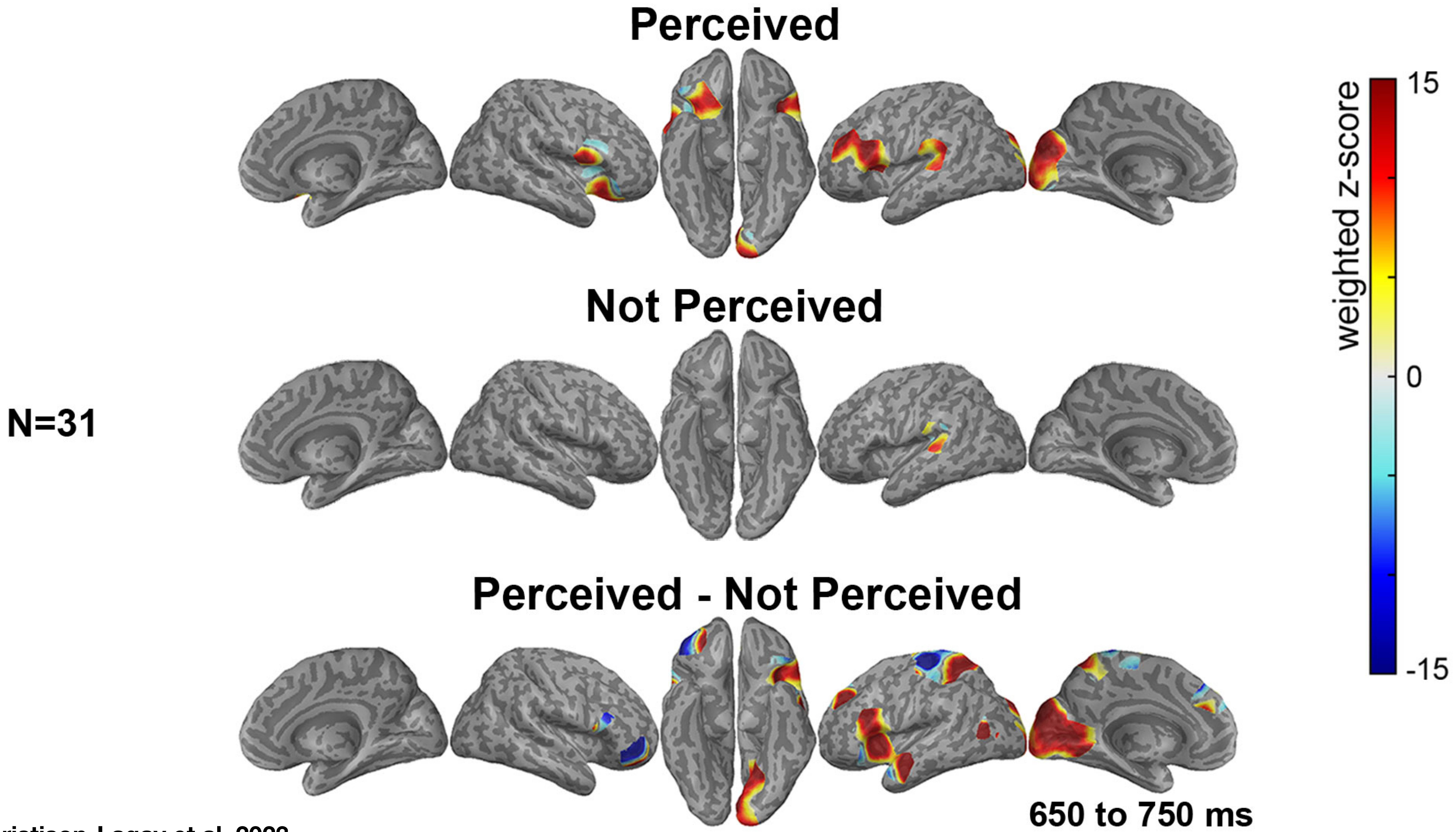


Christison-Lagay et al, 2023

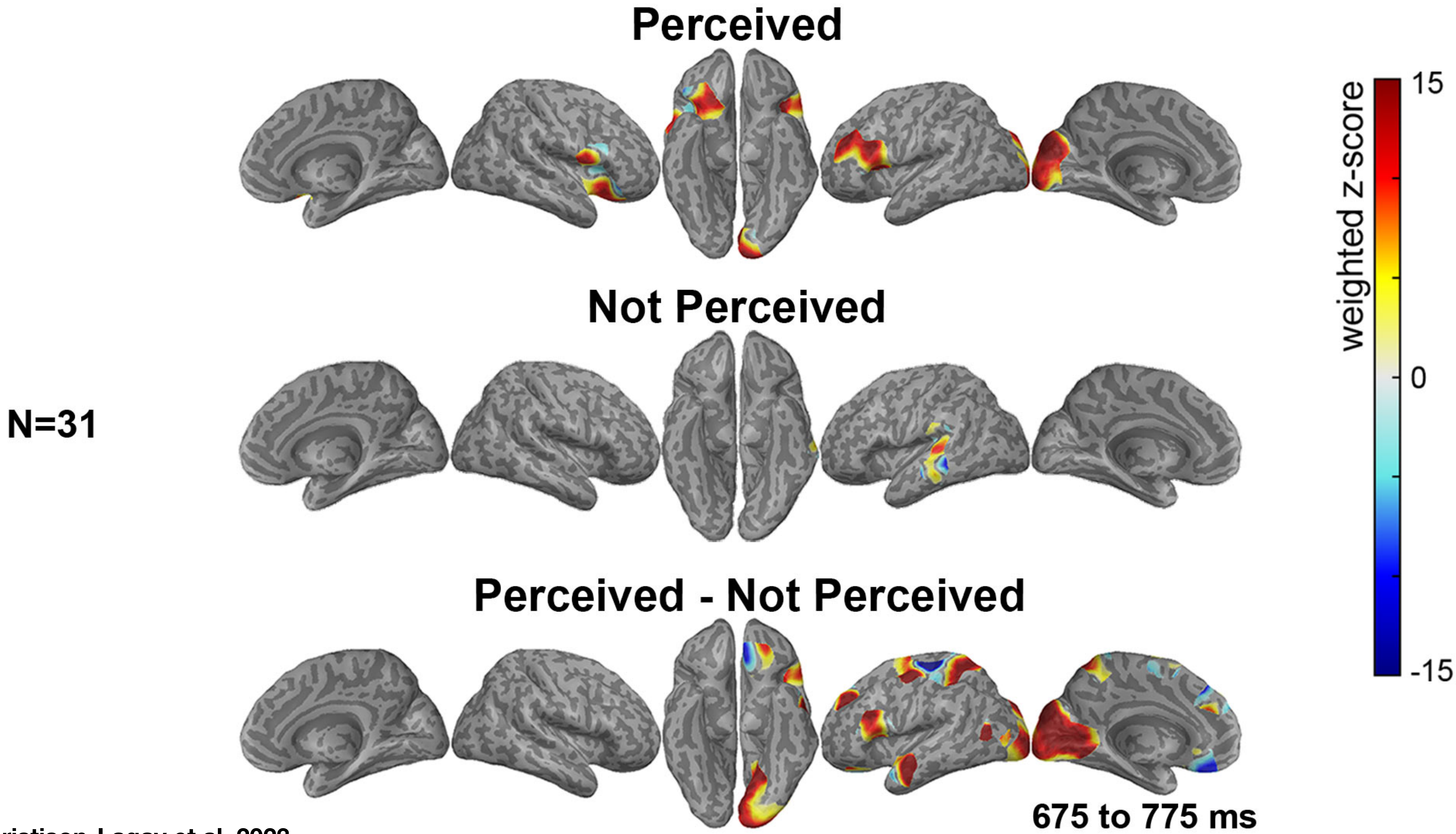


Christison-Lagay et al, 2023

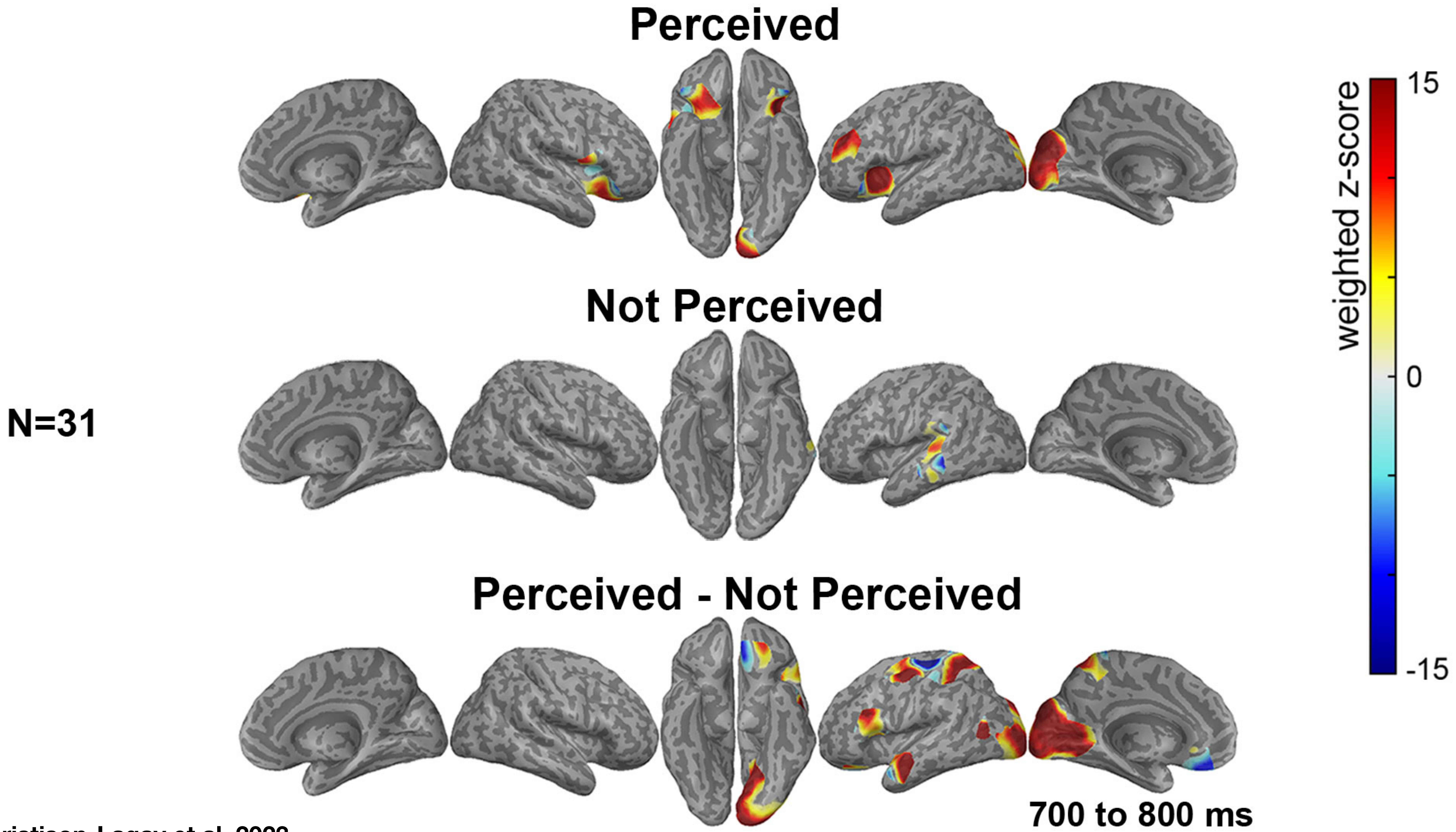


Christison-Lagay et al, 2023

N=31

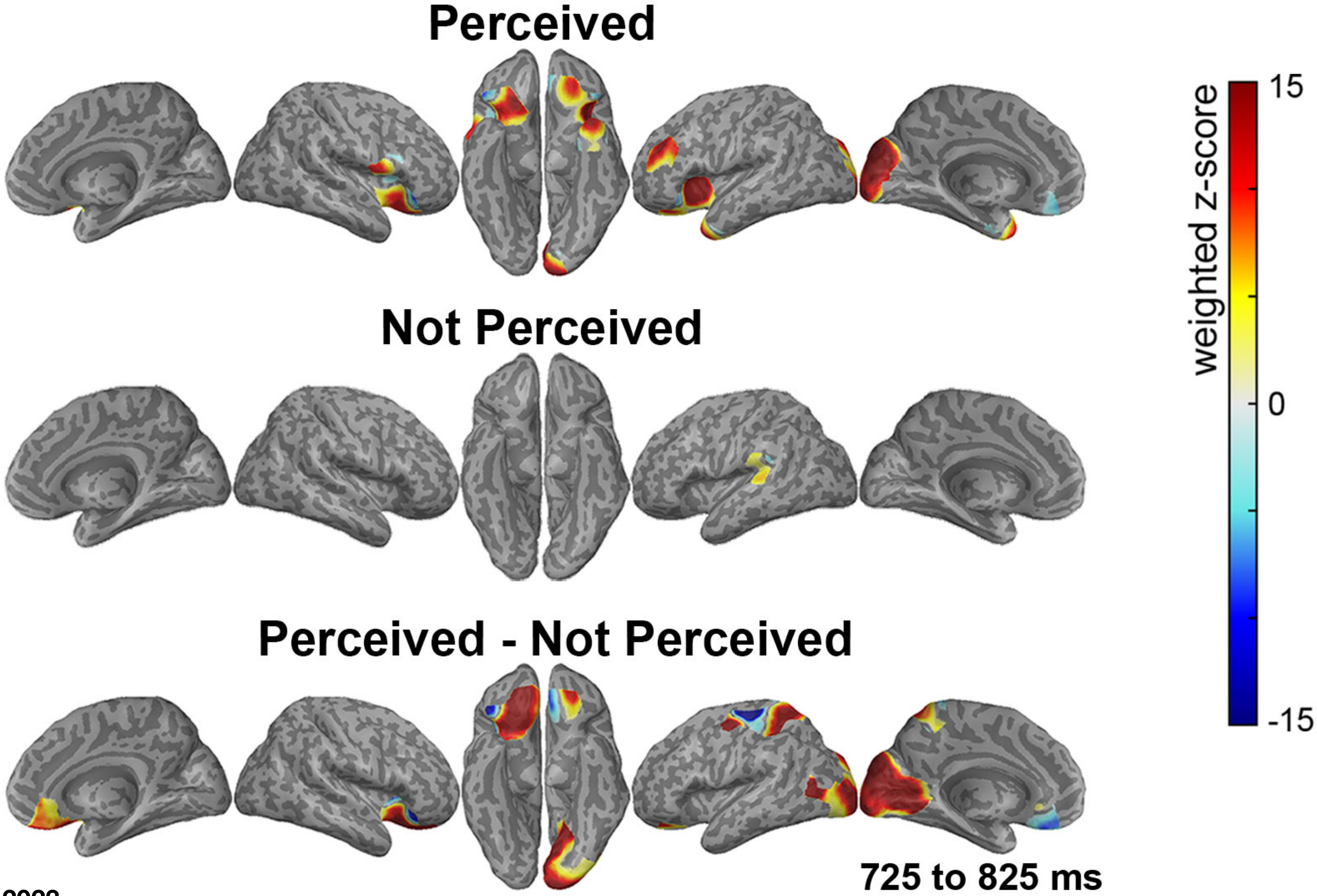


Christison-Lagay et al, 2023

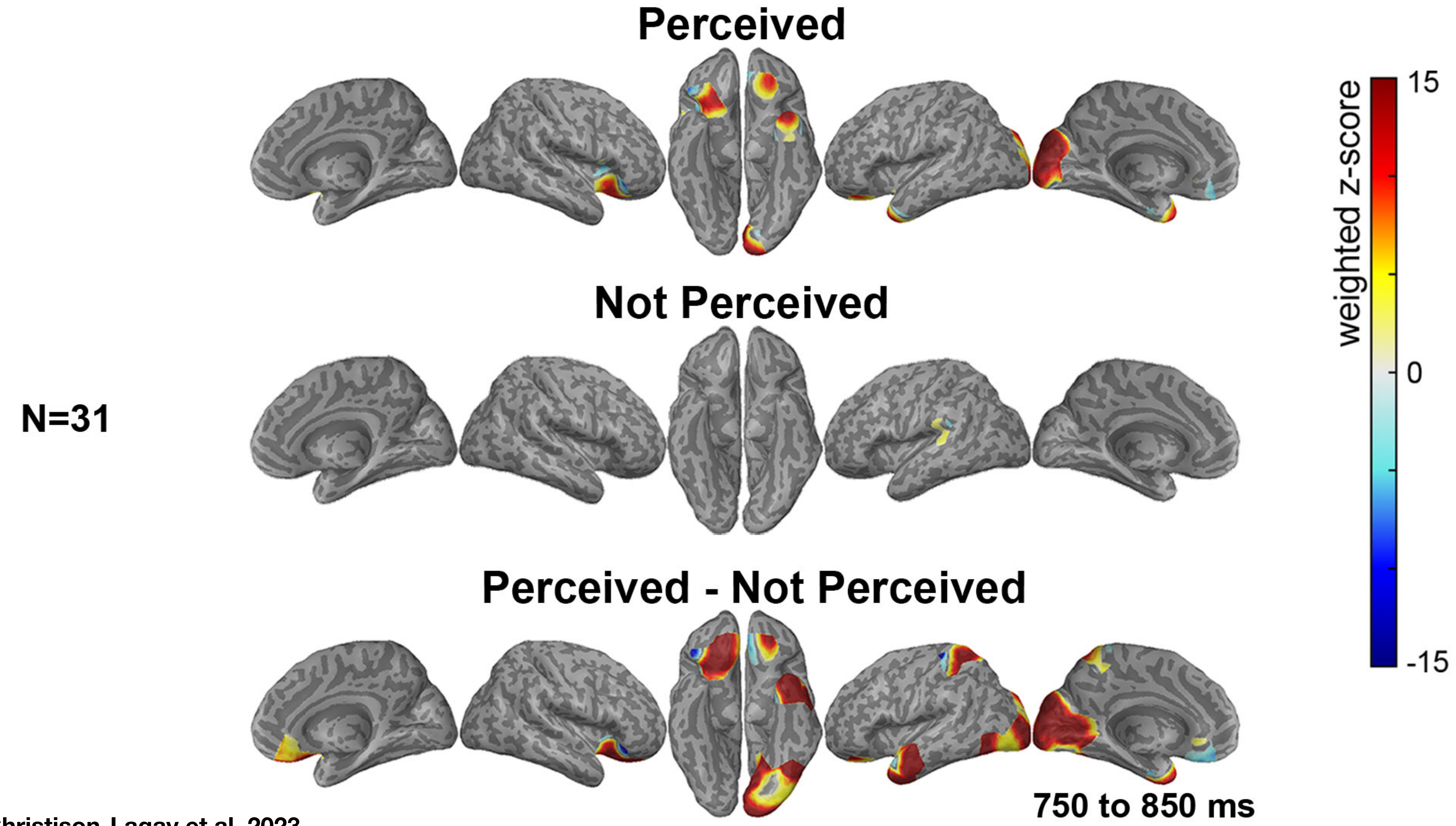


Christison-Lagay et al, 2023

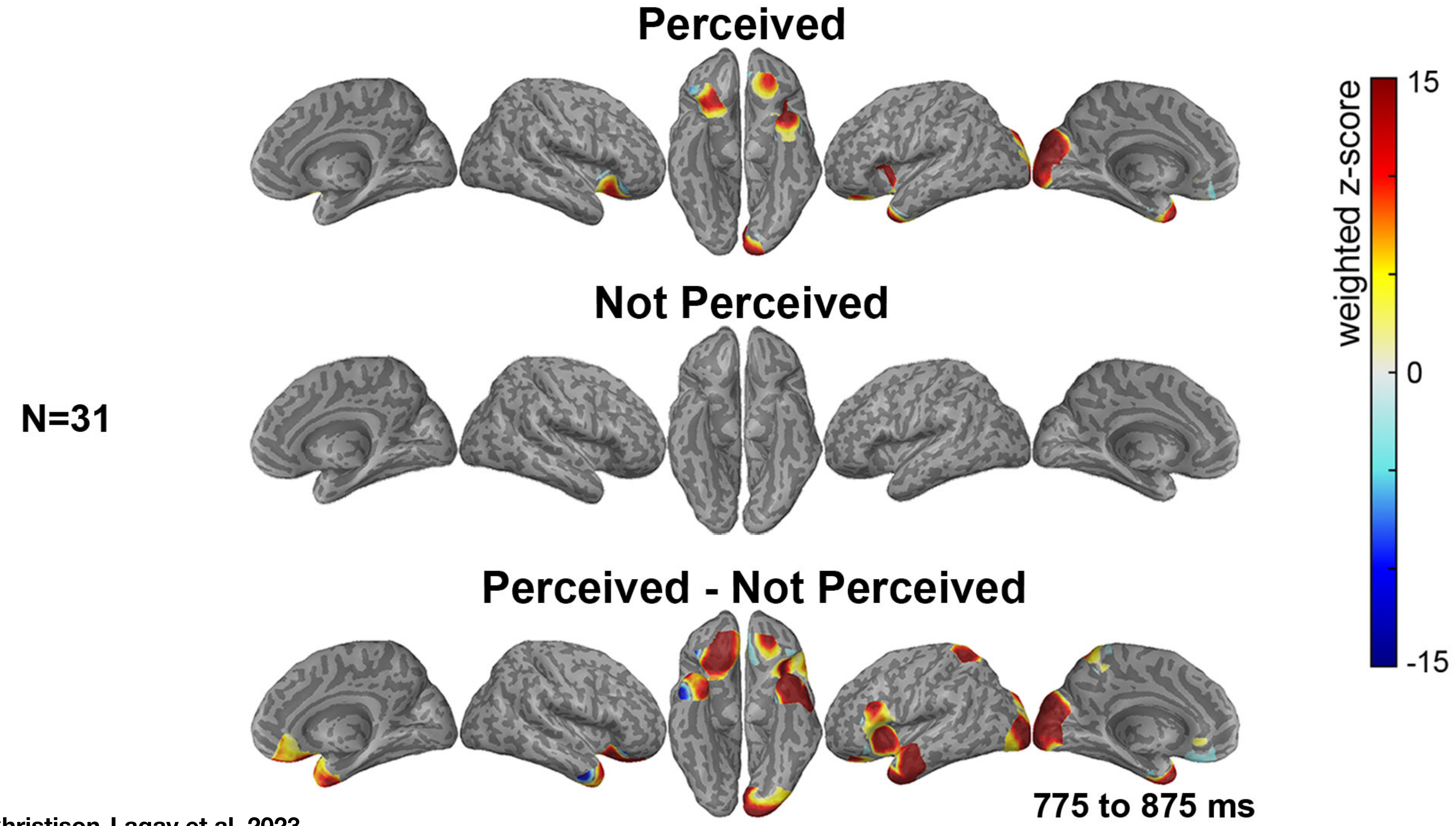


Christison-Lagay et al, 2023

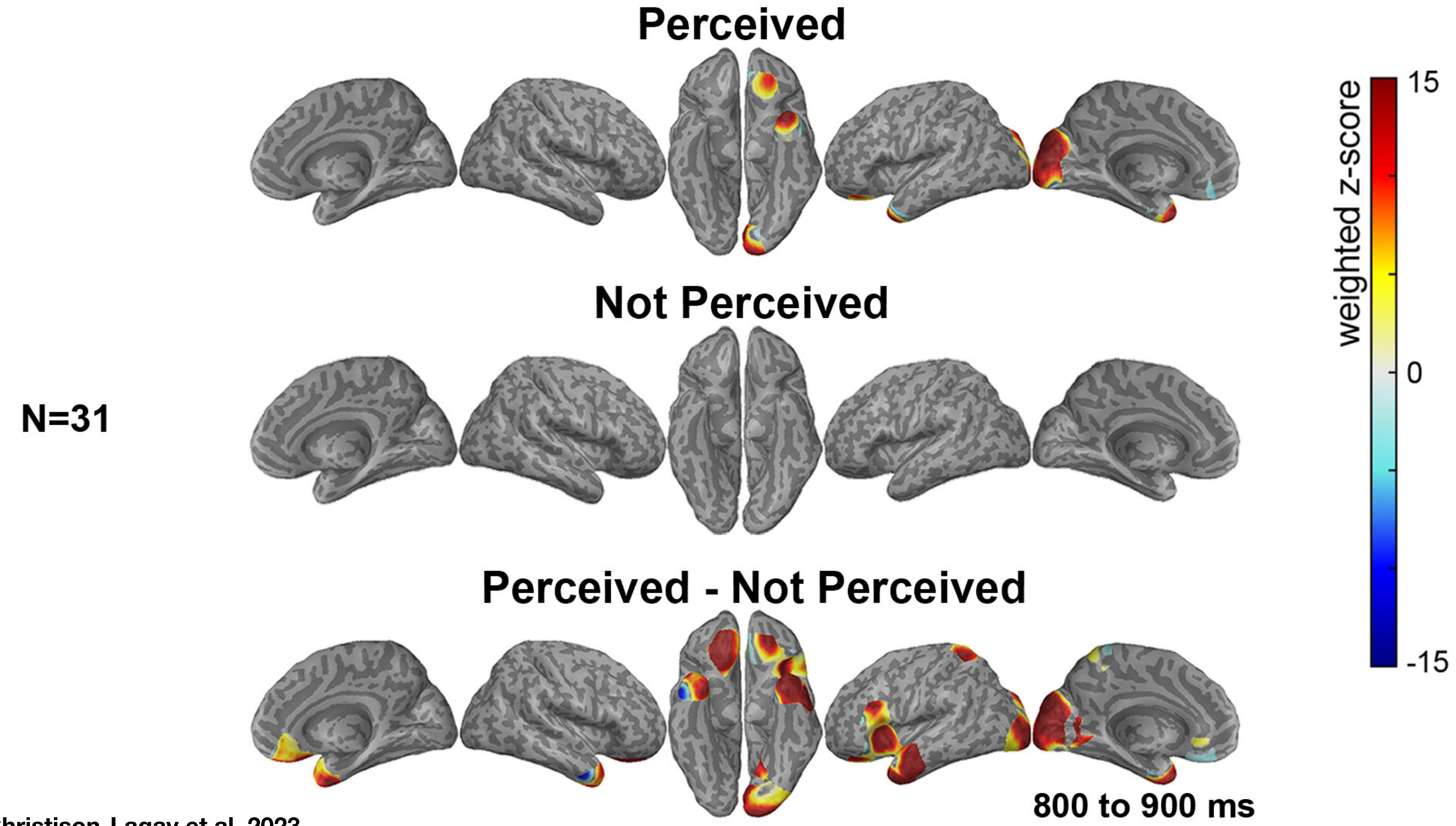


Christison-Lagay et al, 2023

N=31

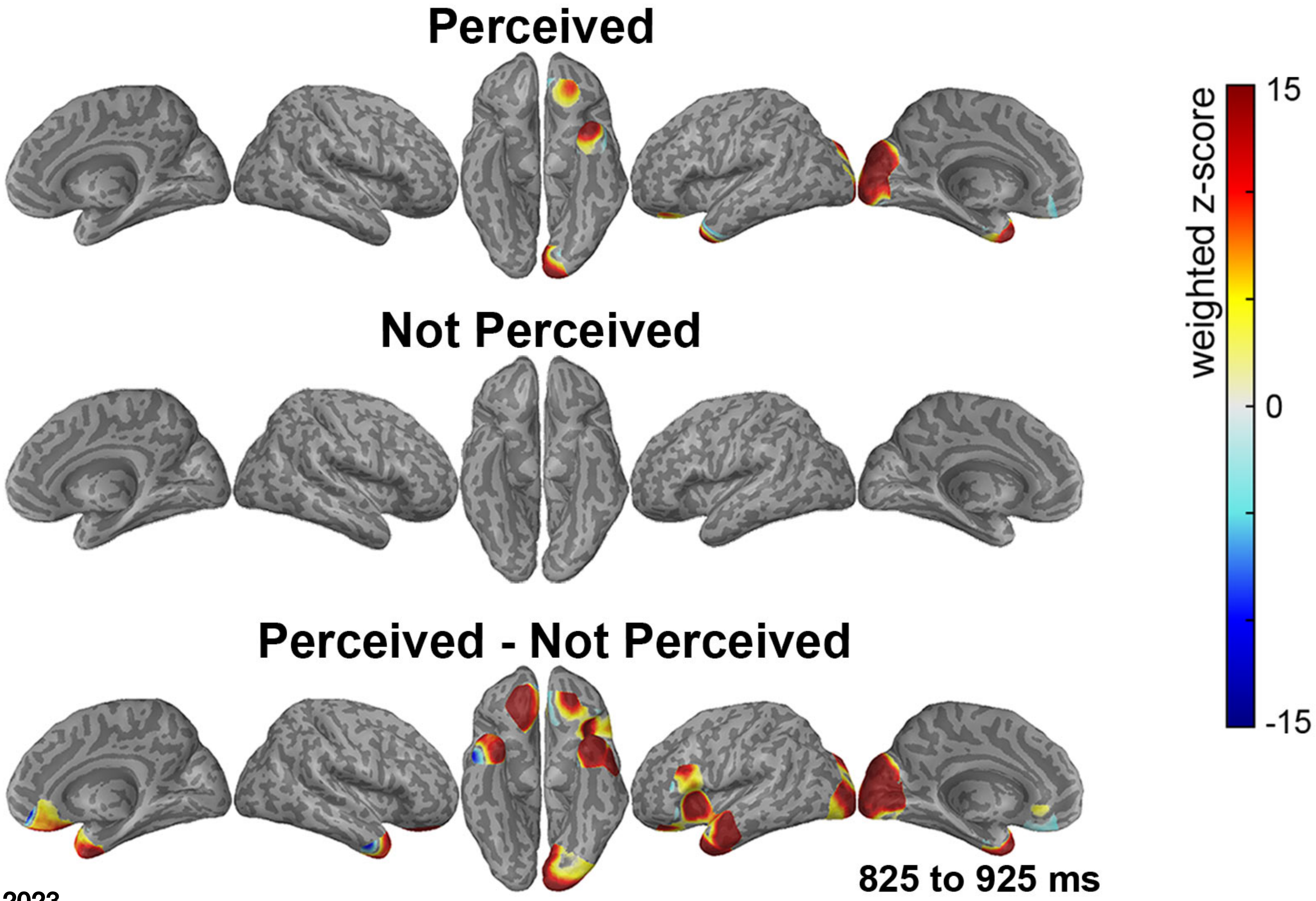


Christison-Lagay et al, 2023

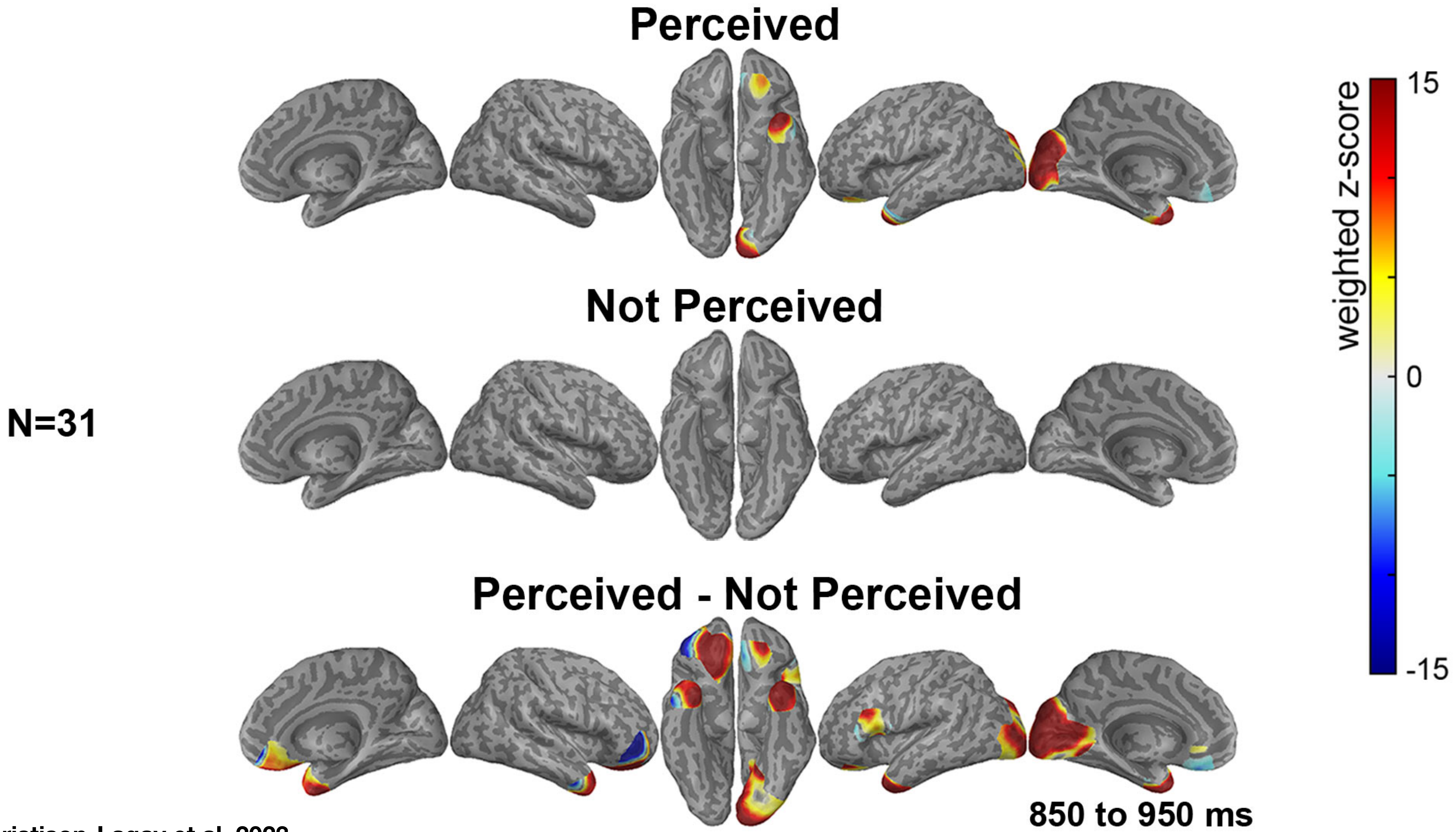


Christison-Lagay et al, 2023

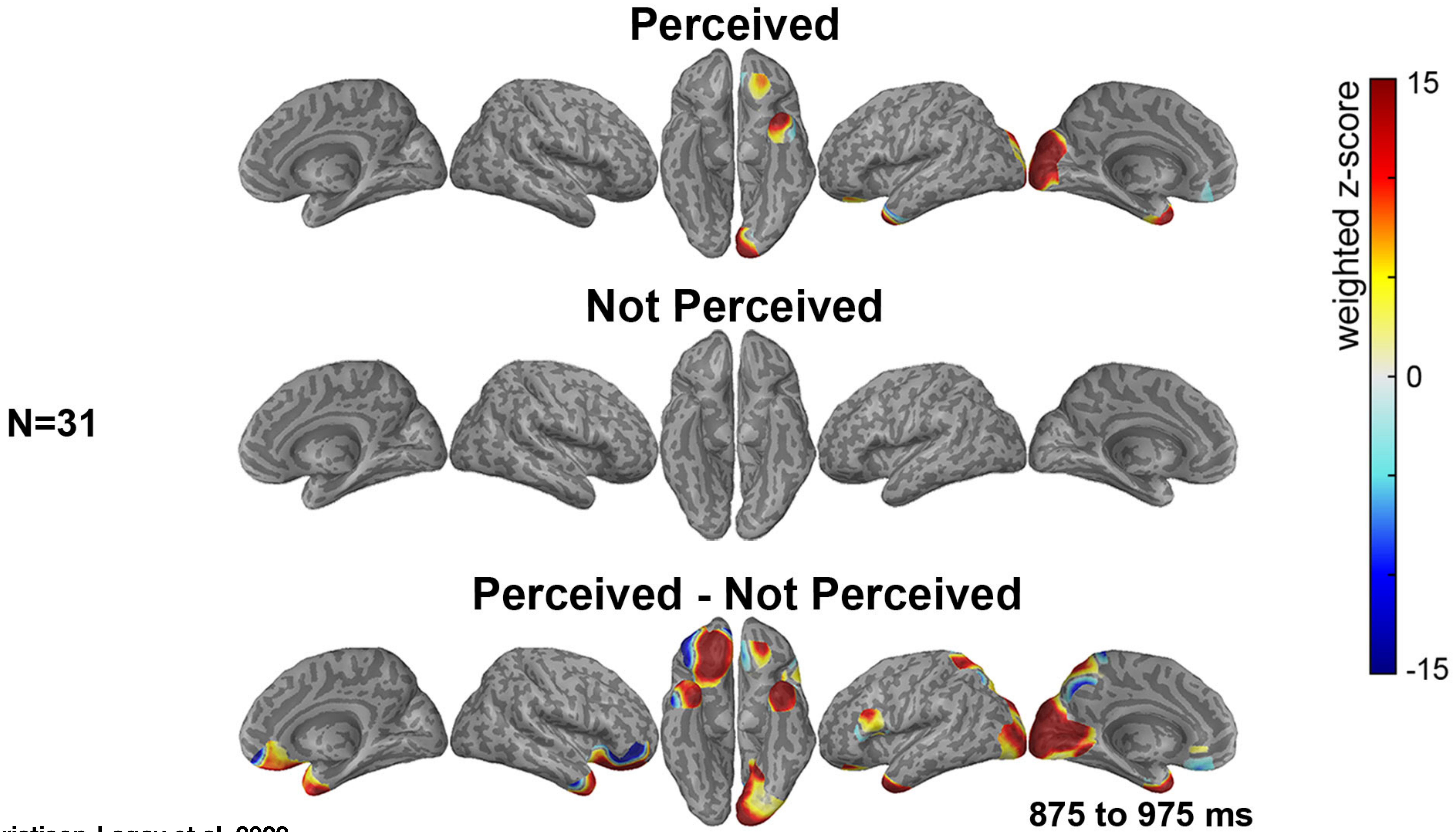


Christison-Lagay et al, 2023

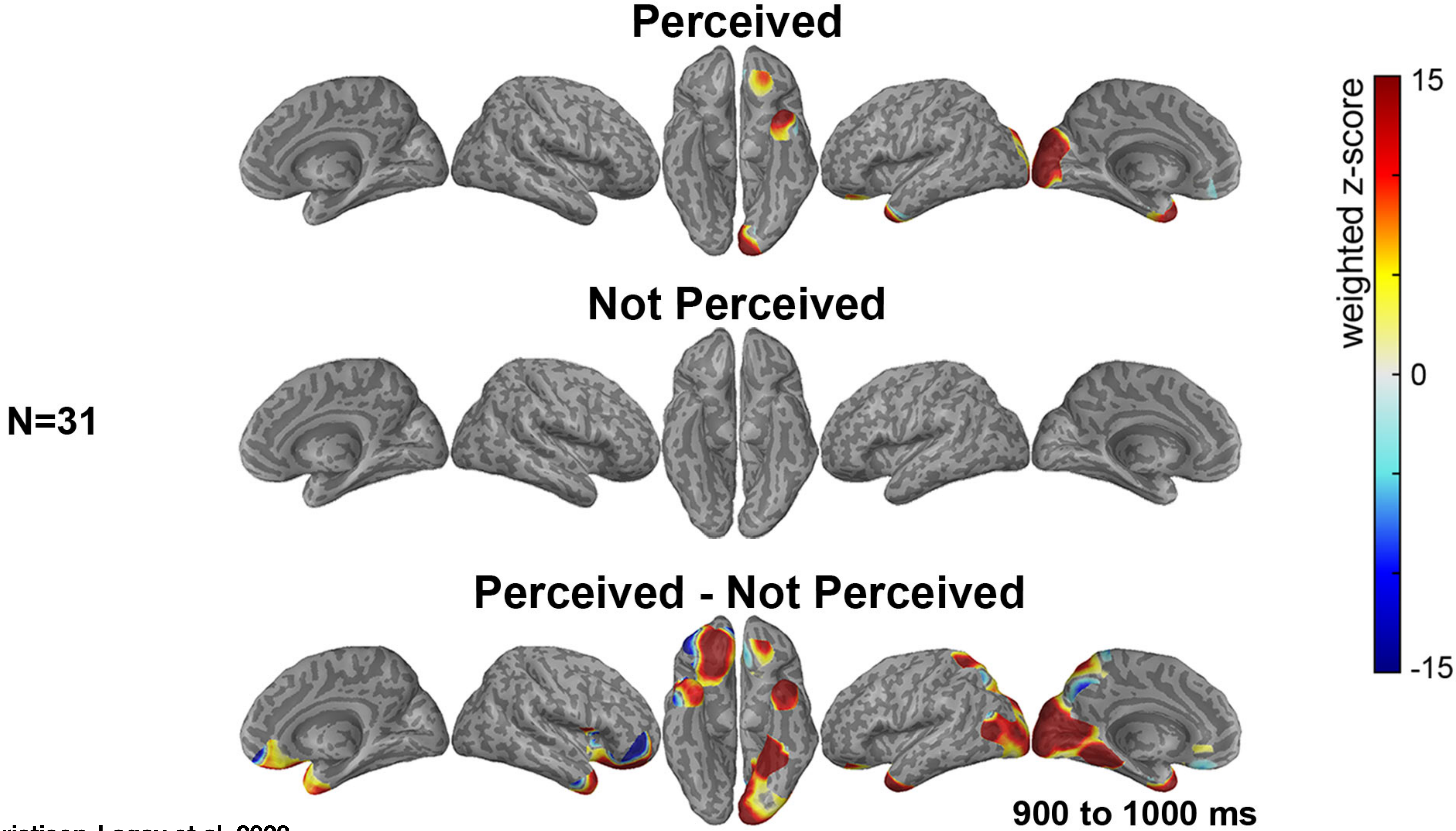

Christison-Lagay et al, 2023

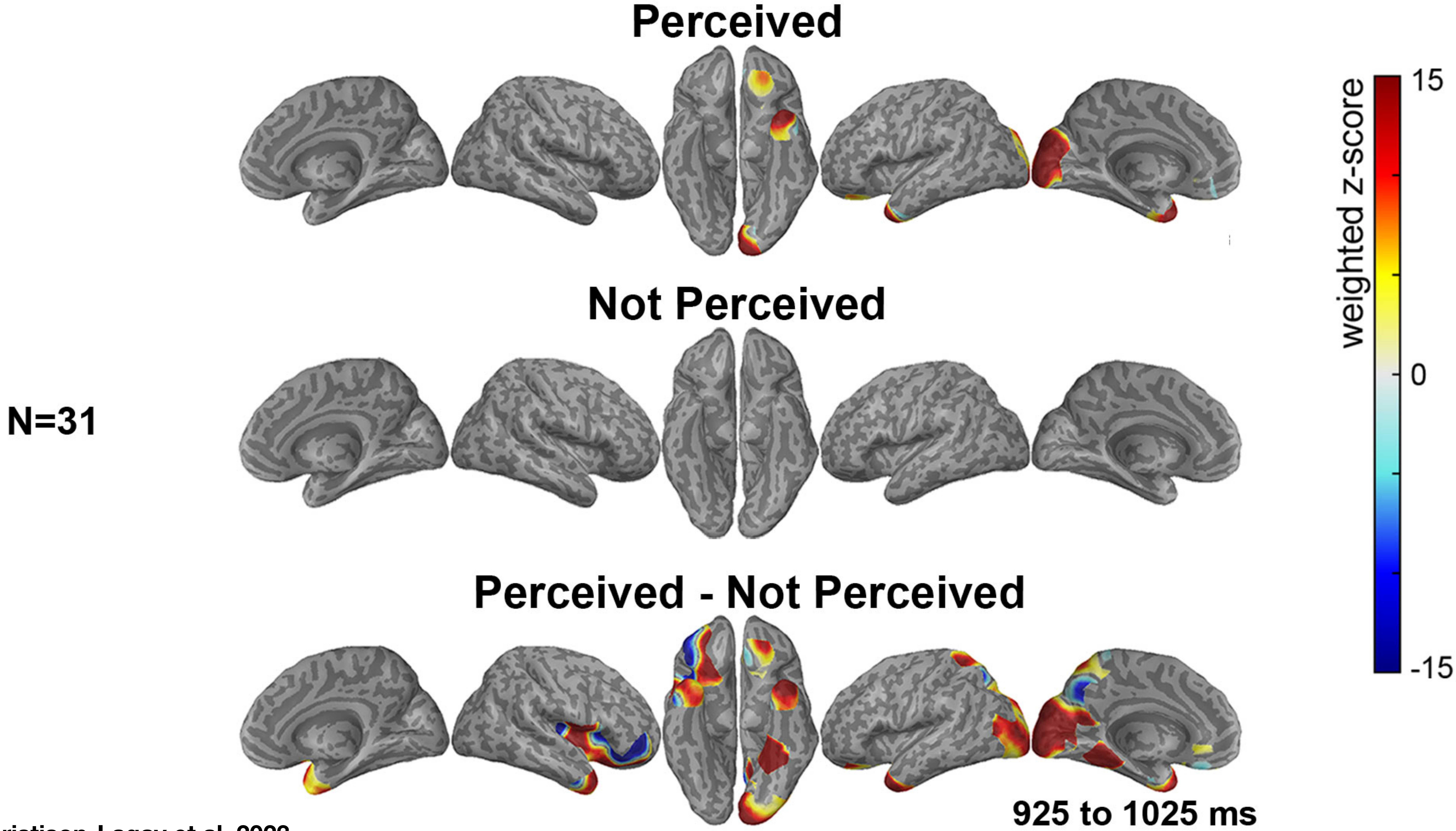


Christison-Lagay et al, 2023

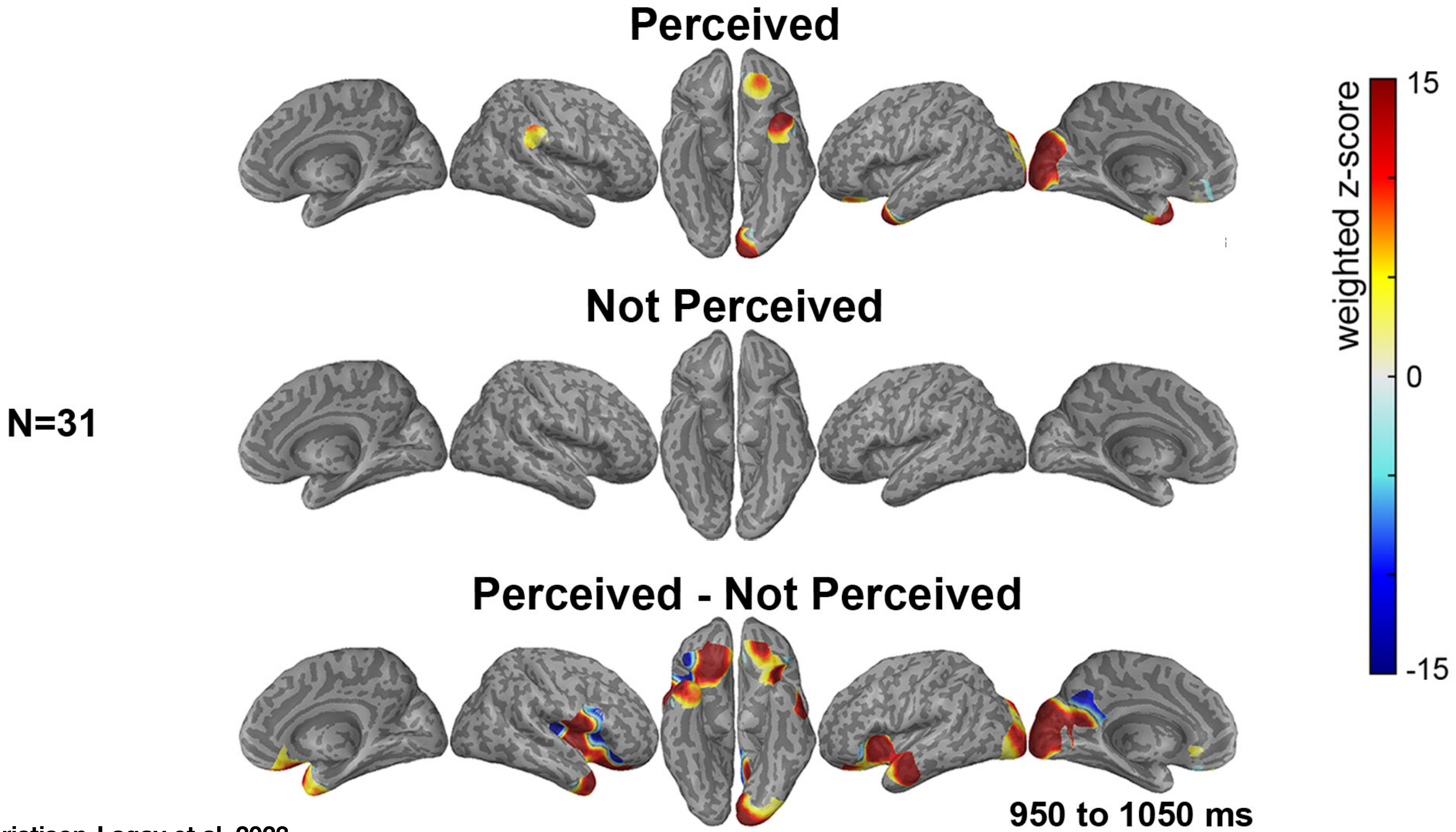


Christison-Lagay et al, 2023

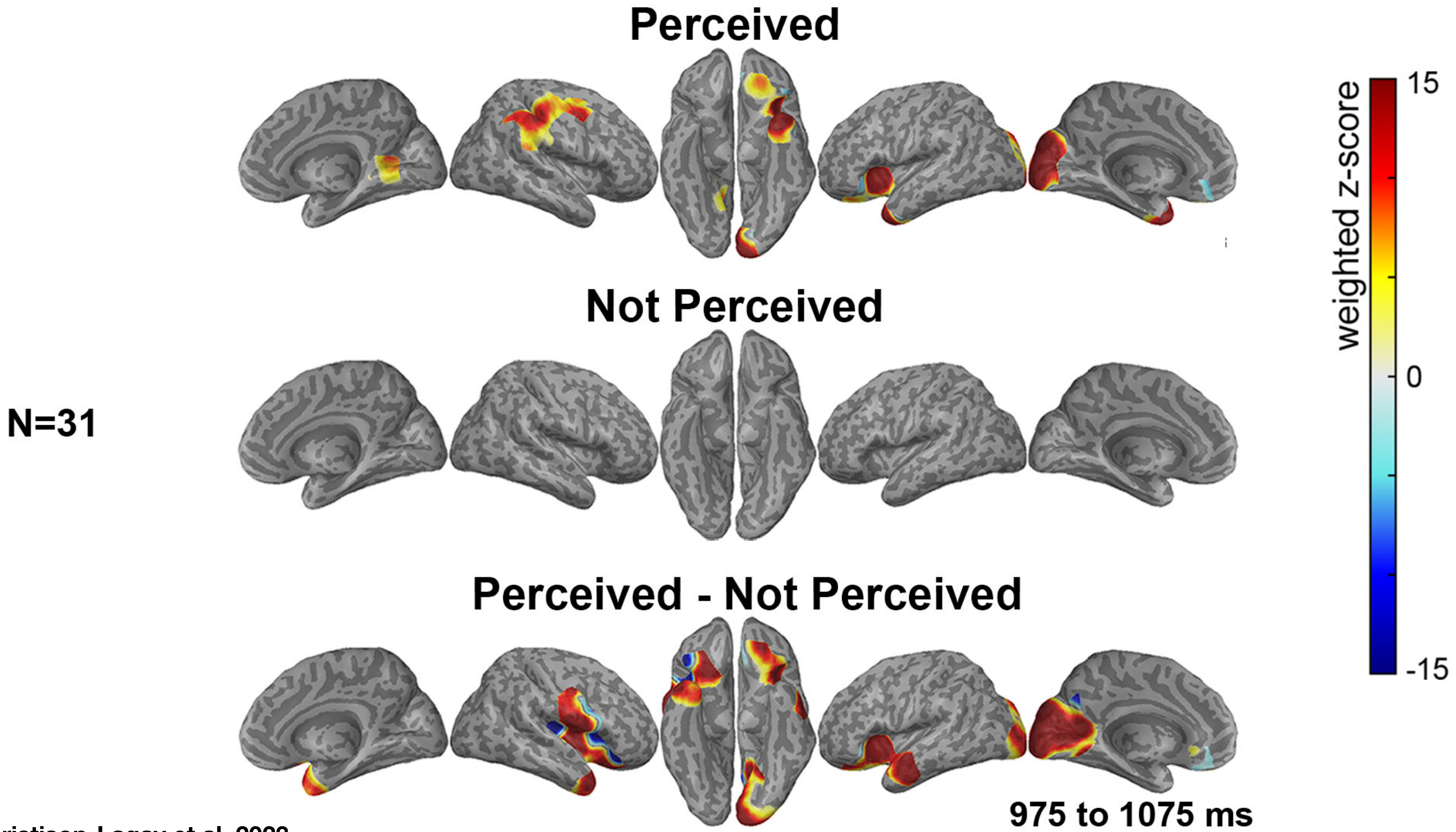

Christison-Lagay et al, 2023

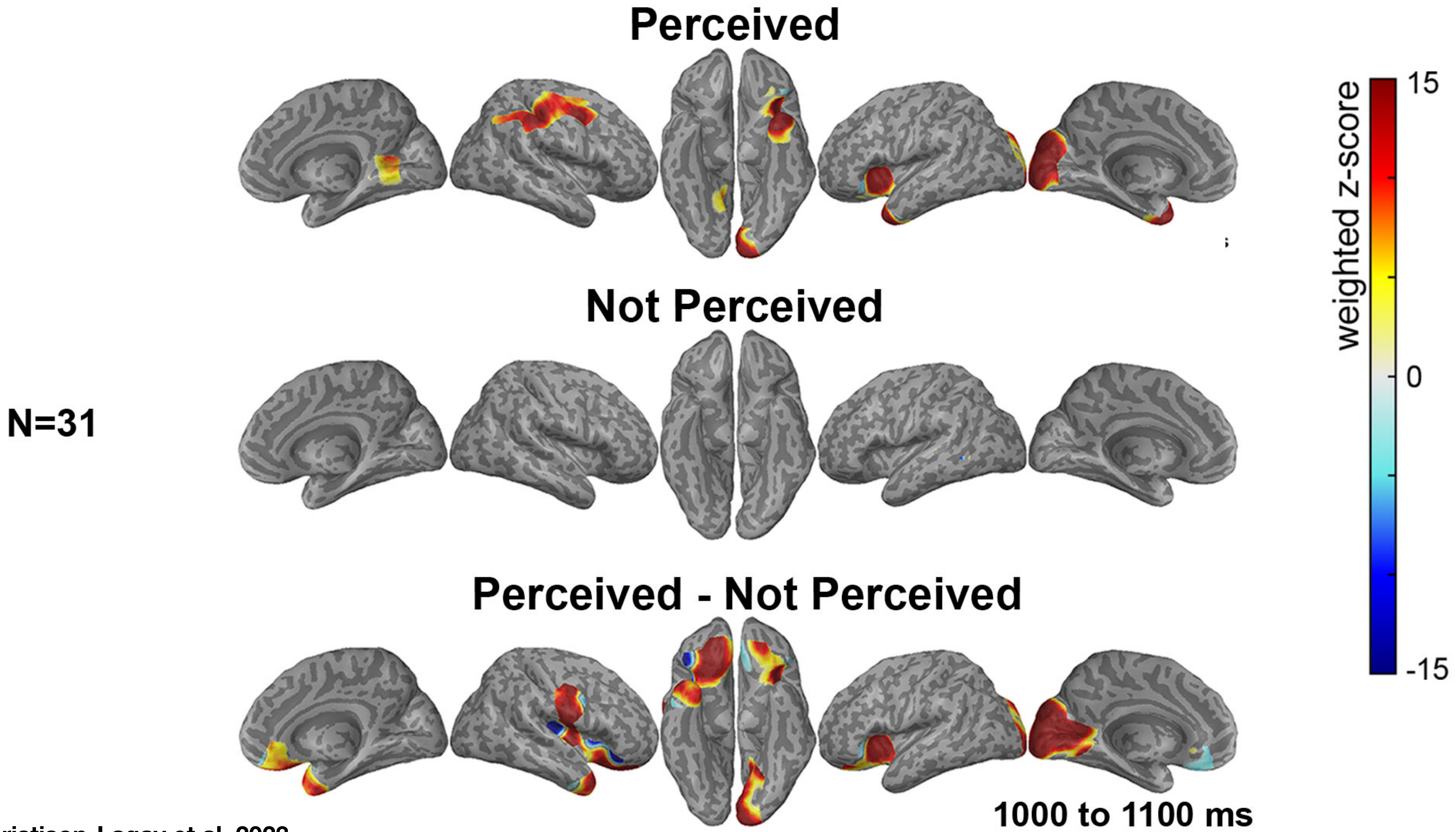


Christison-Lagay et al, 2023

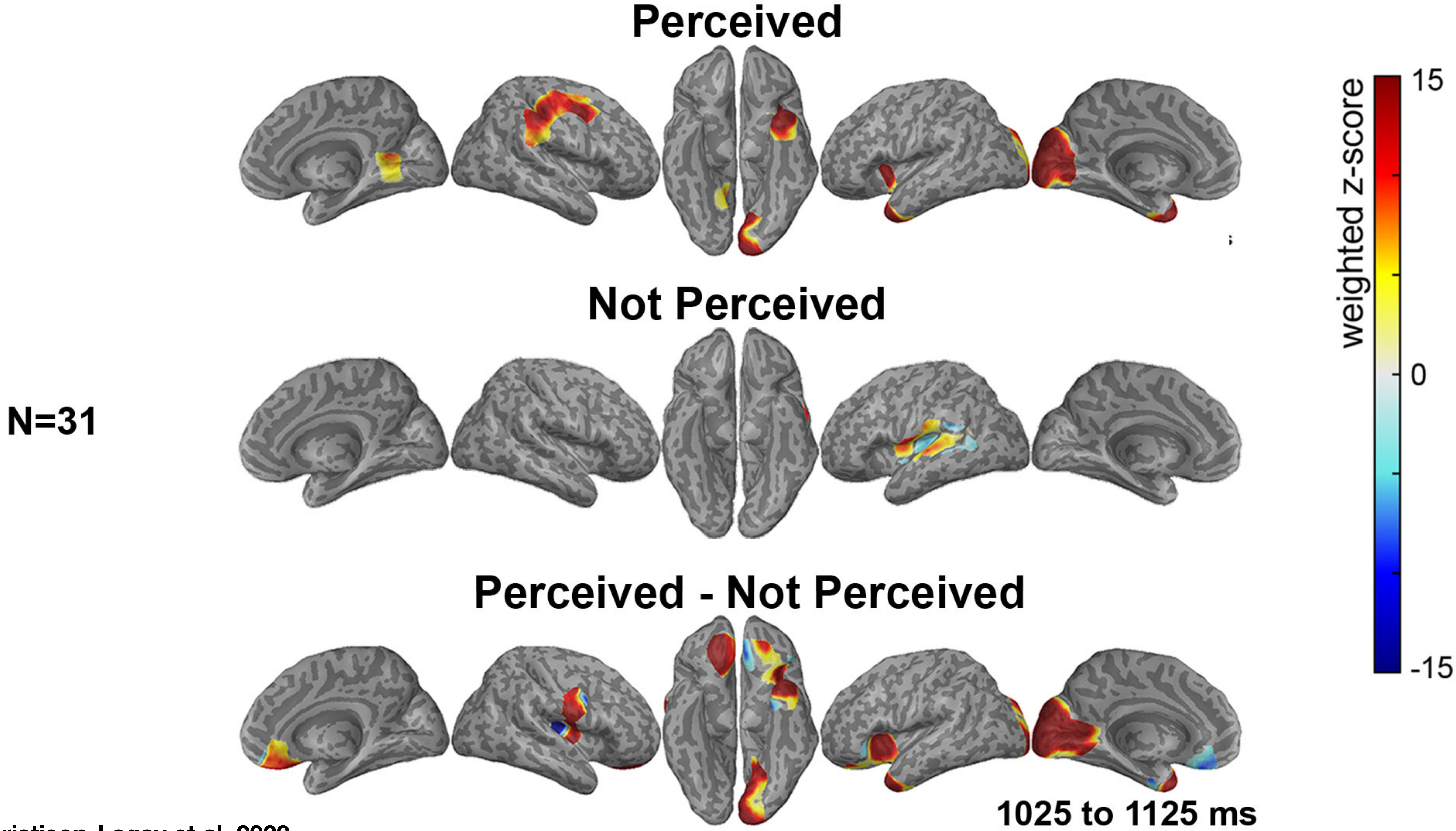


Christison-Lagay et al, 2023

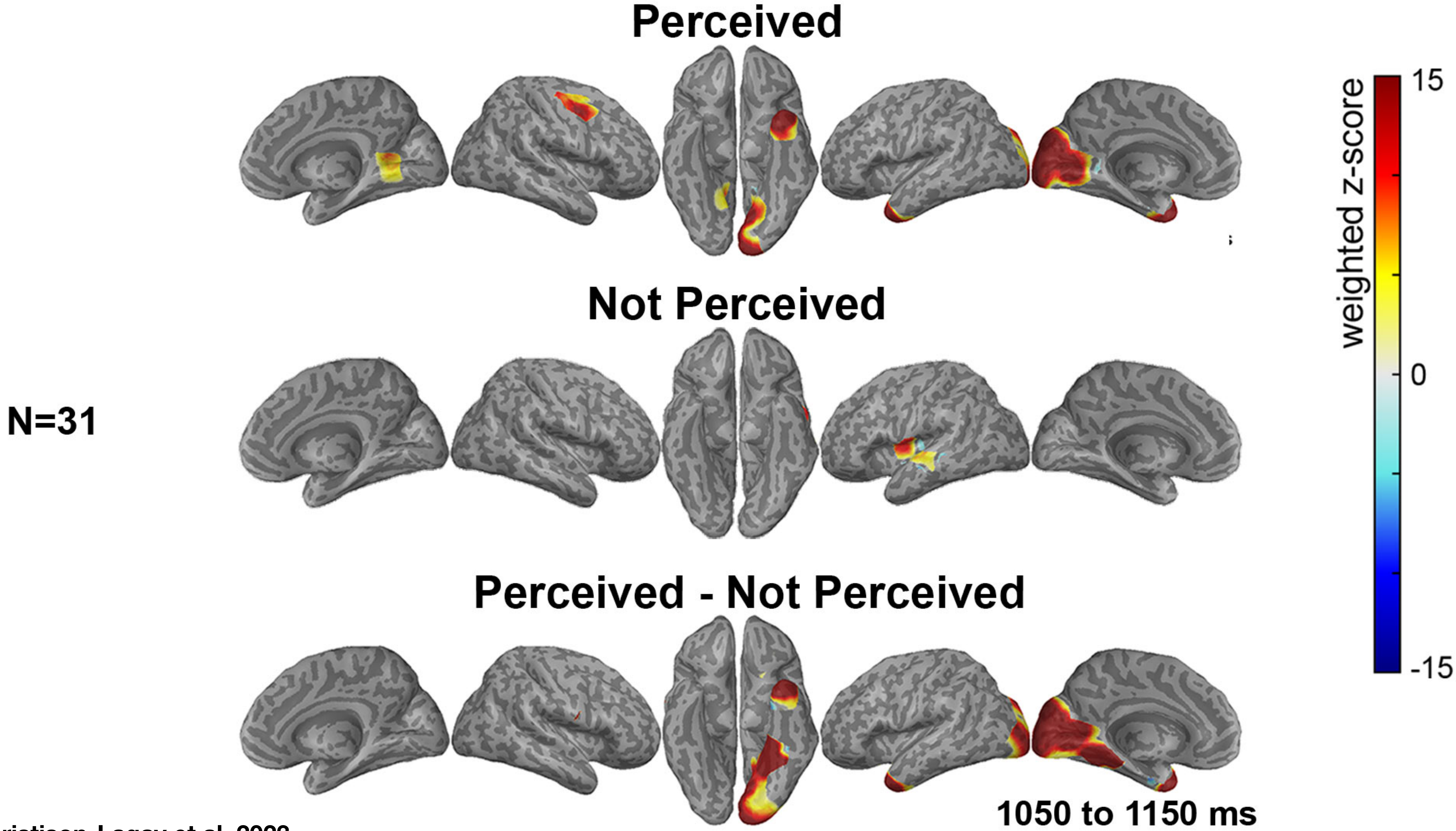


Christison-Lagay et al, 2023

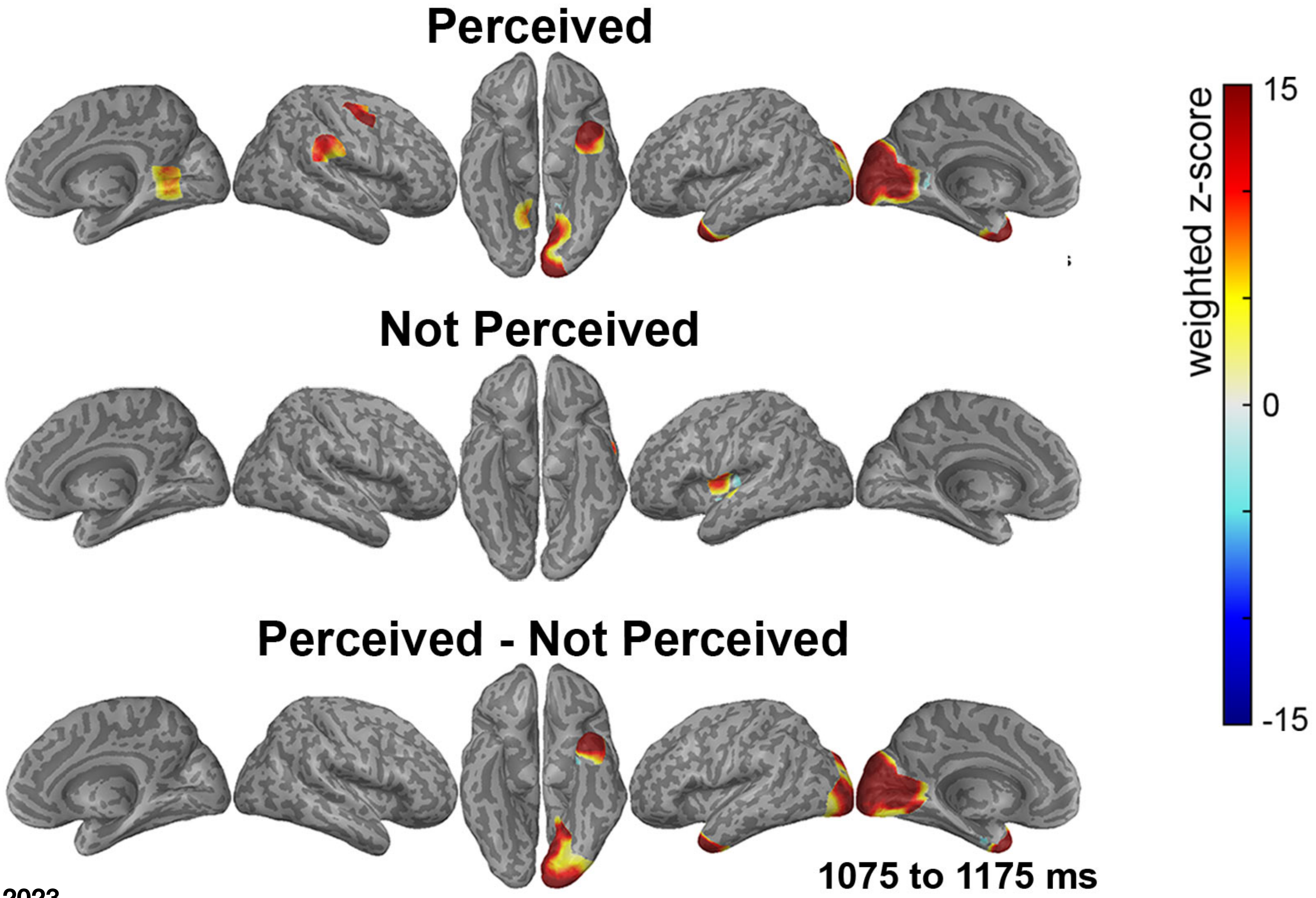


N=31

Christison-Lagay et al, 2023

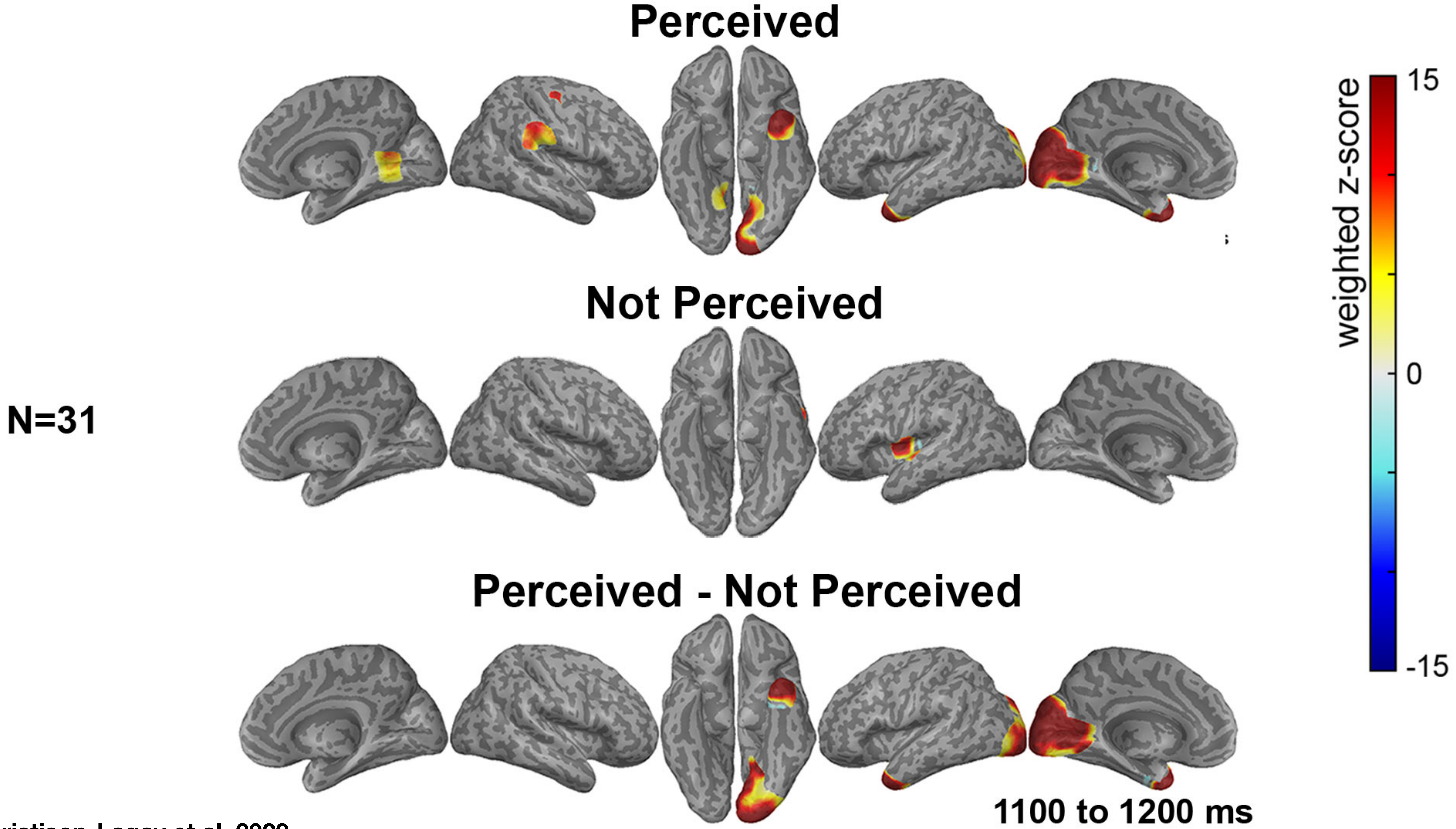

Christison-Lagay et al, 2023

N=31

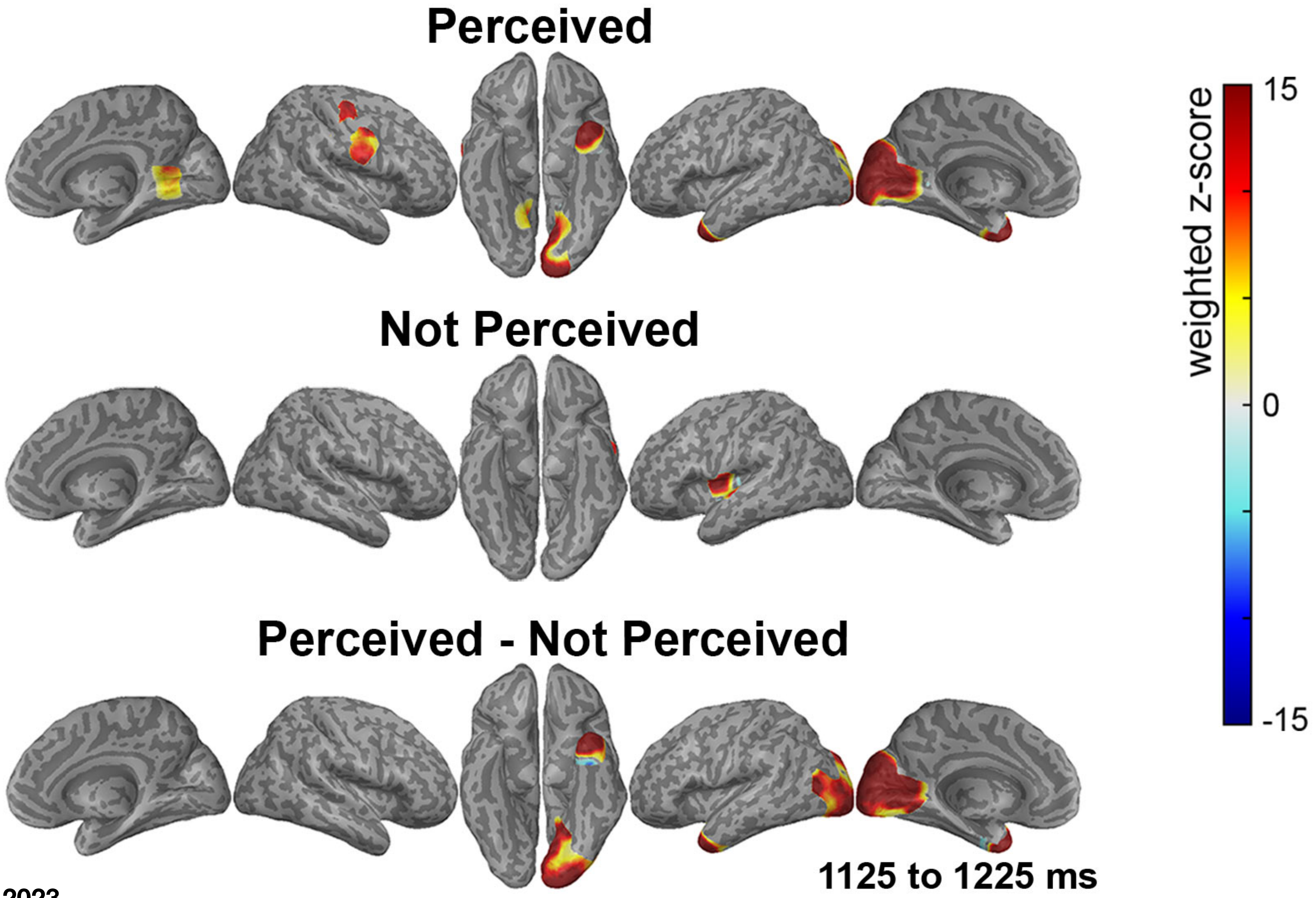


Christison-Lagay et al, 2023

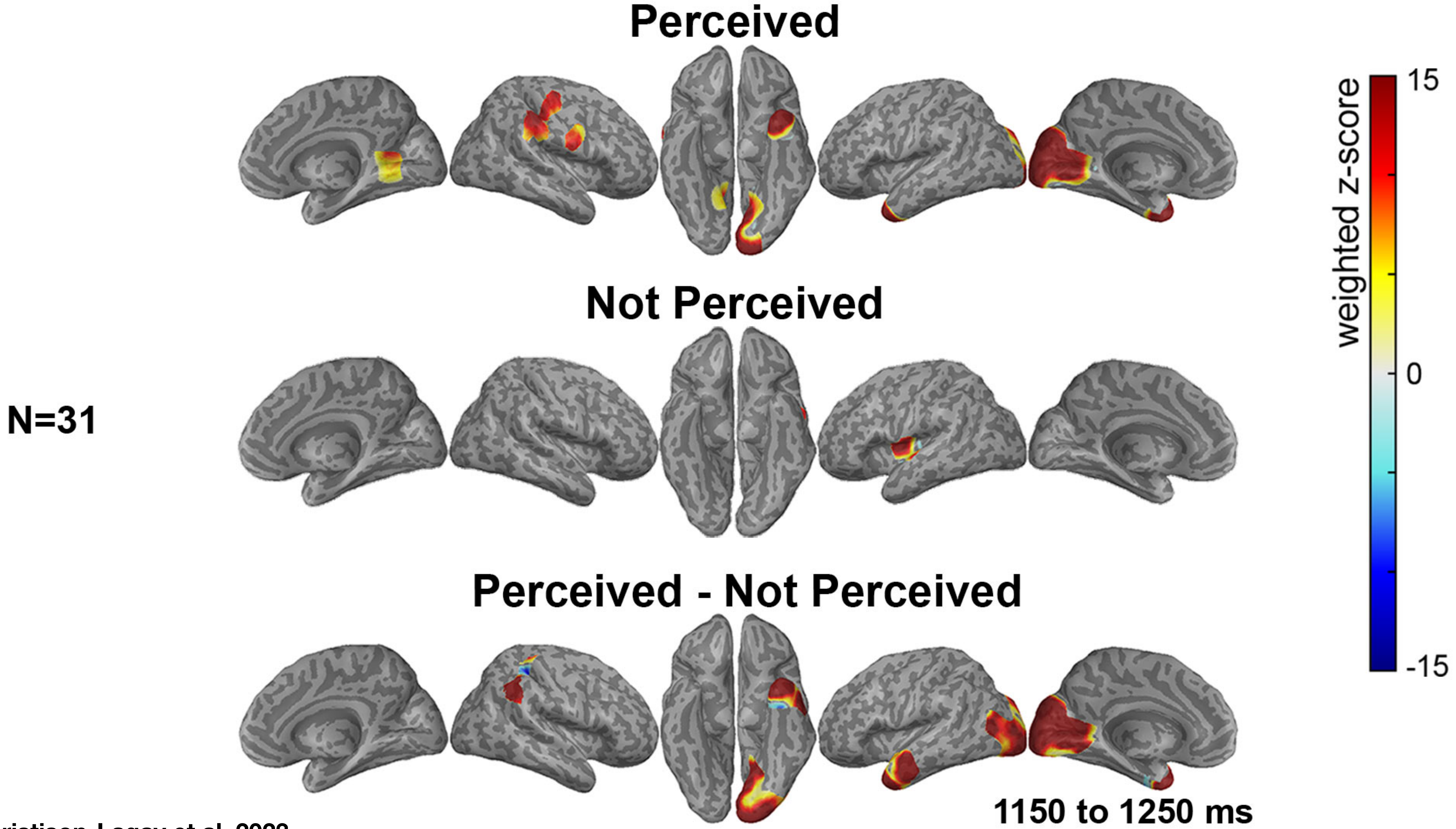


Christison-Lagay et al, 2023

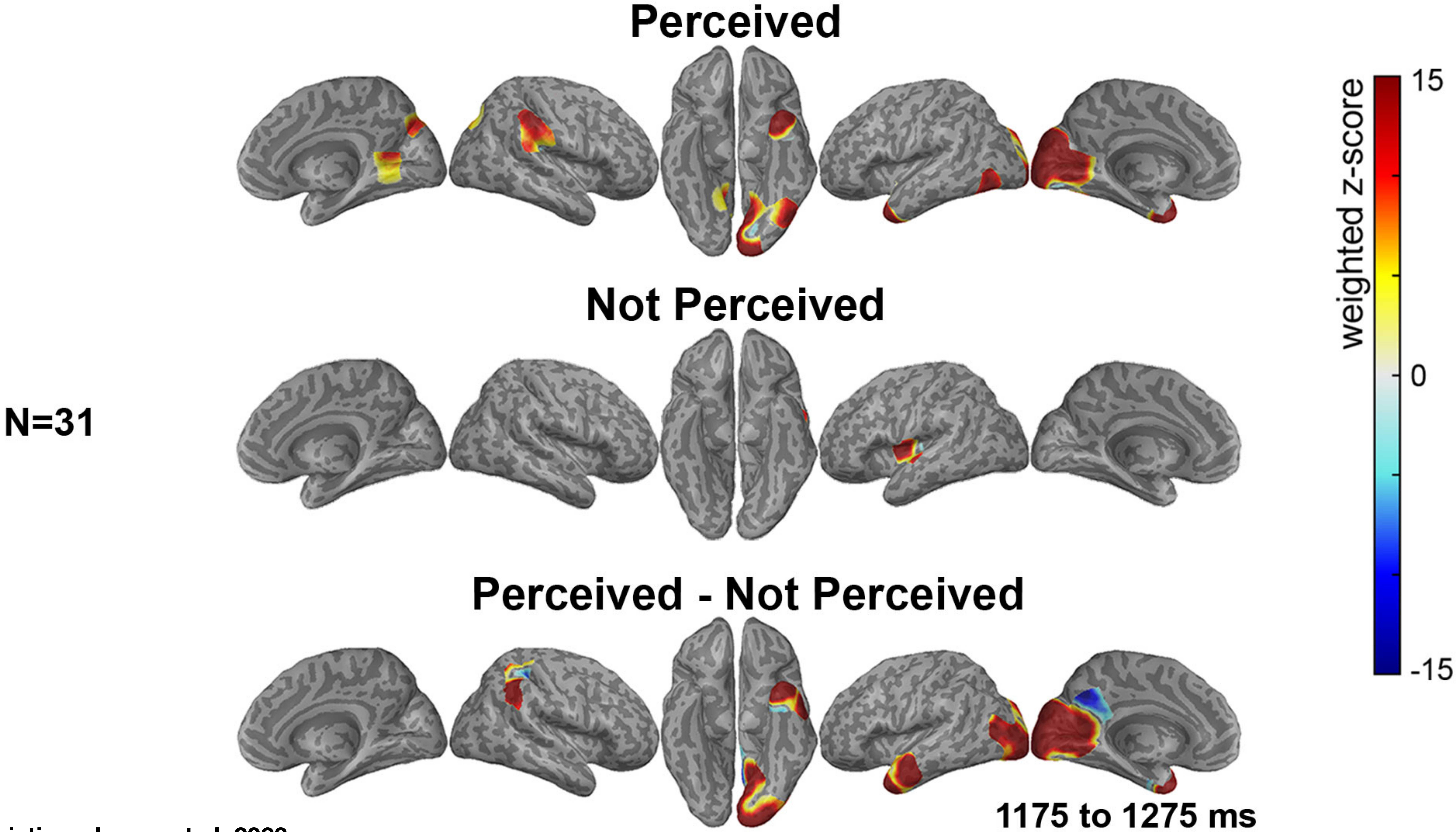


Christison-Lagay et al, 2023

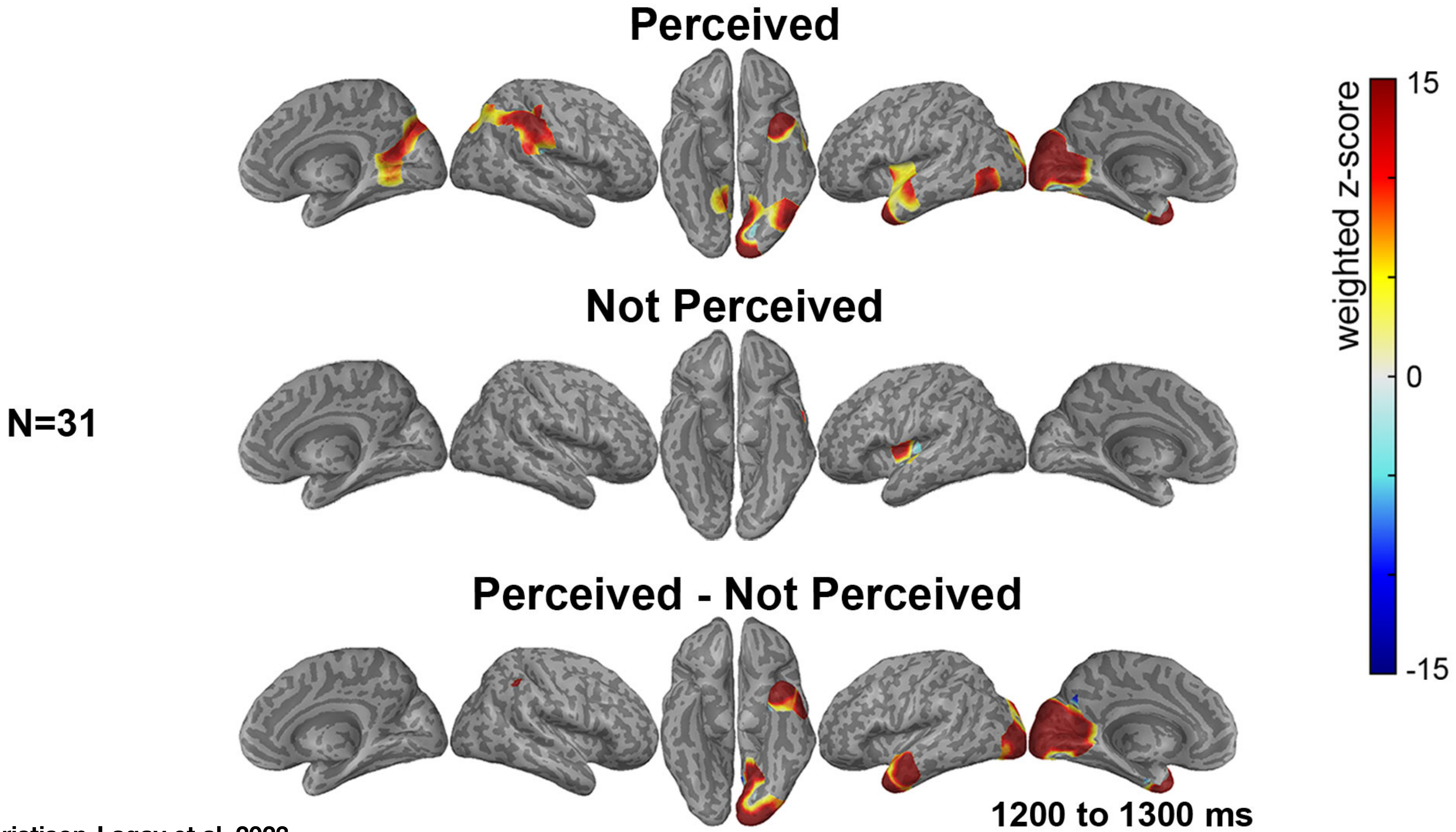


Christison-Lagay et al, 2023

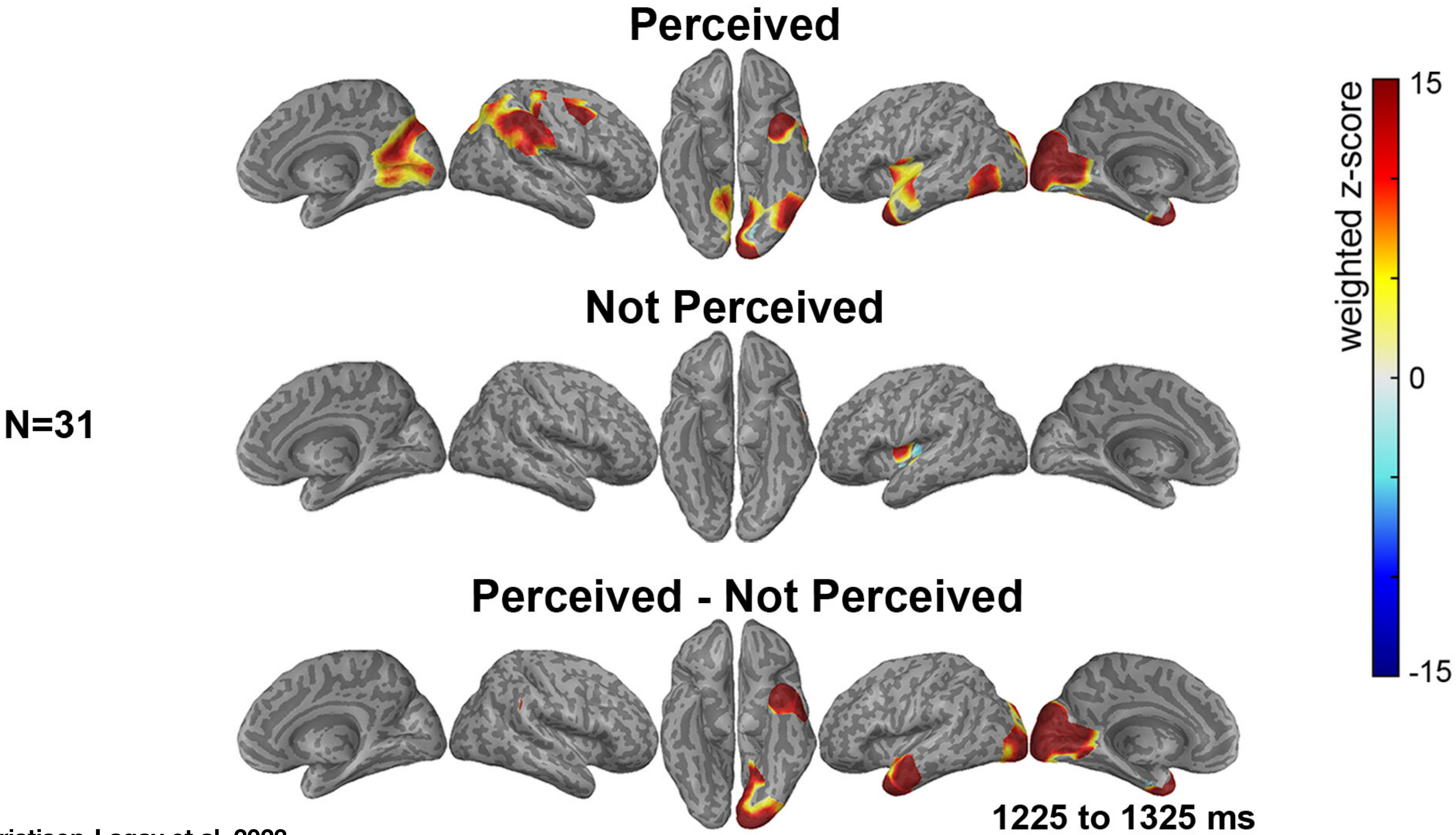


Christison-Lagay et al, 2023

N=31

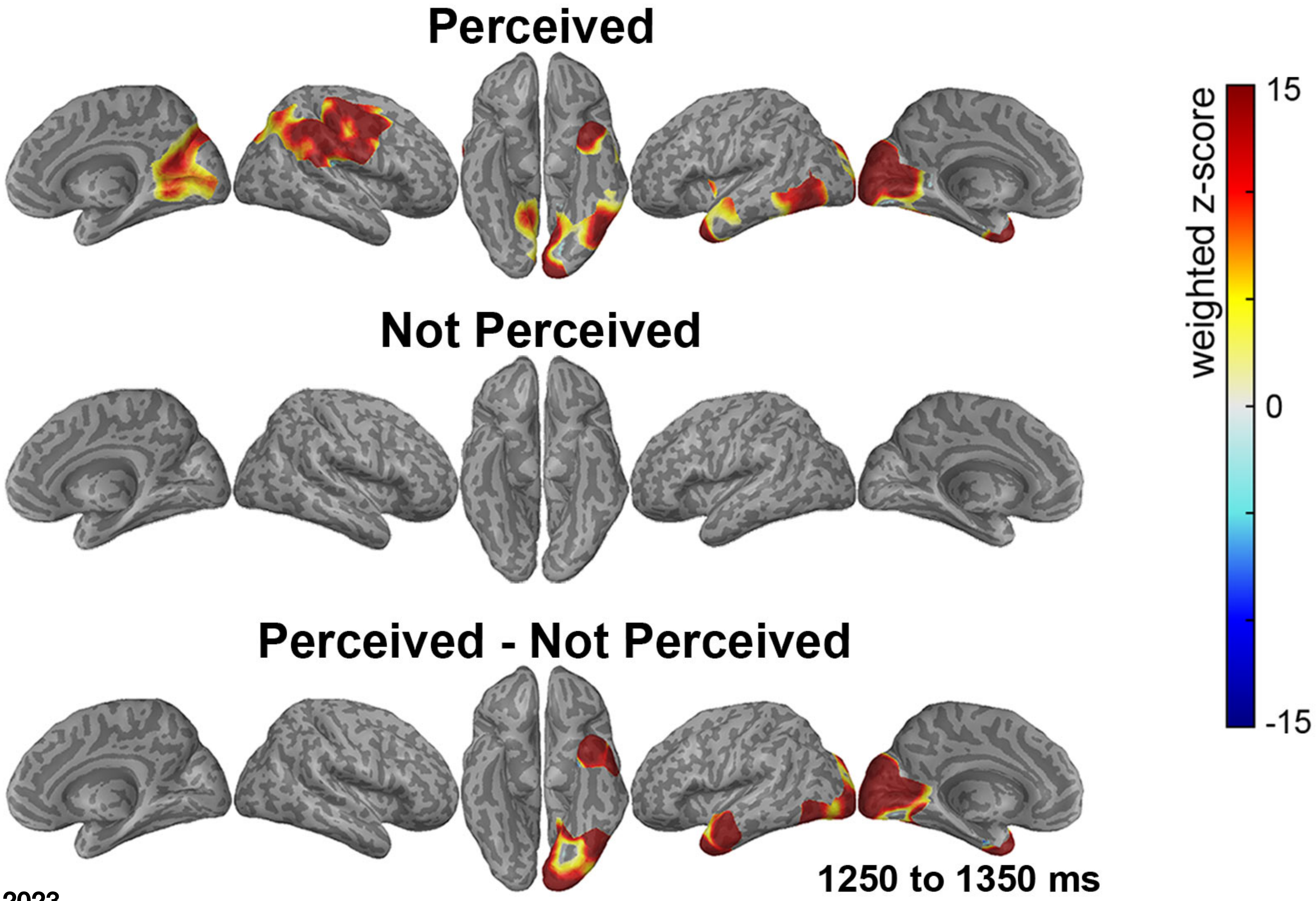


Christison-Lagay et al, 2023

N=31

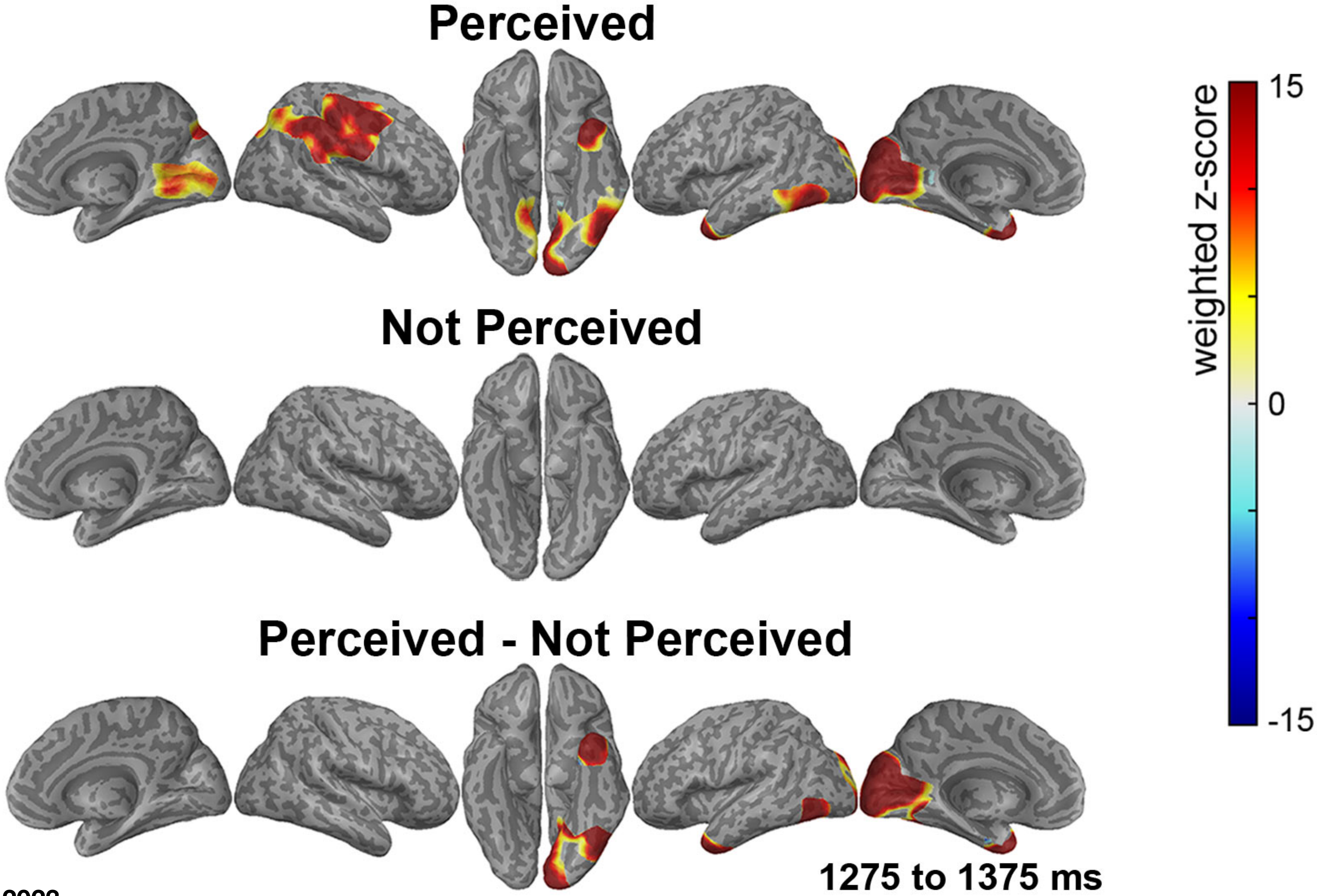


Christison-Lagay et al, 2023

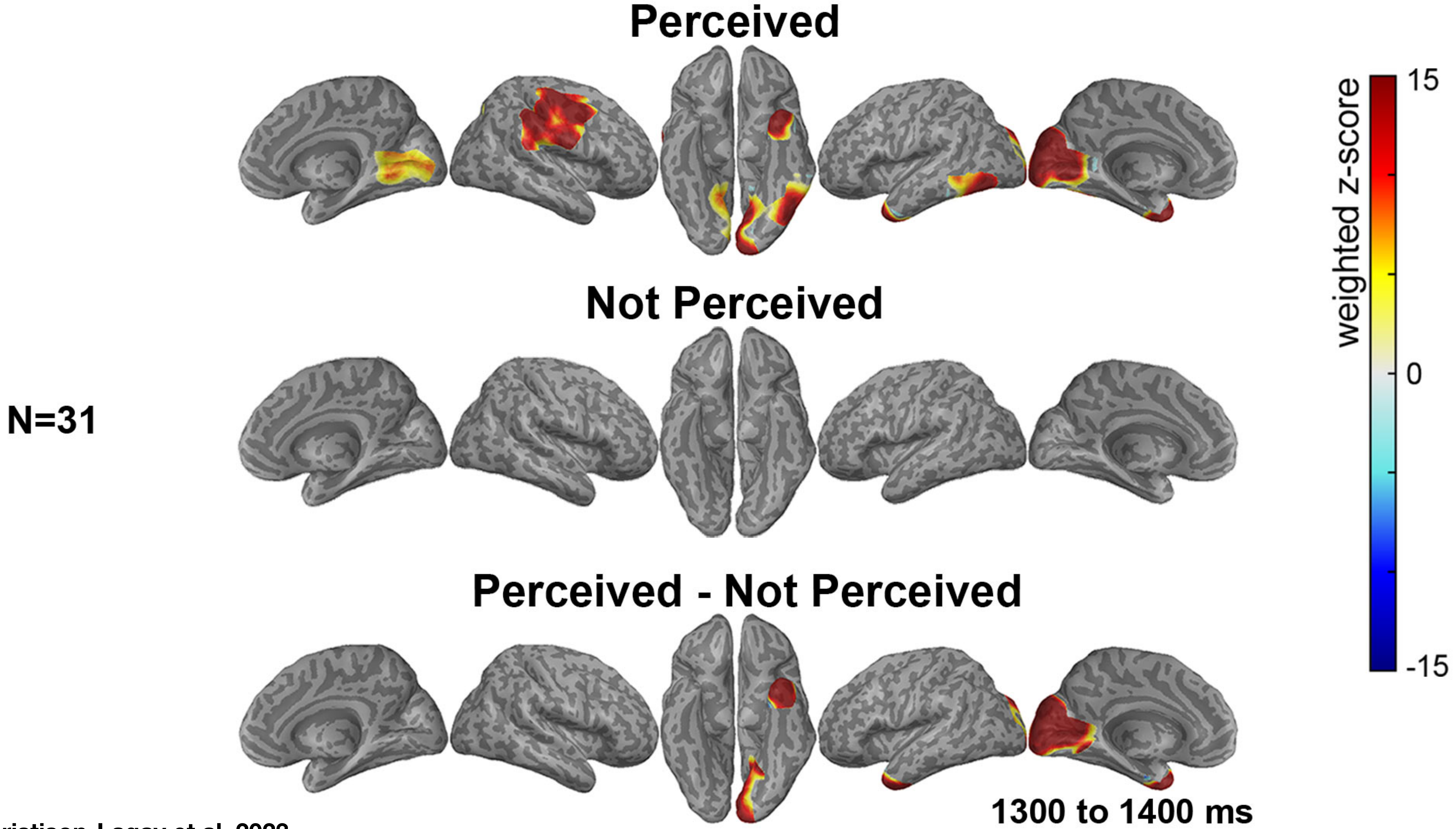


Christison-Lagay et al, 2023

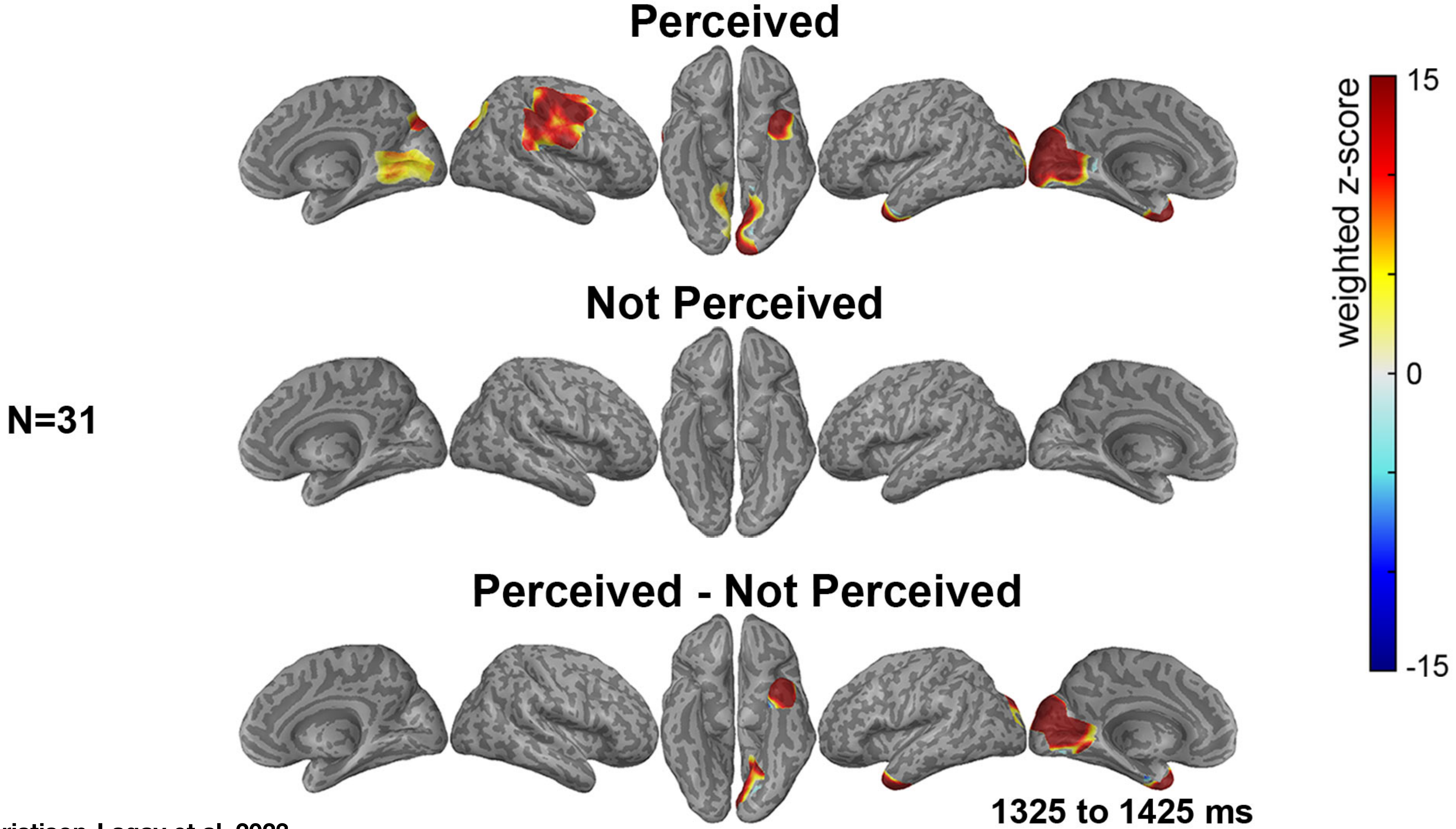


Christison-Lagay et al, 2023

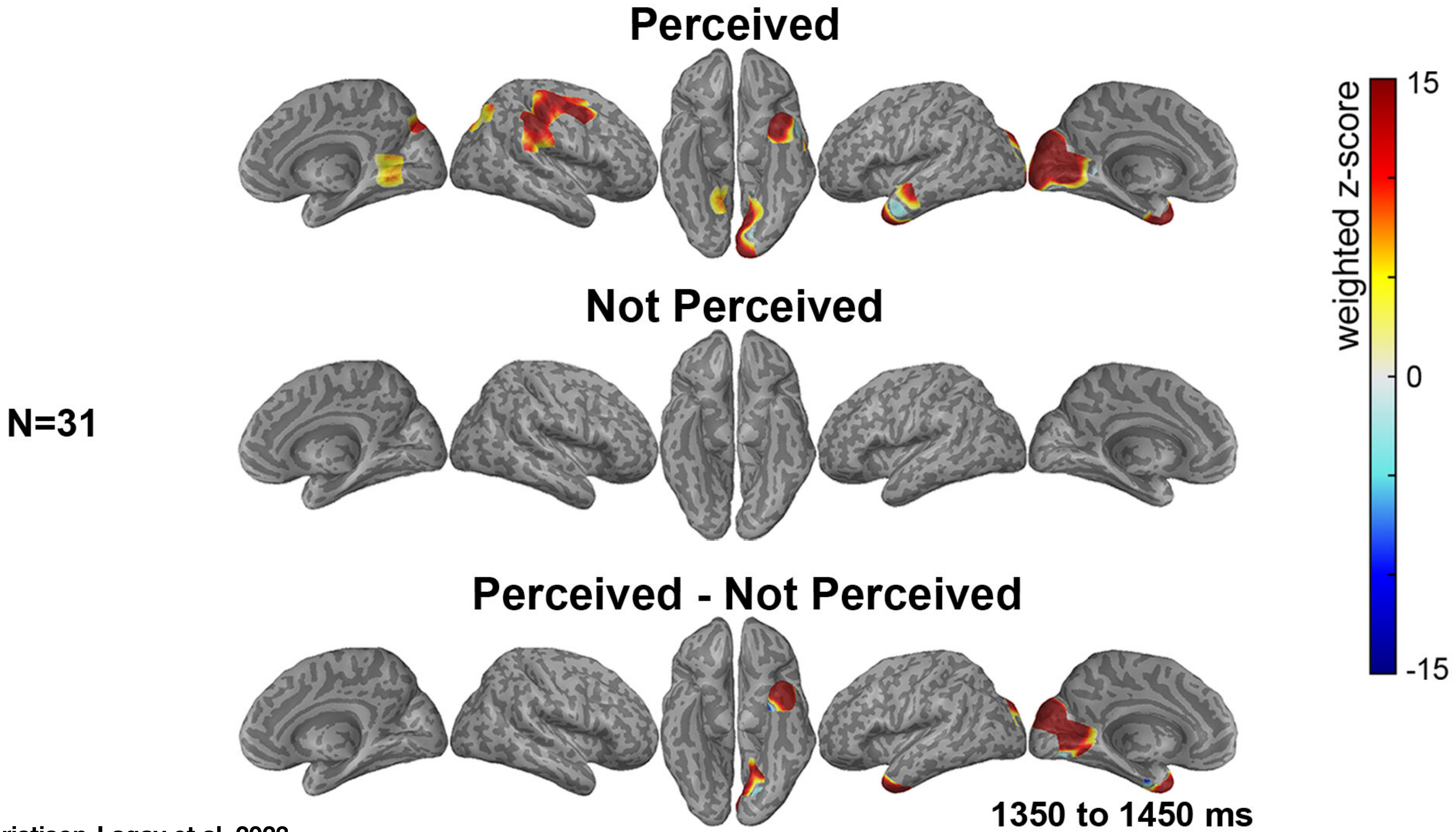


Christison-Lagay et al, 2023

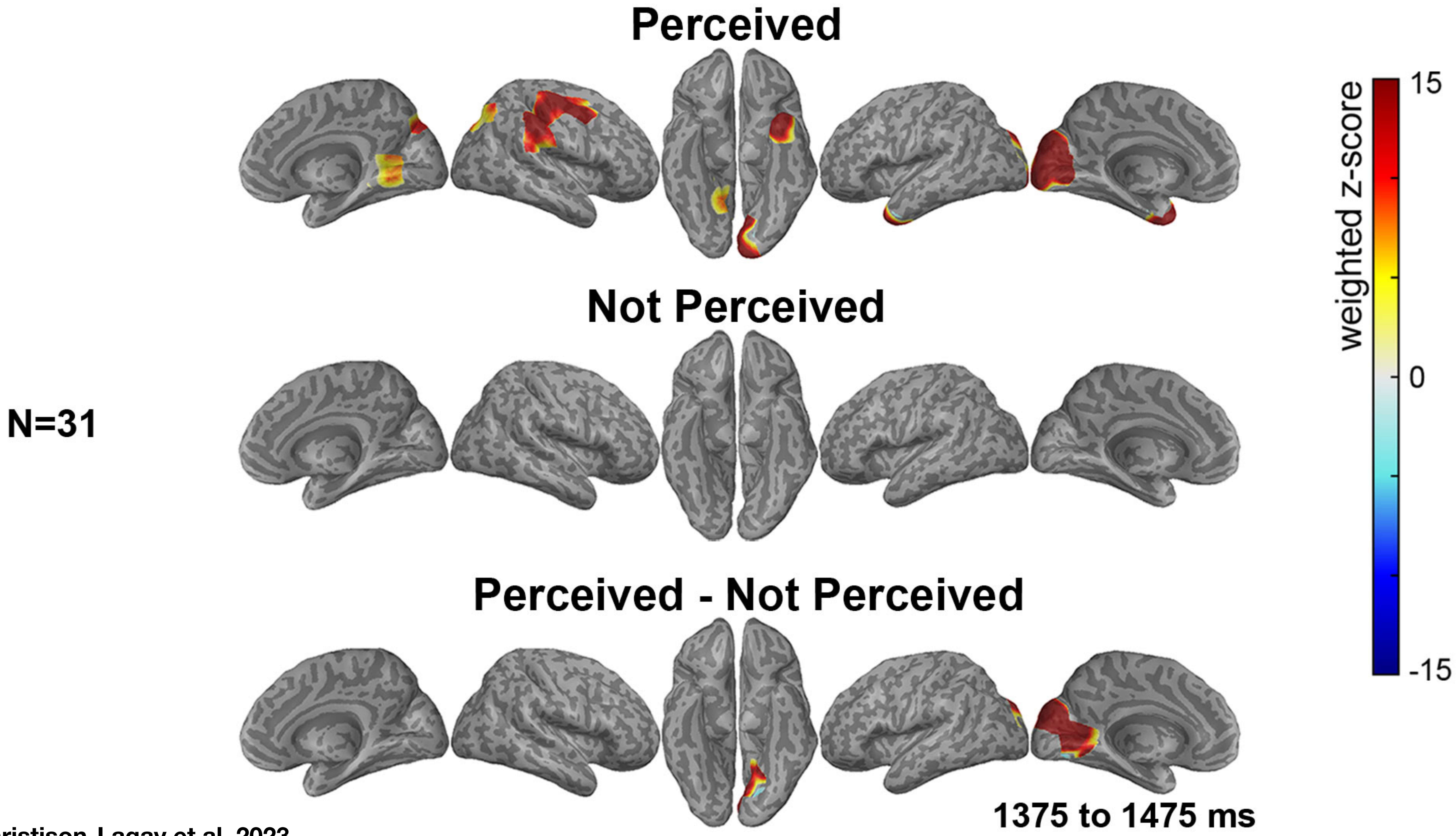


Christison-Lagay et al, 2023

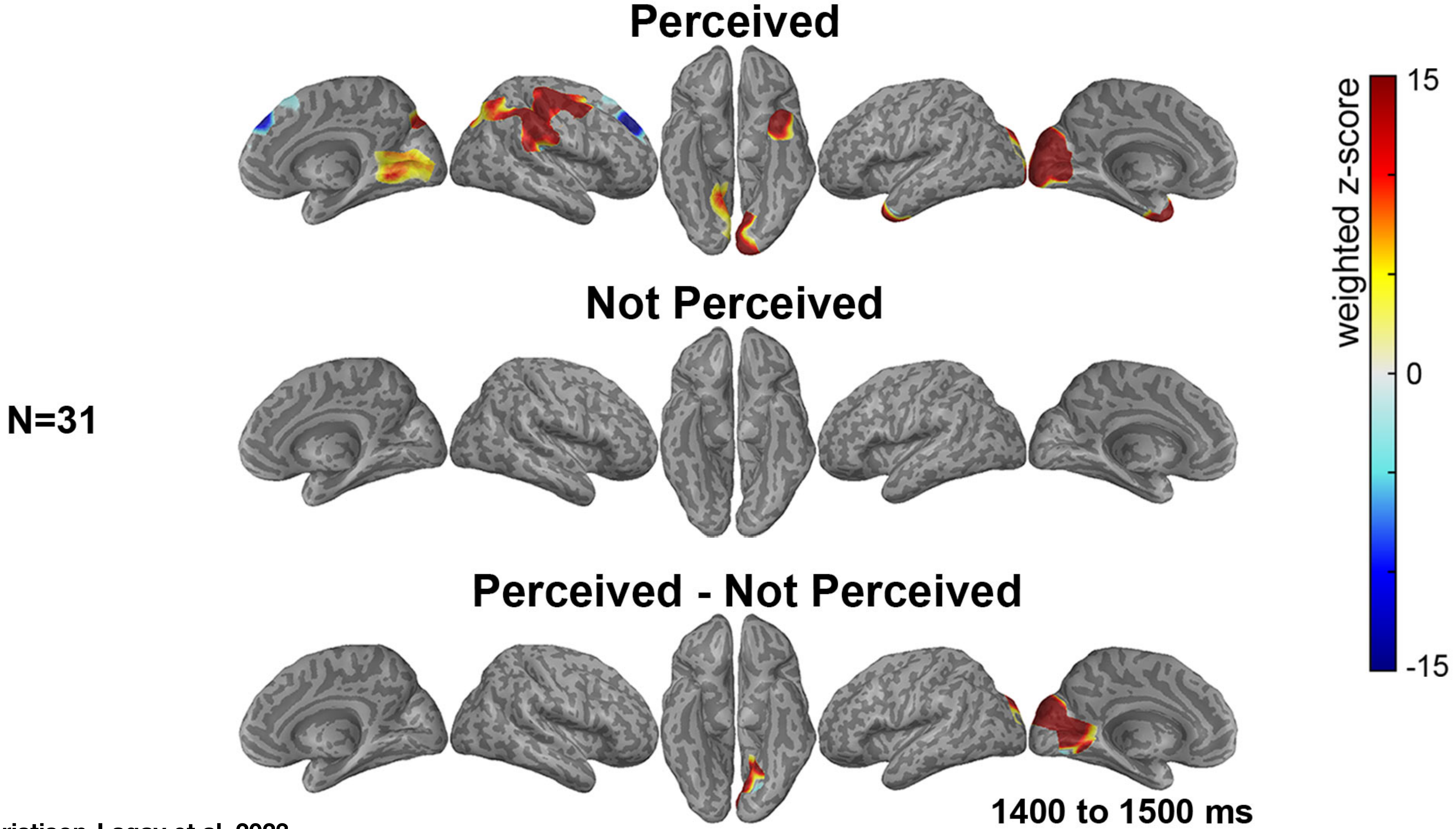


Christison-Lagay et al, 2023
